# Heterogeneous plasticity of amygdala interneurons in associative learning and extinction

**DOI:** 10.1101/2024.09.29.612271

**Authors:** Natalia Favila, Jessica Capece Marsico, Benjamin Escribano, Catarina M. Pacheco, Yael Bitterman, Jan Gründemann, Andreas Lüthi, Sabine Krabbe

**Affiliations:** German Center for Neurodegenerative Diseases (DZNE), Bonn, Germany; Medical Faculty, University of Bonn, Bonn, Germany; Friedrich Miescher Institute for Biomedical Research, Basel, Switzerland; The Hebrew University of Jerusalem, Jerusalem, Israel; University of Basel, Basel, Switzerland

## Abstract

Neural circuits undergo experience-dependent plasticity to form long-lasting memories. Excitatory projection neurons are considered to be the primary neuronal substrate for memory acquisition and storage. However, inhibitory interneurons control the activity of projection neurons in a in a spatially and temporally precise manner, yet their contribution to memory acquisition, storage and expression remains poorly understood. Here, we employ a miniature microscope imaging approach to monitor the activity of large amygdala interneuron populations in freely moving mice during fear learning and extinction at the single cell level. We find that amygdala interneurons display mixed-selectivity and show complex plastic responses at both the ensemble and single neuron level across the acquisition, expression and extinction of aversive memories. In contrast to bidirectional single cell plasticity across distinct fear states, learning-induced changes at the population level occur transiently during conditioning and do not consolidate across days. Examining molecular interneuron subpopulations revealed that disinhibitory vasoactive intestinal peptide (VIP) expressing cells are predominantly activated by high fear states. In contrast, somatostatin (SST) interneurons display a preference for safety cues and thereby suppress excitatory neuron responsiveness. However, responses of individual neurons within the SST and VIP populations are non-uniform, indicating the presence of functional subtypes within classical molecularly-defined interneuron populations. Taken together, we identify complex neuronal plasticity within amygdala interneuron ensembles that goes beyond a passive processing function, suggesting a critical role of inhibitory microcircuit elements for memory selectivity and stability.

## INTRODUCTION

Associative learning enables an organism to link environmental stimuli with their behavioural relevance. This is particularly important under conditions of immediate threat, such as in fear learning. One of the key brain regions regulating the acquisition, expression and extinction of conditioned fear behaviour is the amygdala, a highly conserved temporal lobe structure consisting of distinct subnuclei. Traditionally, excitatory projection neurons (PNs) of the cortex-like basolateral amygdala (BLA) were regarded as the main site of plasticity during memory formation^1^. Yet, learning processes are strongly influenced by dynamic shifts in the balance between excitatory and inhibitory components within neuronal circuits. For example, fear learning reduces extracellular GABA levels^2^ and modifies the expression of the GABA-synthesising enzyme GAD67, GABA receptors and their clustering protein gephyrin in the BLA^3–5^. Furthermore, BLA inhibitory circuits show diverse forms of synaptic plasticity *ex vivo*^6–9^, which might be involved in both fear and extinction memory^10^. However, how interneuron activity changes during memory formation *in vivo* remains largely unexplored.

BLA interneurons are highly diverse and form intricate microcircuits. Distinct interneuron subpopulations can be distinguished based on marker gene expression, morphology, pre- and postsynaptic connectivity or functional properties, all of which are considered to be highly correlated with each other^11–13^. Even though they only represent about 15-20% of the neuronal population in the BLA, interneurons control PN activity in a spatially and temporally precise manner due to their cell type- and cellular compartment-specific postsynaptic targeting. In consequence, selective changes in the activity patterns associated with memory formation of distinct inhibitory subpopulations can have diverse effects on PNs and ultimately amygdala output.

Recent studies started to delineate the contributions of interneuron subpopulations to fear learning and extinction. For example, somatostatin (SST) positive interneurons preferentially target the distal dendrites of BLA PNs and are thus ideally positioned to regulate synaptic inputs from thalamic and cortical sources^14^. Suppression of their activity during conditioning has been shown to enhance learning^14,15^. Furthermore, SSTs play a role in context-dependent fear suppression^16,17^. In contrast, interneurons expressing vasoactive intestinal peptide (VIP) contact other interneuron subpopulations such as SST and parvalbumin (PV) positive cells^18,19^. When activated by aversive events during associative fear conditioning, VIP Interneurons have disinhibitory effects on BLA PNs and can thereby enable excitatory plasticity^18^. In addition, *in vitro* studies have demonstrated cellular plasticity of distinct interneuron subpopulations upon fear and extinction learning^20–22^, highlighting that inhibitory cells undergo experience-dependent plastic changes themselves. However, only few studies recorded the activity of individual neurons during fear learning and extinction. Recent work demonstrated learning-associated plasticity of distinct molecular interneuron subpopulations at the population level using fibre-photometry recodings^23^. Yet, even within the well-defined paradigms of associative fear learning and extinction, individual interneuron subpopulations display high heterogeneity^14,17,18,24^. However, due to the sparsity of BLA interneurons and the resulting low cell numbers per animal in *in vivo* recordings, a systematic classification of response patterns at the single cell level across interneuron subpopulations is still lacking. Furthermore, a characterization of interneuron plasticity would require to reliably record from the same cells over the course of several days, which is difficult to achieve in deep brain regions such as the amygdala.

To address this gap, we employed deep-brain calcium imaging with implanted lenses and miniature microscopes in freely behaving mice during an associative learning paradigm. By targeting all BLA inhibitory cells, we were able to follow large populations of interneurons at single cell resolution across days and provide the first classification of their response types during fear learning, memory expression and extinction. These data reveal that BLA interneurons show complex plastic responses at both the ensemble and single neuron level, with distinct cells being selectively activated or inhibited upon fear or extinction training. Using this classification, we further demonstrate that SST and VIP interneurons are differentially encoding conditioned high and low fear states, with heterogeneity at the single cell level, indicating the presence of functional subtypes within the classical interneuron subpopulations defined by molecular marker expression.

## RESULTS

### Deep-brain imaging of amygdala interneurons

To record the activity of identified BLA interneurons in freely behaving mice at single cell resolution, we employed a gradient refractive-index (GRIN) lens-based imaging approach in combination with miniaturised microscopes^18,25^ (Fig. 1A). Cre-dependent, virally mediated expression of GCaMP6f^26^ in the BLA of *GAD2-Cre* mice^27^ allowed for interneuron-specific Ca^2+^ imaging. Immunohistochemical analysis revealed that all major interneuron subpopulations (SST+, PV+, VIP+ and CCK+ interneurons) were targeted with this approach (Fig. 1B). Mice with head-mounted miniature microscopes underwent a four-day auditory fear conditioning and extinction paradigm (Fig. 1C, Methods). For habituation, mice explored context A and were presented with two different auditory cues (6 kHz and 12 kHz pure tones) used as CS+ and CS– in the fear conditioning session on the next day. For conditioning, the CS+ was paired with an aversive US in the form of a mild foot shock for five times in a different context B, while intermingled CS– presentations were used as control tones. For test and extinction days, the CS– was presented four times, followed by 12 CS+ stimuli in context A. The exact lens placements in BLA subnuclei were confirmed *post hoc ex vivo* and revealed that most implant sites were in the basal amygdala (N = 6 mice) while a minority was at the border between lateral and basal amygdala (N = 3; Fig. S1A). On average, we recorded 58 ± 6 BLA interneurons per mouse (N = 9 mice; Fig. S1B) stably within and across four days (Fig. 1D, Methods). BLA interneurons showed diverse spontaneous activity patterns, as well as cell-specific responses to auditory and aversive stimuli (Fig. 1E-F).

**Figure 1:**
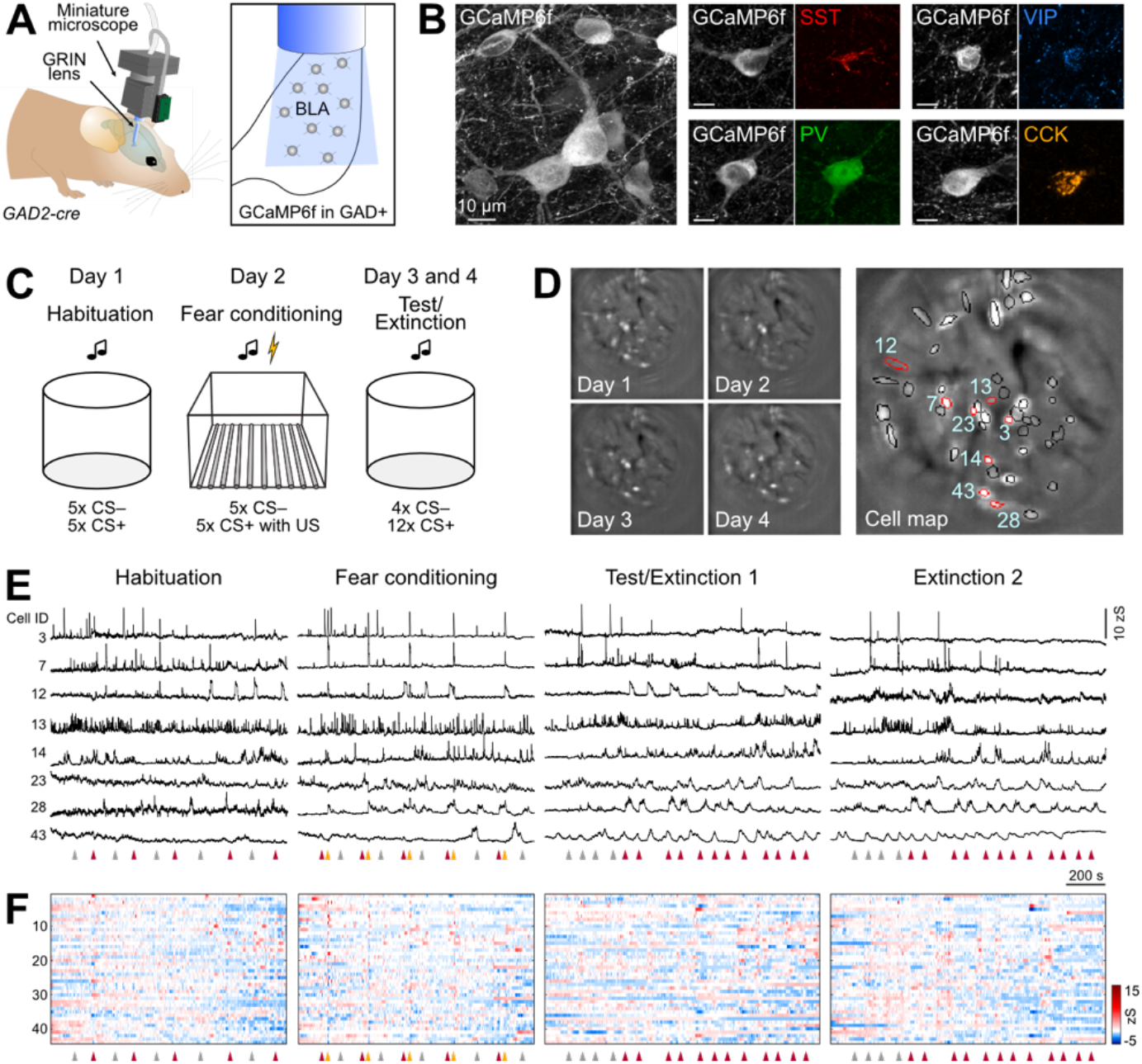
Imaging interneuron activity in the basolateral amygdala of freely moving mice. **A**, Schematic of the approach used for deep-brain calcium imaging of BLA interneurons in freely behaving *GAD2-Cre* mice. Recordings were obtained using a miniature microscope after virus injection for Cre-dependent GCaMP expression and implantation of a gradient-index (GRIN) lens into the basolateral amygdala. **B**, Representative confocal images of Cre-dependent GCaMP6f expression in BLA interneurons. Expression of all main interneuron markers was detected in GCaMP+ neurons (SST, somatostatin; PV, parvalbumin; VIP, vasoactive intestinal peptide; CCK, cholecystokinin). **C**, Scheme of the four-day discriminative auditory fear conditioning paradigm, consisting of habituation, fear conditioning, test and extinction sessions. **D**, Individual motion corrected fields of view (maximum intensity projection) of one example animal across the four-day paradigm (left) and the resulting cell map across all days (right). Circles indicate selected individual components. **E**, Representative example traces of selected interneurons across the four-day paradigm (highlighted with red outlines in D). **F**, Activity map of all identified interneurons from the example mouse in D across the entire paradigm (n = 44 cells). Arrowheads in E and F indicate starting points of CS+ (red), CS– (grey) and US (yellow).

### Mixed selectivity coding in amygdala interneurons

We initially focussed our analysis on the fear conditioning day. We could observe a diversity of cellular responses to the predictive CS+ cue, the aversive US and the neutral CS– control tone across neurons and animals (Fig. 2A-F). On average, CS+ and CS– led to a mild activation of BLA interneurons, while the aversive US induced a strong response (Fig. 2B). However, both inhibition and activation could be seen in individual cells for the different stimuli (Fig. 2A, F; see also Fig. 3A and Fig. S2A), with significantly more CS+ excited than inhibited cells (n = 519 cells; CS+ inhibited 22%, activated 36%) but similar proportions for CS– and US modulated cells (CS– inhibited 27%, activated 33%; US inhibited 36%, activated 40%). A majority of BLA interneurons responded to the auditory cues and the foot shock (Fig. 2C-D), with significantly higher fractions to the aversive stimulus (76 ± 2%) compared to the CS+ (58 ± 3%) and CS– tones (60 ± 2%; N = 9 mice). Mixed selectivity was found in subpopulations of interneurons that responded to combinations of the CS+, CS– or the aversive US, yet this was not enriched above chance level in the overall population (Fig. 2D-E). Furthermore, neurons responding selectively to individual stimuli were spatially intermingled with multisensory interneurons in the BLA, rather than locally clustered (Fig. 2G-H). No obvious differences were observed between animals with basal amygdala implant sites compared to mice with more dorsal lens placements at the border between lateral and basal amygdala (Fig. 2C, E). Together, this data shows that interneurons across BLA subregions are strongly modulated by auditory and aversive stimuli during conditioning. Furthermore, the coincidence of these two signals at the single cell level makes them ideal candidates for cellular plasticity during associative learning.

**Figure 2:**
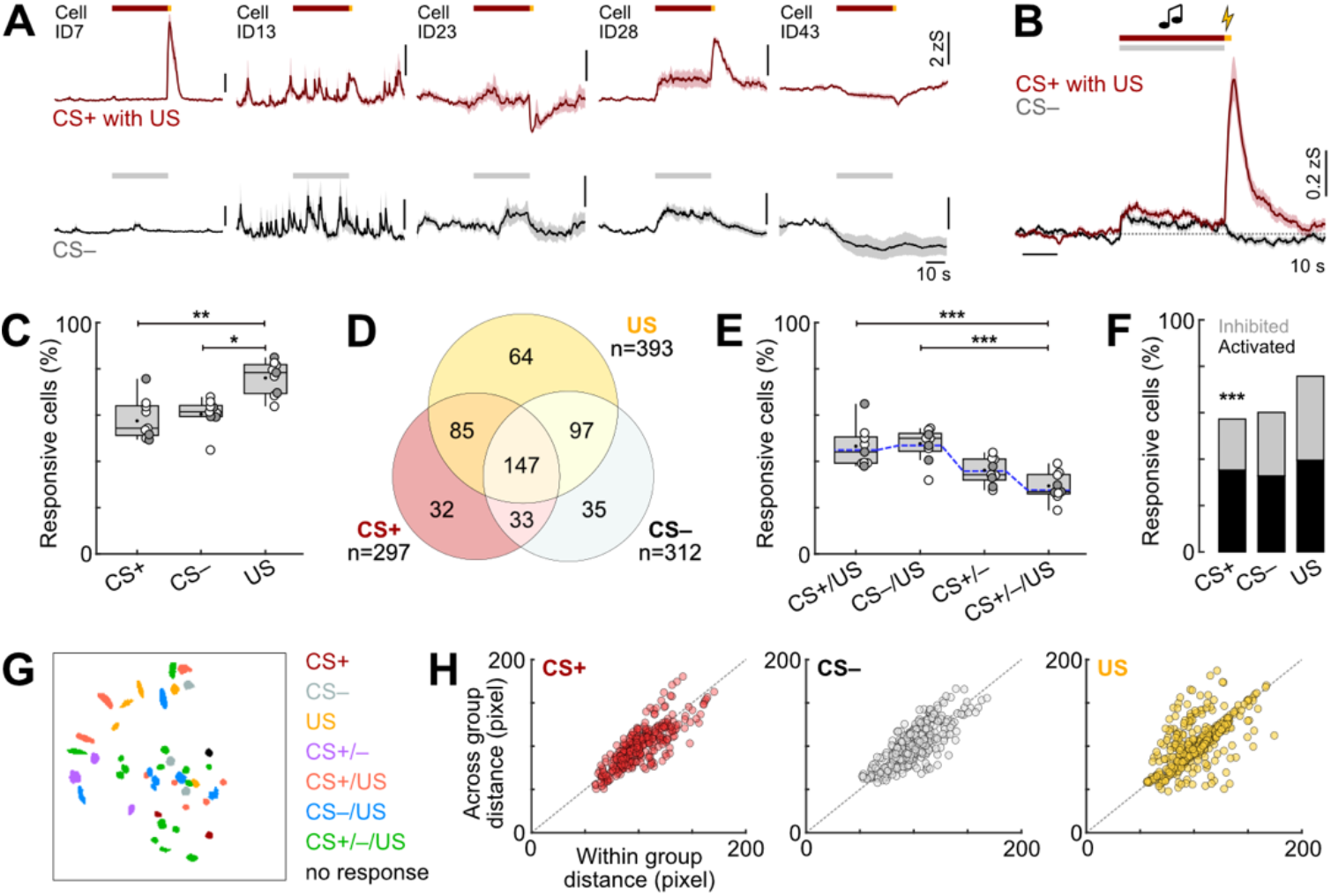
Mixed selectivity coding in amygdala interneurons during fear learning. **A**, Representative example traces illustrating diverse activity patterns of BLA interneurons, averaged across five pairings of CS+ with US (top) or five presentations of the CS– (bottom). Red/grey lines indicate CS+/CS– duration, yellow line US. Cell IDs correspond to recording shown in Figure 1. **B**, CS and US responses from all recorded BLA interneurons across five trials averaged across all mice (N = 9). **C**, Fraction of interneurons responsive to the CS+, CS– and US across distinct animals (N = 9). Friedman test (χ2 = 11.53), *p* = 0.0016, followed by Dunn’s multiple comparisons (CS+ vs. US, *p* = 0.0065; CS– vs. US, *p* = 0.0286). **D**, Overlap of CS+, CS– and US responsiveness in individual interneurons. **E**, Proportion of mixed selectivity CS+, CS– and US coding interneurons (N = 9). Blue line indicates chance overlap level. Friedman test (χ2 = 24.33), *p* < 0.0001, followed by Dunn’s multiple comparisons (CS+/US vs. CS+/CS–/US, *p* = 0.0004; CS–/US vs. CS+/CS–/US, *p* = 0.0002). **F**, Percentage of BLA interneurons with significantly increased or decreased calcium responses during distinct stimulus presentations (n = 519; Chi-Square test with Bonferroni correction: CS+ inhibited vs. activated, *p <* 0.0001). **G**, Example spatial map of mixed selectivity CS and US coding neurons (same animal as in Figure 1). **H**, Pairwise relationship (spatial distance) between ‘within response group’ and ‘across response group’ between recorded amygdala interneurons (n = 519) for CS+, CS– and US. Average traces in A and B are mean with s.e.m.; Tukey box-and-whisker plots in E and F show median values, 25^th^ and 75^th^ percentiles, and min to max whiskers with exception of outliers, dots indicate the mean. Circles represent individual animals (open circles, imaging sites in the basal amygdala (N = 6); filled circles, at the border of the lateral and basal amygdala (N = 3), see also Supplementary Fig. 1). \**p* < 0.05, \*\**p* < 0.01, \*\*\**p* < 0.001. Additional details of statistical analyses are provided in Supplementary Table 1.

**Figure 3:**
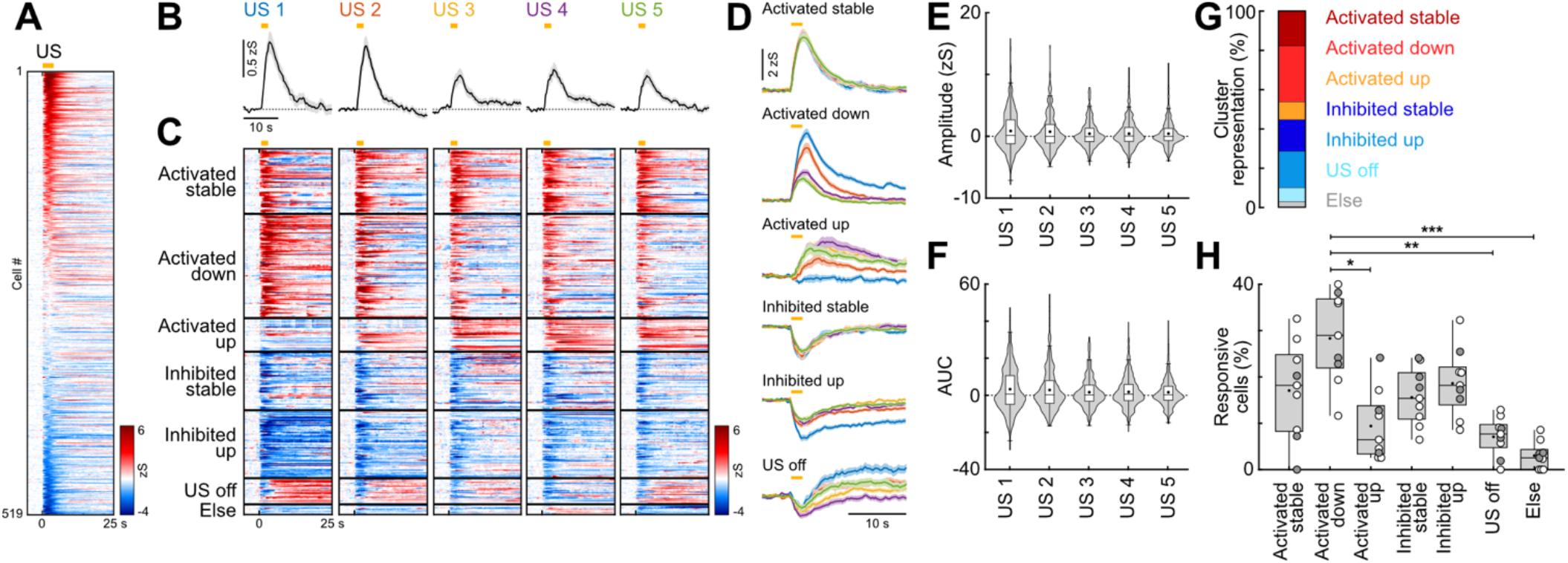
Interneuron responses to aversive stimuli are highly diverse and plastic. **A**, Heatmap of basolateral amygdala interneuron responses to the aversive US (averaged across all five presentations) sorted by response amplitude (n = 519 cells from N = 9 mice). Yellow line indicates US duration. **B**, Average traces of amygdala interneurons during the five aversive US presentations (n = 519). **C**, Heatmap of single cell US responses clustered into groups depending on their US response pattern across the five presentations (n = 393 responsive cells; ‘Activated stable’, n = 70; ‘Activated down’, n = 112; ‘Activated up’, n = 36; ‘Inhibited stable’, n = 63; ‘Inhibited up’, n = 73; ‘US off’, n = 28; ‘Else’, n = 11). **D**, Average traces of neuronal clusters in C. **E**, Average amplitude and **F**, average area under the curve (AUC) during the five US presentations (n = 519). **G**, Proportion of cells in US clusters. **H**, Fraction of interneurons according to US cluster membership across animals (N = 9). Friedman test (χ2 = 30.41), *p* < 0.0001, followed by Dunn’s multiple comparisons (‘Activated down’ vs. ‘Activated up’, *p* = 0.0327; ‘Activated down’ vs. ‘US off’, *p* = 0.0023; ‘Activated down’ vs. ‘Else’, *p* < 0.0001). Average traces in B and D are mean with s.e.m.; violin plots in E and F show distribution of all data points, Tukey box-and-whisker plots in E, F and H show median values, 25^th^ and 75^th^ percentiles, and min to max whiskers with exception of outliers, dots indicate the mean. Circles in H represent individual animals (open circles, imaging sites in the basal amygdala (N = 6); filled circles, at the border of the lateral and basal amygdala (N = 3), see also Supplementary Fig. 1). \**p* < 0.05, \*\**p* < 0.01, \*\*\**p* < 0.001. Additional details of statistical analyses are provided in Supplementary Table 1.

### Interneurons develop plastic responses during associative learning

Previous studies showed adaptive CS and US responses during learning for selected BLA interneuron populations^14,18,23,24^. However, a systematic classification of plasticity response types is still lacking, as previous results have been obtained using fibre photometry or single cell approaches with low cell numbers. Therefore, we next investigated how CS and US responses of individual BLA interneurons change during fear conditioning. On average, interneurons displayed decreasing US amplitudes over the course of the five CS+/US pairings (n = 519; Fig. 3B), as previously reported for VIP and PV BLA interneurons^18,23^. However, this gradual decline was not statistically significant for the population average (Fig. 3E-F). Next, we used a clustering approach to classify cellular responses of significantly modulated interneurons across the five US presentations (n = 393; Fig. 3C, D, G). This revealed distinct types of activated and inhibited patterns, including cells with stable US signalling, but also interneurons that down- or up-regulate their US response with repeated pairings, as well as post-US activated (‘US off’) cells. Across animals, the ‘Activated down’ cluster was most prominent (29%),compared to smaller fractions particularly of the ‘Activated up’ (9%) and ‘US off’ patterns (7%; Fig. 3H). This analysis illustrates that the population average during the aversive stimulus (Fig. 2B, 3B) is dominated by the activated cells, although a comparable fraction of cells displayed inhibition of their intrinsic activity.

We used a similar analysis approach to characterise CS responses of BLA interneurons, for which we could observe both activated and inhibited neurons across all five presentations of the CS+ and CS– (n = 519; Fig. S2A). On the population level, responses to both stimuli occurred during the initial presentations (Fig. 4A, D). However, at the end of the fear learning session only the CS+ induced a clear activation, with a significantly stronger response at the fifth presentation compared to the subsequent CS– (Fig. 4A, D, G), indicating stronger upregulation of interneuron activity during the predictive cue compared to tones that are not paired with an aversive outcome. We next used a clustering approach (see Methods) to define response types in individual interneurons that were significantly modulated by the CS+ (n = 297 cells) and CS– (n = 312; Fig. 4B-C and Fig. 4E-F). This demonstrated that both predictive CS+ and control CS– tones induced cellular plasticity during the conditioning paradigm. Neurons that were only activated or inhibited by the first of these tones were equally represented for both conditions (Fig. 4H). Comparable fractions of BLA interneurons were initially not modulated by the respective CS but became activated (‘Up’ cluster) or inhibited during learning (‘Down inhibited’). However, only for the CS+ a ‘Stable activated’ pattern emerged (24%). This was significantly different from the CS–, that was not associated with stably activated cells. Instead, a fraction of interneurons selectively downregulated their activity after the second presentation (‘Down activated’, 9%). None of these clusters were dominant when comparing their proportions across animals (Fig. S2B-C). Together, these results suggest that the increased CS+ activity that develops at the population level at the end of the conditioning session (Fig. 4A, G) is not mediated by an upregulation of interneuron activity across trials, but achieved by maintaining a stable activation pattern during the session. In contrast, interneurons produce a balanced excitation and inhibition to the CS– during progressive learning, leading to no noticeable CS response at the population level at the end of training (Fig. 4D).

**Figure 4:**
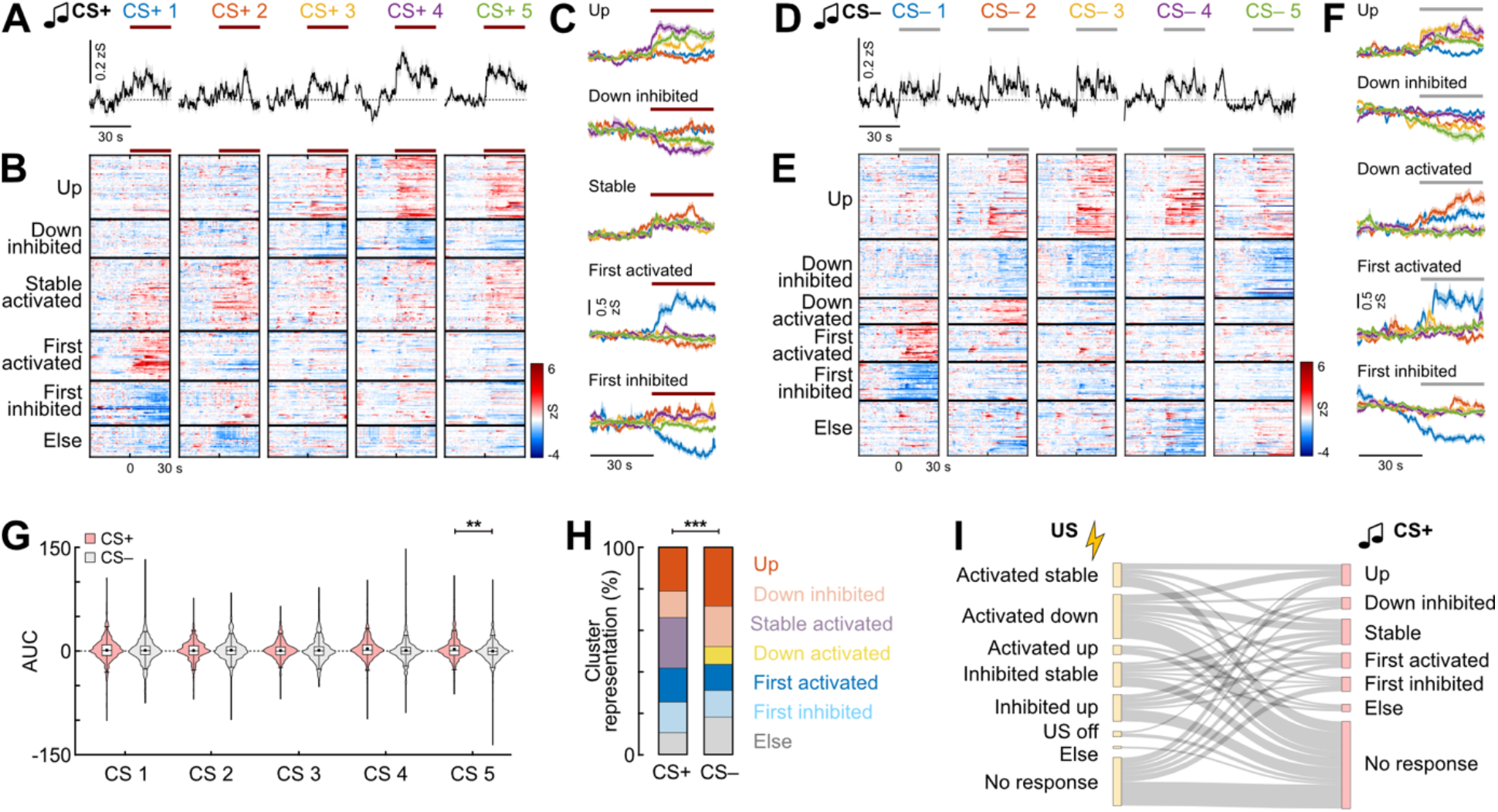
Interneuron responses to auditory cues depend on the predictive value of the stimulus. **A**, Average traces of BLA interneuron activity during five presentations of the predictive CS+ during conditioning (n = 519 cells from N = 9 mice). Line indicates CS duration. **B**, Heatmap of single cell CS+ responses clustered into groups depending on their CS+ response pattern across the five trials (n = 97 responsive cells; ‘Up’, n = 63; ‘Down inhibited’, n = 38; ‘Stable activated’, n = 72; ‘First activated’, n = 49; ‘First inhibited’, n = 44; ‘Else’, n = 31). **C**, Average traces of neuronal clusters in B. **D**, Average traces of BLA interneurons during five presentations of the CS– control tone during conditioning (n = 519 cells). **E**, Heatmap of single cell CS– responses clustered into groups depending on their response pattern across the five trials (n = 312 responsive cells; ‘Up’, n = 88; ‘Down inhibited’, n = 61; ‘Down activated’, n = 27; ‘First activated’, n = 39; ‘First inhibited’, n = 40; ‘Else’, n = 57). **F**, Average traces of neuronal clusters in (E). **G**, Area under the curve (AUC) for CS+ and CS– presentations during conditioning. Paired Wilcoxon test with Bonferroni correction; CS 5, CS+ vs. CS–, *p* = 0.0021; n = 519. **H**, Proportion of cells in CS+ and CS– clusters (CS+, n = 297; CS–, n = 312). Chi-Square test CS+ vs. CS– (χ2(6) = 117.19), *p* < 0.0001; *post hoc* Chi-Square test with Bonferroni correction; ‘Down activated’, *p* < 0.0001; ‘Stable’, *p* < 0.0001. **I**, Sankey plot illustrating the relationship of activity during the aversive US (Figure 3) with CS+ plasticity patterns. Average traces in A, C, D and F are mean with s.e.m.; violin plots in G show distribution of all data points, Tukey box-and-whisker plots in G show median values, 25^th^ and 75^th^ percentiles, and min to max whiskers with exception of outliers. \*\**p* < 0.01, \*\*\**p* < 0.001. Additional details of statistical analyses are provided in Supplementary Table 1.

To investigate whether CS+ plasticity types depend on US activation, we further compared the US and CS+ response patterns of individual neurons during the fear conditioning paradigm (Fig. 4I). These data demonstrate that CS+ activity evolves independently of US activation, as adaptive CS+ responses were observed in interneurons activated and inhibited by the US, but also in cells without any detectable US response at the level of the soma. In summary, recording the activity of large BLA interneuron populations at the single cell level during auditory fear learning revealed distinct plastic response types, both for the aversive US teaching signal but also for tone cues. The significant differences between CS+ and CS– illustrate differential plasticity of BLA interneurons depending on the predictive value of a stimulus.

### Amygdala interneurons signal high and low fear states across conditioning and extinction

We next assessed how BLA interneuron CS responses change across days during fear expression and extinction. To this end, we first compared neuronal responses to the CS+ and CS– at the population level before conditioning (habituation), after conditioning (24 h after learning) and after extinction in the same behavioural context. For both CS+ and CS–, we could observe tone activation on the population level in the habituation session (n = 519 cells; Fig. 5A, D and Fig. S3), which overall decreased across the session (Fig. S4). At the single cell level, responsive neurons could be classified into activated and inhibited cells, with comparable proportions of these clusters between CS+ and CS– before conditioning (Fig. S4A-E). Since 6 kHz and 12 kHz tones were counterbalanced as CS+ and CS– in the experimental cohort, we further analysed whether these frequencies induced differential activity in BLA interneurons (Fig. S4F-J). We could observe similar clusters of activated and inhibited neurons with no bias for either frequency.

**Figure 5:**
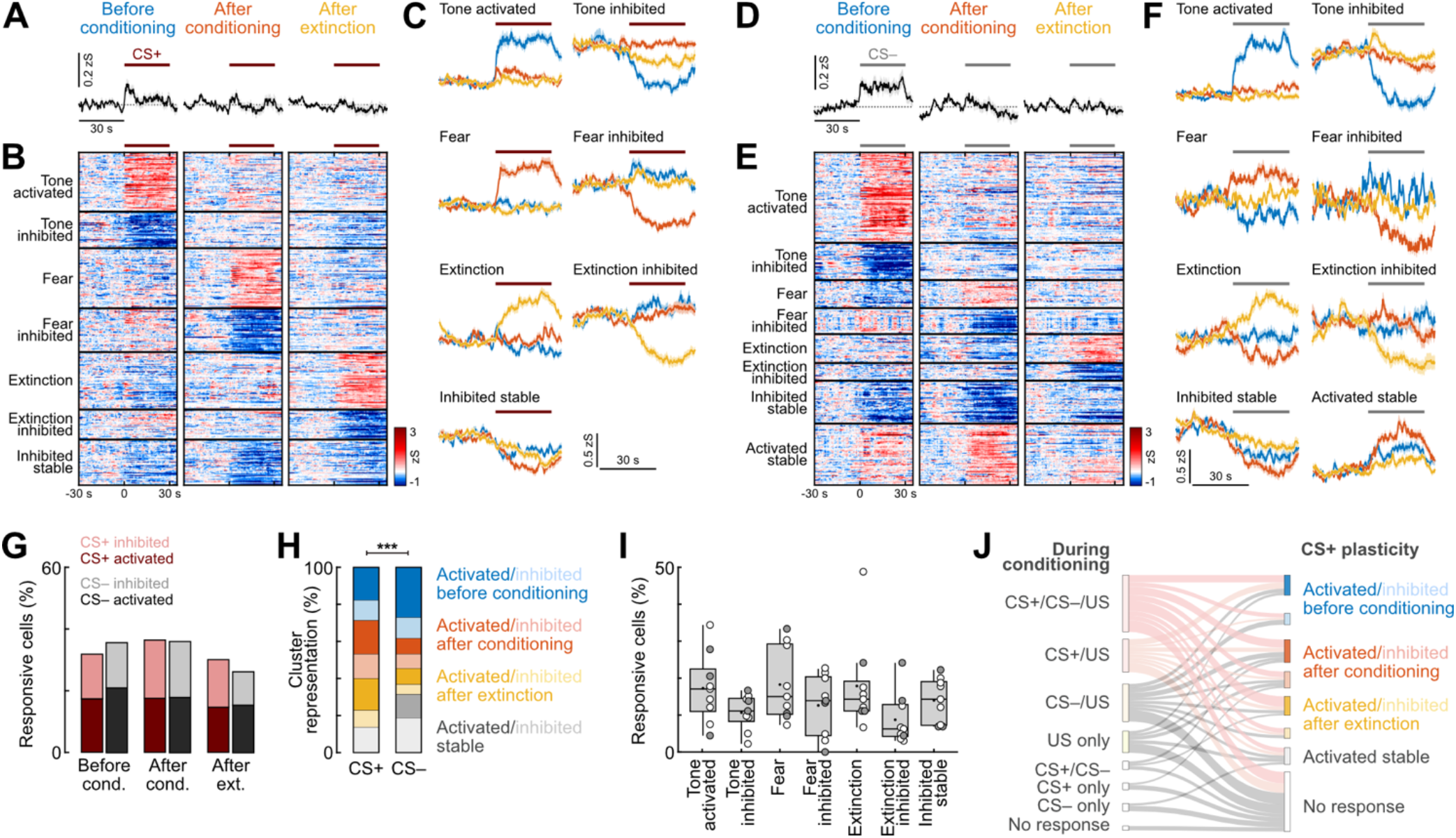
Amygdala interneurons encode high and low fear states. **A**, Average traces of BLA interneuron activity during the CS+ before conditioning, after conditioning and after extinction (n = 519 cells from N = 9 mice). Line indicates CS duration. **B**, Heatmap of single cell CS+ responses clustered into groups depending on their CS+ response pattern (n = 365 responsive neurons; ‘Tone activated’/activated before conditioning, n = 65; ‘Tone inhibited’/inhibited before conditioning, n = 40; ‘Fear’/activated after conditioning, n = 66; ‘Fear inhibited’/inhibited after conditioning, n = 48; ‘Extinction’/activated after extinction, n = 63; ‘Extinction inhibited’/inhibited after extinction, n = 33; ‘Inhibited stable’, n = 50). **C**, Average traces of CS+ clusters in B. **D**, Average traces of BLA interneuron activity during the CS– before conditioning, after conditioning and after extinction (n = 519 cells). **E**, Heatmap of single cell CS– responses clustered into groups depending on their CS– response pattern (n = 357 responsive neurons; ‘Tone activated’/activated before conditioning, n = 97; ‘Tone inhibited’/inhibited before conditioning, n = 40; ‘Fear’/activated after conditioning, n = 30; ‘Fear inhibited’/inhibited after conditioning, n = 28; ‘Extinction’/activated after extinction, n = 31; ‘Extinction inhibited’/inhibited after extinction, n = 19; ‘Inhibited stable’, n = 46; ‘Activated stable’, n = 66). **F**, Average traces of CS– clusters in (E). **G**, Proportions of responsive neurons across the behavioural paradigm (n = 519). **H**, Proportion of cells in CS+ and CS– clusters (CS+, n = 365; CS–, n = 357). Chi-Square test CS+ vs. CS– (χ2(7) = 105.84), *p* < 0.0001; *post hoc* Chi-Square test with Bonferroni correction; ‘Activated before conditioning’, *p =* 0.0275; ‘Activated after conditioning’, *p* = 0.0016; ‘Activated after extinction’, *p* = 0.0074; ‘Activated stable’, *p* < 0.0001. **I**, Fraction of interneurons according to CS+ cluster membership across animals (N = 9). **J**, Sankey plot illustrating the relationship of activity during fear conditioning with across-day CS+ plasticity patterns. Average traces in A, C, D and F are mean with s.e.m.; Tukey box-and-whisker plots in I show median values, 25^th^ and 75^th^ percentiles, and min to max whiskers with exception of outliers, dots indicate the mean. Circles in I represent individual animals (open circles, imaging sites in the basal amygdala (N = 6); filled circles, at the border of the lateral and basal amygdala (N = 3), see also Supplementary Fig. 1). \*\*\**p* < 0.001. Additional details of statistical analyses are provided in Supplementary Table 1.

Across days, no clear response of BLA interneurons could be observed after conditioning or after extinction for either CS on the population level (Fig. 5A, D). Yet, individual neurons showed significant activation or inhibition (Fig. S3A-B, Fig. 5G). Overall, proportions of neurons significantly activated or inhibited by the auditory cues were comparable between CS+ and CS– on all behavioural days (Fig. 5G). Clustering CS responses of significantly modulated cells across all days (CS+, n = 365; CS–, n = 357) revealed the emergence of distinct response types in individual interneurons (Fig. 5B-C and Fig. 5E-F). For both tones, these were associated with neutral tone presentations (activated or inhibited only during habituation), high fear states (only after conditioning) and low fear states (only after extinction). A subset of interneurons was further stably inhibited or activated across days, although the stably activated cluster could only be observed for CS– control tones (19%), suggesting that dynamic changes in CS coding across days are associated with aversive outcomes. While the predictive CS+ induced significantly stronger activation at the single cell level after conditioning (CS+ 18%, CS– 8%) and after extinction (CS+ 17%, CS– 9%), a larger fraction of interneurons was selectively activated by the CS– only before conditioning (CS+ 18%, CS– 27%; Fig. 5H). This cluster ‘Activated before conditioning’ was further significantly enriched across all animals for the CS– (N = 9 mice; Fig. S3D), whereas CS+ response types were evenly represented (Fig. 5I). Together, this data indicates that individual interneurons develop prominent responses to predictive cues in high and low fear states associated with learning and extinction, which could not be detected by monitoring the population average. Further, a comparison between responses during conditioning with fear and extinction coding revealed that this plasticity of BLA interneurons is independent of US activation during conditioning, indicating that a CS-US coincidence, at least on the level of somatic Ca^2+^ read-outs, is not necessary for memory-associated cellular plasticity across days (Fig. 5J).

### Inhibitory population activity changes day-to-day without losing overall stimulus representation

To determine if stimulus identity could be decoded from interneuron activity patterns, we trained multiclass decoders using binary linear support vector machines (SVM) on each training day^28,29^. To account for variations in cell population sizes between animals, we selected 37 cells randomly from each animal and report the average decoder accuracy of 100 independent iterations. Decoders accurately distinguished between CS+, CS–, and baseline using interneuron activity in all sessions, with a mean accuracy of 94 ± 3% (Fig. 6A). This accuracy was not due to better decoding of one class over another, as all three classes showed high precision, recall, and F1 scores (see Methods for details), indicating a balanced performance on each day (Fig. S5D). Decoders trained solely on CS responsive interneurons (i. e. significantly responsive to CS+ or CS–, see Fig. 4 and Fig. 5) maintained similar accuracy (92 ± 4%). However, using non-CS responsive interneurons reduced the accuracy to 76 ± 11%, which was nevertheless higher than chance-level accuracy obtained from decoders trained on randomly shuffled labels (47 ± 0.2%). To eliminate the possibility that discrepancies in non-CS responsive interneuron decoder precision were attributable to a reduced number of available cells, we trained control decoders on a random subset of all interneurons, ensuring an equivalent number of cells to those used in the non-CS decoders (Fig. S5A).

**Figure 6:**
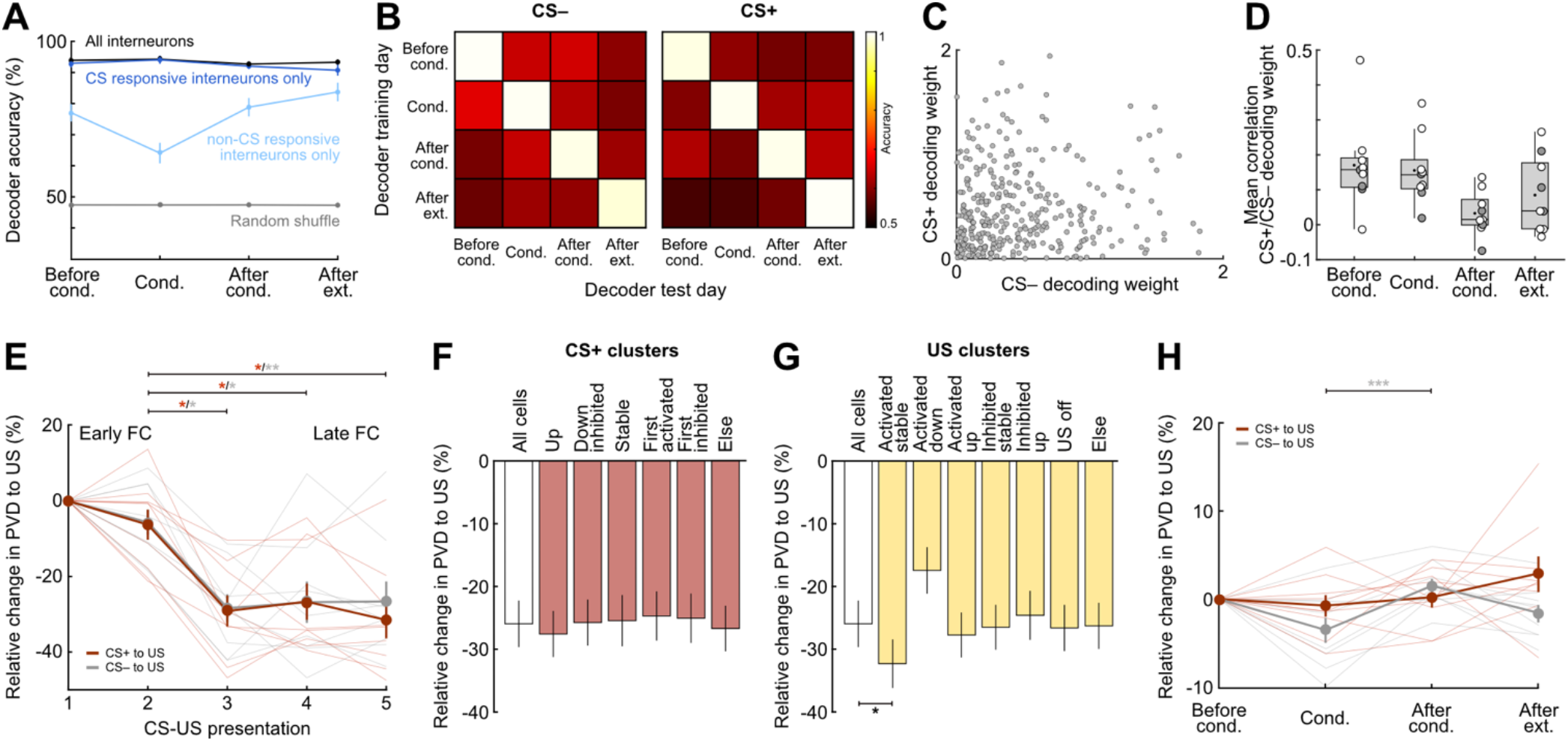
Interneuron population activity dynamically changes day to day without losing overall stimulus representation. **A**, Mean accuracy of multiclass intra-day decoder of CS+, CS– and baseline for each day of the behavioural paradigm across all animals (N = 9 mice) and iterations (n = 100 iterations). Decoding accuracy is also shown for decoders based only on CS responsive cells, non-CS responsive cells and randomly shuffled training labels. **B**, Accuracy of intra-day and inter-day two-way decoders trained to classify CS+/baseline and CS–/baseline. **C**, Example scatterplot showing the absolute value of the decoding weight for the CS+ and CS– for all interneurons used in one iteration after conditioning (n = 333 cells from N = 9 mice). **D**, Mean correlation of 100 iterations per animal between the absolute value of the CS+ and CS– decoding weight in each session (N = 9). **E**, Mean relative change in population vector distance (PVD) between the CS+ and US (red) and between the CS– and US (grey) across the 5 CS-US pairings during fear conditioning (N = 9). Changes in PVD were normalized to the first CS-US pairing. CS+/US distance Friedman test (χ2 = 13.9), *p* = 0.0030, followed by Dunn’s multiple comparisons (pairing 2 vs. 3, *p* = 0.0115; pairing 2 vs. 4, *p* = 0.0209; pairing 2 vs. 5, *p* = 0.0115). CS–/US distance Friedman test (χ2 = 14.2), *p* = 0.0026, followed by Dunn’s multiple comparisons (pairing 2 vs. 3, *p* = 0.023; pairing 2 vs. 4, *p* = 0.047; pairing 2 vs. 5, *p* = 0.023). **F**, Mean change between early and late fear conditioning was calculated as the difference between the average difference during pairings 1-2 vs. 4-5 for each animal (N = 9). Decomposition of the contribution of interneurons based on their CS+ activity patterns (see Figure 4) was performed by removing the cells of each CS+ cluster and recalculating the mean change in PVD. Friedman test (χ2 = 13.9), *p* = 0.0304, followed by Dunn’s multiple comparisons (non-significant). **G**, Average change between early and late fear conditioning for US clusters (see Figure 3), contribution calculated as in E (N = 9). Friedman test (χ2 = 36.0), *p* < 0.0001, followed by Dunn’s multiple comparisons (‘All cells’ vs. ‘Activated stable’, *p* = 0.0272). **H**, Same as D, but calculated across days (N = 9). Results were normalized to the average distance between CS-US in habituation. CS–/US distance Friedman test (χ2 = 14.2), *p* < 0.0001, followed by Dunn’s multiple comparisons (‘Conditioning’ vs. ‘After conditioning’, *p* = 0.0005). Data is presented as mean with s.e.m.; except for D showing Tukey box-and-whisker plots with median values, 25^th^ and 75^th^ percentiles, and min to max whiskers with exception of outliers, dots indicate the mean. Circles in D represent individual animals (open circles, imaging sites in the basal amygdala (N = 6); filled circles, at the border of the lateral and basal amygdala (N = 3), see also Supplementary Fig. 1). Semi-transparent lines in E and H represent individual animals. \**p* < 0.05, \*\**p* < 0.01, \*\*\**p* < 0.001. Additional details of statistical analyses are provided in Supplementary Table 1.

We next assessed changes in CS+ and CS– encoding across days. To this end, we trained two-way SVMs to decode CS+ vs. baseline, and CS– vs. baseline on each day. These models were then used to decode stimuli identity on subsequent days. Although stimuli identity could be decoded within each day (mean intraday accuracy: CS+, 95 ± 4%; CS–, 95 ± 3%), the same interneurons could not decode stimuli identity on another day. Decoding accuracy dropped close to chance level to an average of 60 ± 3% for CS+ and 62 ± 3% for CS– when using a model trained on a different day (Fig. 6B). This decrease was mainly due to poor decoding of CS+ and CS–, as accuracy, precision, and F1 score were higher for baseline than for CS+ or CS– (Fig. S5E-F). This indicates that individual interneurons change dynamically from day to day, but information remains encoded at the population level on individual days.

To investigate interneuron selectivity for CS+ or CS–, we obtained the corresponding absolute decoding weights from the two-way decoders for each interneuron and calculated the correlation between them. If interneurons selectively encoded CS+ or CS–, one would expect a negative correlation between their weights, where a higher decoding weight for CS+ would corresponds to a lower decoding weight for CS– (see^29^). However, the correlation between CS+ and CS– decoding weights was close to zero after the conditioning session (Fig. 6C-D), suggesting that interneurons were not tuned to a stimulus and display mixed selectivity and broad tuning, consistent with our single cell analysis results (Fig. 2). Controls calculating the correlation between CS+ decoding weights showed high correlation (Fig. S5B-C).

Moreover, we evaluated how fear conditioning altered the differentiability of CS+ and CS– from the US as learning progressed. For population vector distance (PVD) analysis, we calculated the Euclidean distance between the evoked population vector responses to CS+/CS– and the US for the multidimensional space of n interneurons in each individual mouse^30^. To probe whether the population evoked responses were getting closer or farther away as fear conditioning progressed, we normalized the PVD change to the distance between CS and US during the first pairing. We found that by the end of the session (pairings 4 and 5), the distance between CS+ and US, as well as CS– and US, had significantly decreased, averaging a reduction of 29 ± 14% and 27 ± 16%, respectively (Fig. 6E). This indicates that both stimuli’s representations became closer to the US over the course of training. Distinct activation patterns of interneurons might contribute differently to this distance decrease between CS+ and US. Therefore, we re-ran the analysis while removing interneurons from different clusters based on their US and CS+ activity patterns during fear conditioning and calculated the difference between early (pairings 1-2) and late (pairings 4-5) fear conditioning. Removing interneurons of the CS+ clusters had no significant effect on the distance (Fig. 6F). Interestingly, removal of interneurons of the ‘Activated stable’ US cluster decreased the distance between CS+ and US even more (‘Activated stable’, 32% vs. ‘All cells’, 26%; Fig. 6G), while removing interneurons with the ‘Activated down’ pattern increased the distance, albeit not significantly (17%). Thus, for the most part, the highly-defined interneuron response clusters for CS+ or US were not critical for the change in PVD between the CS+ and US during conditioning.

We further examined PVD changes across learning and extinction and found that the encoding for both CS+ and CS– with respect to the US remained overall stable, with average changes across days of 0.8 ± 4.6% and 1.2 ± 3.9%, respectively (Fig. 6H). The CS– showed statistically significant fluctuations, moving closer to the US during fear learning, farther away after conditioning, and returning to baseline after extinction. However, these changes were minimal (before conditioning, -3.5 ± 4.2%; after conditioning, 1.5 ± 2.6%; after extinction, -1.6 ± 3.0%) and exhibited high variance in individual animals. Thus, unlike BLA PN ensembles that display a lasting decreased in PVD and thus an increase in the similarity of CS+ and US representations after conditioning^28^, amygdala interneurons showed comparably stable representations of both CS+ and CS– across fear learning and extinction. Overall, our results indicate that BLA interneurons undergo heterogeneous plastic changes in single cell response patterns, yet representations of conditioned stimuli are stably encoded at the population level across fear learning and extinction.

### Molecular interneuron subpopulations contribute differently to the encoding of fear states

Finally, we aimed to identify how distinct molecularly defined interneuron subpopulations would contribute to the activity patterns we detected with the unbiased *GAD2-Cre* imaging approach. Given their previously proposed opposing roles during fear learning^14,18,23^, we chose to characterize response dynamics in SST and VIP BLA interneurons. To this end, we performed experiments in *SST-Cre* and *VIP-Cre* mice and recorded these interneuron subpopulations across the learning paradigm (Fig. S6, S7). On average, we could reliably follow 29 ± 5 SST interneurons per animal (N = 4 mice; Fig. S6B) across the four days, and 25 ± 3 VIP cells per mouse (N = 6).

We first re-analysed the previously published dataset from the fear conditioning day^18^ with a novel focus on response plasticity during learning. When comparing activity at the population level, stronger US activation was seen in VIP interneurons compared to SST cells (SST, n = 114 cells; VIP, n = 101 cells; Fig. 7A-C). The average US response was stable in SST interneurons across the five pairings with the CS+, while it decreased in VIP cells, as previously reported^18^. Yet, overall VIP activation was still significantly stronger compared to SST interneurons at the end of the session (Fig. 7B). At the single cell level, a higher fraction of VIP interneurons was significantly activated by the US and a larger proportion of SST cells significantly inhibited (Fig. 7G). To identify US response types in the two interneuron populations, we repeated the clustering of activity patterns based on the categories previously established with the unbiased imaging approach. Significantly more VIP interneurons were found in the ‘Activated stable’ group, which represented almost half of the VIP responses (SST 16%, VIP 48%). In contrast, the cluster ‘Activated up’ which gradually develops US responses during fear learning was only detectable in SST interneurons (6%). While SST interneurons also showed higher fractions of US inhibited cells (‘Inhibited stable’: SST 26%, VIP 10%; ‘Inhibited up’: SST 17%, VIP 8%), the different cluster representation was not found to be statistically significant in our dataset. Overall, these results show that differences in aversive US coding between VIP and SST interneurons are mainly driven by stable activation and inhibition in these populations, respectively.

**Figure 7:**
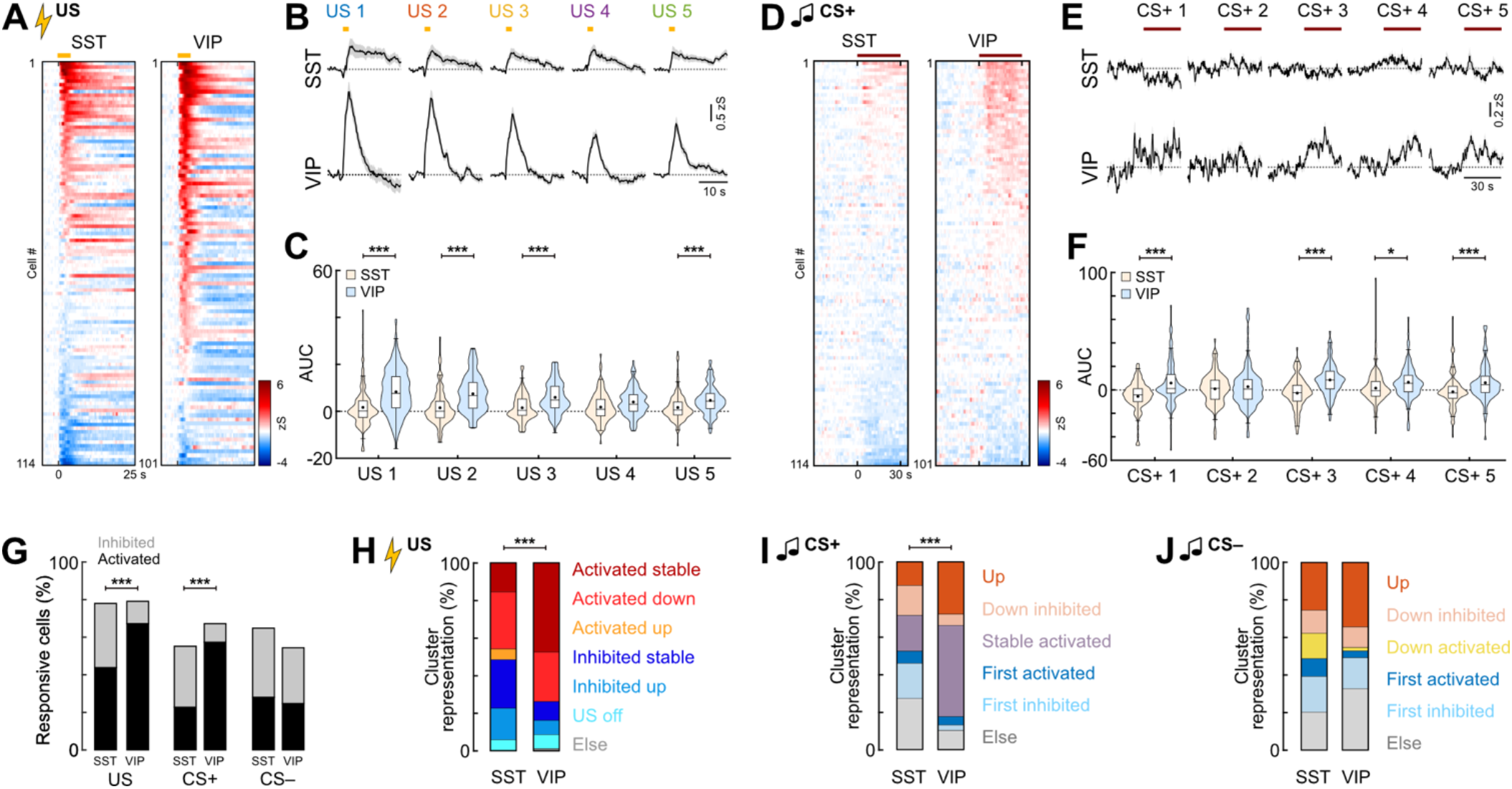
Differential activity during fear learning in molecular interneuron subpopulations. **A**, Heatmap of SST and VIP interneuron responses to the aversive US (averaged across all five presentations), sorted by response amplitude (SST, n = 114 cells from N = 4 mice; VIP, n = 101, N = 4). **B**, Average US responses in SST and VIP BLA interneurons during fear learning during the five trials (SST, n = 114; VIP, n = 101). Yellow line indicates US duration. **C**, Area under the curve (AUC) during the aversive US across all five trials. Paired Wilcoxon test with Bonferroni correction, SST vs. VIP; US 1, *p* < 0.0001; US 2, *p* < 0.0001; US 3, *p* < 0.0001; US 5, *p* = 0.0005. **D**, Heatmap of SST and VIP BLA interneuron responses to the predictive CS+ during conditioning (averaged across all five presentations), sorted by individual response amplitude (SST, n = 114 cells; VIP, n = 101). Line indicates CS duration. **D**, Average CS+ responses in SST and VIP interneurons across the five presentations (SST, n = 114; VIP, n = 101). **E**, Area under the curve (AUC) during CS+ presentations in conditioning for SST and VIP interneurons SST, n = 114; VIP, n = 101). Paired Wilcoxon test with Bonferroni correction; SST vs. VIP; CS+ 1, *p* < 0.0001; CS+ 3, *p* < 0.0001; CS+ 4, *p* = 0.0287; CS+ 5, *p* = 0.0003. **G**, Proportions of responsive neurons during fear learning (SST, n = 114; VIP, n = 101). CS+, Chi-Square test (χ2(2) = 30.885), *p* < 0.0001; SST vs. VIP, *post hoc* Chi-Square test with Bonferroni correction, ‘Activated’, *p* < 0.0001; ‘Inhibited’, p = 0.0004; US, Chi-Square test (χ2(2) = 16.663), *p* < 0.0001; SST vs. VIP, *post hoc* Chi-Square test with Bonferroni correction, ‘Activated’, p = 0.0028; ‘Inhibited’, *p* = 0.0007. **H**, Proportions of cells in US clusters for SST and VIP interneurons (SST, n = 89; VIP, n = 80). Chi-Square test (χ2(6) = 28.635), *p* < 0.0001; SST vs. VIP, *post hoc* Chi-Square test with Bonferroni correction, ‘Activated stable’, *p* = 0.0001. **I**, Proportion of cells in CS+ clusters for SST and VIP interneurons (SST, n = 63; VIP, n = 68). Chi-Square test CS+ vs. CS– (χ2(5) = 28.155), *p* < 0.0001; *post hoc* Chi-Square test with Bonferroni correction; ‘Stable activated’, *p* = 0.0046; ‘First inhibited’, *p* = 0.0418. **J**, Proportion of cells in CS– clusters (SST, n = 74; VIP, n = 55). Average traces in B and E are mean with s.e.m.; violin plots in C and F show distribution of all data points, Tukey box-and-whisker plots median values, 25^th^ and 75^th^ percentiles, and min to max whiskers with exception of outliers, dots indicate the mean. \**p* < 0.05, \*\*\**p* < 0.001. Additional details of statistical analyses are provided in Supplementary Table 1.

Next, we addressed how CS coding in BLA interneuron subpopulations changes during associative learning. We observed significant differences to the predictive CS+ between the SST and VIP subpopulations. At the population level, we found a stronger CS+ activation in VIP interneurons (Fig. 7D-F). This was reflected in stronger CS+ activation in individual VIP compared to SST cells, which were predominantly inhibited (Fig. 7G). Although SST cells appeared to be more activated across all five CS– presentations (Fig. S8E), differences in CS– activity could not be detected on the population level (Fig. S8F-G), nor in the fractions of significantly modulated cells (Fig. 7G). We further used the clustering approach to assign the previously determined classification of CS responses (Fig. 4) to the molecular interneuron subpopulations (Fig. S8C-D, H-I). For the CS+, we detected a significantly different cluster distribution between SST and VIP interneurons (Fig. 7I), with a higher fraction of stably activated neurons in VIP (SST 19%, VIP 49%), but smaller fraction of cells inhibited by the first CS+ (SST 19%, VIP 3%). In contrast, no significant differences in CS– cluster distribution could be observed between the two interneuron types (Fig. 7J). Overall, this data demonstrates that interneuron subpopulations are highly diverse even within a molecular subpopulation, with heterogeneous plasticity patterns visible for both activated and inhibited neurons. However, our analysis suggests that certain activity patterns are enriched in interneuron subpopulations. For example, given the high fraction of stable CS+ activated cells in the VIP interneurons, the noticeable CS+ responses we observed in the general BLA interneuron population during fear learning (Fig. 4) could be strongly driven by these cells.

Finally, we compared SST and VIP activity patterns across fear learning and extinction (SST, n = 114 cells from N = 4 mice; VIP, n = 152, N = 6). Like the conditioning day, differential responses between the subpopulations were mainly visible for the conditioned CS+ tone, and less for the CS– control tone. On the population level, SST interneurons were initially predominantly inhibited during the habituation session, however, upon extinction showed increased CS+ responses (Fig. 8A-C). In contrast, VIP interneurons were activated during habituation and after conditioning but showed suppressed activity after extinction. In comparison, this led to a stronger signalling of VIP interneurons upon neutral – but novel – tone presentations in habituation, while SST interneurons dominated after extinction (Fig. 8A-C). Analysis of the fractions of significantly modulated cells demonstrated that these differences are driven by suppression of SST interneuron activity before and after conditioning, since these cells showed significantly higher fractions of inhibited neurons (Fig. 8D). In contrast, after extinction, SST interneurons displayed more excitatory responses and a reduced fraction of unresponsive cells. This effect was not detectable for the CS– control tone. Only after conditioning, SST interneurons showed more inhibition and VIP cells more activation during the CS– (Fig. S9A-D). No differences in the number of cells significantly modulated by the CS– were detected between the two analysed interneuron subpopulations over the course of learning and extinction (Fig. S9D). For both the CS+ and the CS–, clustering of CS responses revealed a higher proportion of extinction activated neurons in SST interneurons (SST 13%, VIP 7%) but a higher proportion of VIP cells stably activated during the entire paradigm (SST 12%, VIP 27%; Fig. 8E and Fig. S9E-I). While none of these disparities between the two subpopulations were found to be statistically different for the CS+, the higher proportion of VIP neurons stably encoding the CS– across days was found to be significant. Taken together, our data demonstrate a stronger activation of VIP interneurons to novel stimuli and in conditioned high fear states (CS before and after conditioning). In contrast, higher activity in SST neurons emerges in low fear states after extinction, indicating that these cells could preferentially signal safety conditions. However, a detailed analysis of response patterns showed that all CS response types can be detected in both molecular subpopulations. This was also reflected in a bimodal distribution of CS response magnitudes, e.g. in individual SST interneurons after fear extinction (Fig. 8C), indicating the presence of further functional subtypes within these classical interneuron subpopulations.

**Figure 8:**
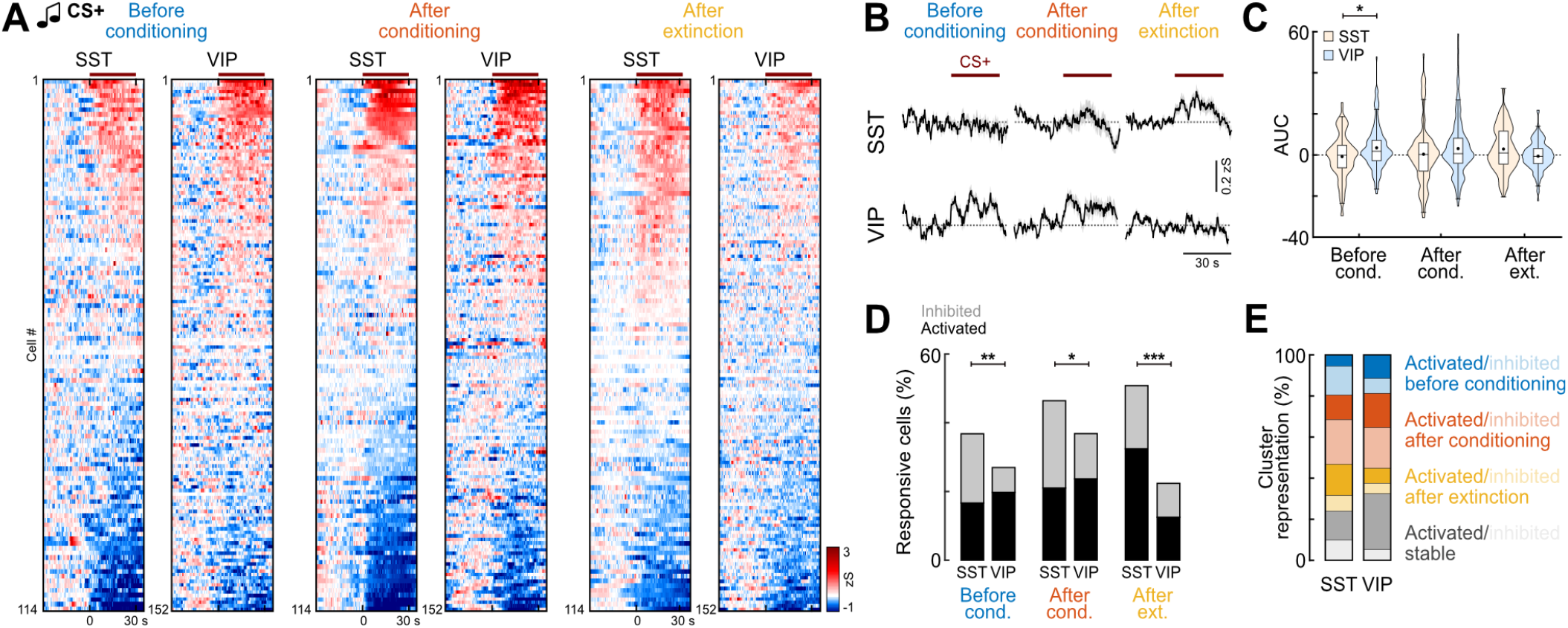
Interneuron subpopulations contribute differently to the encoding of fear states. **A**, Heatmap of SST and VIP BLA interneuron responses to the predictive CS+ before conditioning, after conditioning and after extinction (averaged across four presentations each, as used later for clustering), sorted by individual response amplitude (SST, n = 114 cells from N = 4 mice; VIP, n = 152, N = 6). Line indicates CS duration. **B**, Corresponding average CS+ responses in SST and VIP interneurons across days (SST, n = 114; VIP, n = 152). **C**, Area under the curve (AUC) during CS+ presentations in conditioning for SST and VIP interneurons SST, n = 114; VIP, n = 152). Mann-Whitney test with Bonferroni correction; SST vs. VIP; ‘Before conditioning’, *p* = 0.0154. **D**, Proportions of responsive neurons across the behavioural paradigm (SST, n = 114; VIP, n = 152). ‘Before conditioning’, Chi-Square test (χ2(2) = 9.7873), *p* = 0.0075, SST vs. VIP, *post hoc* Chi-Square test with Bonferroni correction, ‘Inhibited’, *p* = 0.0098; ‘After conditioning’, Chi-Square test (χ2(2) = 6.5609), *p* = 0.0376, SST vs. VIP, *post hoc* Chi-Square test with Bonferroni correction, ‘Inhibited’, *p* = 0.0496; ‘After extinction’, Chi-Square test (χ2(2) = 23.938), *p* < 0.0001, SST vs. VIP, *post hoc* Chi-Square test with Bonferroni correction, ‘Activated’, *p* = 0.0005, ‘No response’, *p* < 0.0001. **E**, Proportion of cells in CS+ clusters for SST and VIP interneurons (SST, n = 92; VIP, n = 96). Average traces in B are mean with s.e.m.; violin plots in C show distribution of all data points, Tukey box-and-whisker plots in C show median values, 25^th^ and 75^th^ percentiles, and min to max whiskers with exception of outliers, dots indicate the mean. \**p* < 0.05, \*\**p* < 0.01, \*\*\**p* < 0.001. Additional details of statistical analyses are provided in Supplementary Table 1.

## DISCUSSION

Here, we used deep-brain imaging to follow large populations of amygdala interneurons at single cell resolution across days and provide the first classification of their response types during fear learning, memory expression and extinction. We report that similar to neighbouring PNs^25,28,31^, BLA interneurons develop complex activity patterns with plastic changes across associative fear learning and extinction. This plasticity was seen both at the level of individual cells and the neuronal population coding, and differed for distinct molecular interneuron subpopulations.

### Encoding of high and low fear states in interneuron subpopulations

At the single cell level, BLA interneurons most prominently responded to the instructive US foot shock, but also to the predictive CS+ and control CS– tones during conditioning (Fig. 2). Analysis of the population average of BLA interneurons suggested a strong activation that declined over the course of repeated US presentations and thus predictive learning. At the same time, CS+ but not CS– responses on average increased in BLA interneurons. However, clustering of individual neuronal responses revealed highly diverse cellular responses beyond uniform activation. BLA interneurons showed plastic responses to both the predictive CS+ and the control CS–. Yet, CS+ and CS– functional clusters differed within the general inhibitory population – cells stably-activated across all five presentations were selectively detected during the CS+, while a cluster of activated neurons that decreased their responses over the trials were only present during the CS–. This suggests that the overall increased CS+ response in the interneuron population was not simply caused by an upregulation of individual activity across trials, but was the result of stable activation of a subset of interneurons throughout the session, while CS– clusters displayed balanced up- and downregulation.

Across fear learning and extinction, amygdala interneurons showed similar CS activity patterns as previously described for PNs^25,28,31^, such as selectively increased activity before conditioning, after fear learning and after extinction (Fig. 5). For each of these categories, we also found interneuron clusters that were significantly inhibited by the CS. Since amygdala interneurons are tonically active *in vivo*, leading to persistent inhibition of downstream PNs or interneurons^14,18,24^, suppression of their activity after learning can induce selective disinhibition, allowing for circuit computations necessary for learning and memory expression. This inhibitory plasticity can for example stabilise memory traces or increase the selectivity of engrams^32^, or enhance the contrast between distinct long-range PN circuits associated with fear or extinction states^22,33^.

Some of these activity patterns were enriched in SST and VIP interneurons during distinct auditory cues or in discrete behavioural states. VIP interneurons were overall more excited by novel auditory cues during habituation, and displayed overall stronger CS+ responses compared to SST interneurons during the progression of fear learning. VIP interneurons showed a significantly larger fraction of stably activated neurons and more upregulation of CS+ responses during conditioning, while SST interneurons had a comparably higher fraction of CS+ inhibited cells. This is consistent with the notion that PN dendritic disinhibition increases auditory processing necessary for associative fear learning^14,15,18^. Increased activity in VIP interneurons, which enables such disinhibition, might facilitate the processing of novel and salient information in microcircuits, as recently also shown in other brain areas^34–36^. In the amygdala, elevated VIP activity in high fear states could similarly enable the integration of context-dependent information or promote higher-order conditioning. In contrast, SST interneurons were predominantly activated in low fear states after extinction, which was recently also reported for hippocampal SST cells^37^. Our findings that SST interneurons preferentially signal safety in fear learning is consistent with previous reports, showing increased BLA SST interneuron activity to learned non-threatening cues^17^. In associative fear conditioning, increased activity of dendrite-targeting SST interneurons would perturb processing of inputs at PN dendrites and thus dampen PN responsiveness to auditory cues, which would represent a mechanism to suppress fear neuron activity^31^.

### Mechanisms of interneuron plasticity

The question remains as to where this plasticity is located – are interneurons and their synapses indeed undergoing plastic changes themselves, or are changes in activity patterns across learning and extinction simply imposed by synaptic inputs? Both local PNs^15,25,28^ and long-range excitatory afferents e.g. from auditory thalamus^30^ display similar activity patterns in associative learning as observed here for amygdala interneurons. Since they have been shown to impinge on several BLA interneuron subpopulations^18,23,38^, auditory thalamus or local PN inputs could relay plastic activity patterns, which would additionally lead to feed-forward inhibition of other interneurons within the interconnected BLA microcircuits.

In addition, there is ample evidence from biochemical, electrophysiological and anatomical *ex vivo* studies to support the idea of local plasticity within BLA interneurons. For example, aversive learning alters the expression of enzymes for GABA synthesis, GABA receptors and their scaffolding protein gephyrin in the BLA^3–5^. Glutamatergic inputs to amygdala interneurons can be potentiated by tetanic stimulation, which leads to an increase of inhibitory synaptic drive onto PNs^6–8^. Conversely, fear learning can reduce excitatory inputs onto distinct subpopulations of lateral amygdala interneurons such as PV cells^20^. Together, these *in vitro* data point to a bidirectional regulation of interneuron plasticity upon associative learning, which is in line with our observations of cells that are selectively activated or inhibited in high fear states. Furthermore, fear conditioning has been shown to induce structural remodelling of GABAergic synapses that can be reversed by extinction^39^. However, extinction does not simply reverse conditioning-induced potentiation^39,40^ which would resemble a process of forgetting. Considered to be a new form of context-dependent safety learning that suppresses the original fear memory^41^, extinction has additionally been associated with potentiation of excitatory synapses on inhibitory interneurons, but also GABAergic synapses on PNs^10,12^. This is reflected in extinction-mediated up- and downregulation of CS responses in distinct subsets of interneurons in the present study. Altogether, our *in vivo* results are therefore consistent with previous *ex vivo* work on cellular mechanisms of associative fear learning and extinction.

Our data offer additional insights into behaviourally-mediated plasticity of individual inhibitory BLA neurons. Interneurons displayed mixed selectivity during learning, with cells encoding the CS+, CS– and US being spatially intermingled in the BLA (Fig. 2 and Fig. 6C, D). A large proportion of interneurons was activated by both the predictive CS+ and the instructive US, making them ideal candidates for cellular plasticity during Hebbian learning. However, upregulation of CS+ responses during fear learning (Fig. 4) or expression (Fig. 5) was independent of US activation during conditioning, suggesting that somatic CS-US coincidence is not necessary for learning-associated adaptation of interneuron activity. Yet, our somatic recordings cannot capture converging CS and US inputs in dendrites, which could induce dendritic plasticity in interneurons^42–44^ and thus represent a potential source of CS response enhancement. Moreover, similar proportions of interneurons down-regulated their CS response upon fear learning, including cells that were significantly activated by the US. This supports the idea that memory processes involve diverse forms of plasticity beyond classical Hebbian learning in interneurons, as previously proposed for amygdala PNs^15,28^.

### Ensemble coding in amygdala interneurons

At the population level, we found that CS+, CS– and baseline could be reliably decoded from interneuron activity within each experimental day, but not across behavioural states (Fig. 6). This suggests that CS information is stably encoded within the ensemble, although the activity of individual interneurons changes dynamically from day to day during learning and extinction. Decoding accuracy was highest when all interneurons or only ‘CS responsive’ cells were included in the sample, while decoders trained on ‘non-CS responsive’ interneurons exhibited reduced accuracy (although, still higher than randomly shuffled data). This might be due our strict definition of a responsive neuron, which is based on at least three significant responses during four CS+ or CS– presentations, respectively. The higher-than-random accuracy for ‘non-CS responsive’ interneurons suggests that these cells indeed carry valuable information about the auditory cues that eludes our strict definition of ‘responsive’.

Using population vector distance analysis, we could further observe that the representation of both the CS+ and the CS– was getting more similar to the aversive US during learning. In contrast to BLA PNs, where this effect is selective for CS+ encoding and mainly mediated by up- and downregulation of CS responses^28^, we found that defined individual CS+ clusters of interneurons contributed little to this effect. Instead, the reduction in the population vector distance between CS+ and the aversive US signal seems to be driven by interneurons across all clusters, yet can be significantly affected by the ‘Activated stable’ US cluster. This effect might be driven by VIP interneurons, since this cluster is overrepresented in this interneuron subpopulation (Fig. 7). Moreover, unlike the plasticity at the single neuron level, learning-induced changes at the population level during conditioning were transient and did not persist over days. This suggests that although representations of sensory stimuli change during fear learning, unlike PNs^28^, this transient shift does not consolidate overnight, which might ensure valence-free representations of environmental cues within the BLA interneuron population. Additional studies will be needed to address how individual interneuron subpopulations contribute to these high dimensional representations.

### Implications of functional interneuron diversity

Remarkably, across days and stimuli, amygdala interneurons displayed high diversity in their response patterns, even within a molecular subpopulation. Population averages were dominated by activated neurons even when similar proportions of cells were classified as activated or inhibited (see e.g. Fig. 3). This is likely due to asymmetric effects of suppression or increase of neuronal spiking, which will strongly depend on the baseline firing rate of any given neuron, but also individual differences in calcium buffering as well as non-linearity of GCaMP indicators^45^. Together, these results highlight that any interpretation that a molecularly-defined interneuron group is homogeneously activated based on population averages (e.g. fibre-photometry recordings) should be made with caution. Similarly, opto- or chemogenetic manipulations that uniformly drive the entire population of any given interneuron subpopulation will artificially overrule the physiologically diverse response patterns during behaviour^46^.

But how can one make sense of this functional diversity of interneurons? Non-uniform responses across a neuronal population of amygdala neurons have behavioural advantages, as they enable for example avoidance of generalisation by selective and precise computations to specific environmental cues, and allows for circuit adaptations in dependence of varying internal states^25,47,48^. However, the broad molecular subpopulations currently used to classify interneurons obscure the diverse molecular, morphological, physiological and functional characteristics of individual cells. A true definition of an interneuron subtype must be based on its functional properties and connectivity, i.e., its role in a neural circuit. Molecular classification provides genetic entry points to interneuron targeting and might correlate with other cellular properties, but the currently employed genetic mouse lines, such as *SST-Cre* or *VIP-Cre*, might not be sufficient to account for the reported diversity in morphology, connectivity and function at the single cell level^14,18,23^. Indeed, a recent study on cortical SST interneurons has shown selective morphology and connectivity of SST molecular subclasses^49^. In the BLA, VIP interneurons targeting other interneurons vs. those connecting to PNs can be identified by co-expression patterns with cholecystokinin (CCK)^19^. Recent advances in single-cell RNA sequencing now enable an even finer grained interneuron classification in the BLA^50–52^, allowing to target further subclasses with intersectional genetic approaches. Yet, these are limited to two or three genes^53^, and will greatly reduce cell numbers that can be recorded simultaneously and thus affect the interpretability of results. In the future, novel high-plex spatial transcriptomics after unbiased *in vivo* functional recordings^54–56^ can help to determine whether discrete molecular and anatomical characteristics of interneurons also correlate with distinct functional properties.

## METHODS

### Animals

All animal procedures were performed in accordance with institutional guidelines at the Friedrich Miescher Institute for Biomedical Research and were approved by the Veterinary Department of the Canton of Basel-Stadt. Heterozygous (cre/wt) *GAD2-Ires-Cre, VIP-Ires-Cre* and *SST-Ires-Cre* mice^27^ fully backcrossed to a C57BL/6J background were used for virally mediated, Cre-dependent expression of a calcium indicator. Experiments were performed with male (*GAD2-Cre, VIP-Cre, SST-Cre*) and female (*GAD2-Cre*) mice aged 2-3 months at the time of injection. Animals were kept in a 12 h light/dark cycle with access to food and water *ad libitum* and were individually housed after implant surgeries. All experiments were conducted during the light cycle. Of note, a different analysis on fear conditioning day data from *VIP-Cre* and *SST-Cre* mice of this study was previously published^18^.

### Surgical procedures

Surgical procedures were performed as previously described^18,25^. In brief, mice were anaesthetised using isoflurane (3-5% for induction, 1-2% for maintenance; Attane, Provet) in oxygen-enriched air (Oxymat 3, Weinmann) and fixed on a stereotactic frame (Model 1900, Kopf Instruments). Injections of buprenorphine (Temgesic, Indivior UK Limited; 0.1 mg/kg body weight subcutaneously 30 min prior to anaesthesia) and ropivacaine (Naropin, AstraZeneca; 0.1 ml locally under the scalp prior to incision) were provided for analgesia. Postoperative pain medication included buprenorphine (0.1 mg/kg in the drinking water; overnight) and injections of meloxicam (Metacam, Boehringer Ingelheim; 1 mg/kg subcutaneously) for up to three days if necessary. Ophthalmic ointment (Viscotears, Bausch & Lomb) was applied to avoid eye drying. Body temperature of the experimental animal was maintained at 36 °C using a feedback-controlled heating pad (FHC). AAV2/9.CAG.flex.GCaMP6f or AAV2/9.CAG.flex.GCaMP6s^26^ (400 nl, University of Pennsylvania Vector Core, UPenn) was unilaterally injected into the BLA using a precision micropositioner (Model 2650, Kopf Instruments) and pulled glass pipettes (tip diameter about 20 µm) connected to a Picospritzer III microinjection system (Parker Hannifin Corporation) at the following coordinates from bregma: AP -1.5 mm, ML -3.3 mm, DV 4.1-4.5 mm below the cortical surface. The skin incision was closed with polypropylene suture (Prolene 6-0, Ethicon) and the animal placed into a recovery cage on a heating pad until fully mobile. Two weeks after virus injection, a gradient-index microendoscope (GRIN lens, 0.6 x 7.3 mm, GLP-0673, Inscopix) was implanted into the BLA using the same surgical approach. A sterile needle (0.7 mm diameter) was used to make an incision above the implant site. The GRIN lens was subsequently lowered into the brain with a micropositioner (coordinates from bregma: AP -1.6 mm, ML -3.2 mm, DV 4.5 mm below the cortical surface) using a custom-build lens holder and fixed to the skull using UV light-curable glue (Loctite 4305, Henkel). The skull was sealed with Scotchbond (3M), Vetbond (3M) and finally dental acrylic (Paladur, Heraeus). A custom-made head bar for animal fixation during the miniature microscope mounting procedure was embedded into the dental cement. Mice were allowed to recover for at least one week after GRIN lens implantation before starting to check for GCaMP expression.

### Deep-brain calcium imaging

Starting one week after GRIN lens implantation, mice were head-fixed to check for sufficient expression of GCaMP using a miniature microscope (nVista HD, Inscopix). Two to four weeks after the implant surgery, mice were briefly anaesthetised with isoflurane to fix the microscope baseplate (BLP-2, Inscopix) to the skull using light-curable composite (Vertise Flow, Kerr). The microscope was removed, and the baseplate capped with a baseplate cover (Inscopix) whenever the animal was returned to its home cage. The microscope was mounted daily immediately before starting the behavioural session. Mice were habituated to the brief head-fixation on a running wheel for miniature microscope mounting for at least three days before the behavioural paradigm. Imaging data was acquired using nVista HD software (Inscopix) at a frame rate of 20 Hz with an LED power of 40-80% (0.9-1.7 mW at the objective, 475 nm), analogue gain of 1-2 and a field of view of 650 x 650 µm. For individual mice, the same imaging parameters were kept across repeated behavioural sessions.

### Behaviour

Two different contexts were used for the associative fear learning paradigm. Context A (retrieval context) consisted of a clear cylindrical chamber (diameter: 23 cm) with a smooth floor, placed into a dark-walled sound attenuating chamber under dim light conditions (approximately 25 lux). The chamber was cleaned with 1% acetic acid. Context B (fear conditioning context) contained a clear square chamber (26 x 26 cm) with an electrical grid floor (Coulbourn Instruments) for foot shock delivery, placed into a light-coloured sound attenuating chamber with bright light conditions (approximately 180 lux), and was cleaned with 70% ethanol. Both chambers contained overhead speakers for delivery of auditory stimuli, which were generated using a System 3 RP2.1 real time processor and SA1 stereo amplifier with RPvdsEx software (all Tucker-Davis Technologies). A precision animal shocker (H13-15, Coulbourn Instruments) was used for the delivery of alternating current (AC) foot shocks through the grid floor. Behavioural protocols for stimulus control were generated with Radiant Software (Plexon) via TTL pulses. On day 1, mice were habituated in context A. Two different pure tones (conditioned stimulus, CS; 6 kHz and 12 kHz, total duration of 30 s, consisting of 200 ms pips repeated at 0.9 Hz; 75 dB sound pressure level) were presented five times each in an alternated fashion with a pseudorandom ITI (range 60-90 s, 2 min baseline before first CS). On day 2, mice were conditioned in context B to one of the pure tones (CS+) by pairing it with an unconditioned stimulus (US; 2 s foot shock, 0.65 mA AC; applied after the CS at the time of next expected pip occurrence). The other pure tone was used as a CS– and not paired with a US. CS+ with US and CS– were presented alternating five times each in a pseudorandom fashion (ITI 60-90 s), starting with the CS+ after a 2 min baseline period. Animals remained in the context for 1 min after the last CS– presentation and were then returned to their home cage. On day 3 and 4, fear memory was tested, and extinction induced in context A. After a 2 min baseline period, the CS– was presented four times, followed by 12 CS+ presentations (ITI 60-90 s). The use of 6 kHz and 12 kHz as CS+ was counterbalanced across animals. Timestamps of calcium imaging frames, behavioural videos and external stimuli were collected for alignment on a master clock using the MAP data acquisition system (Plexon).

### Histology

Mice were deeply anaesthetized with urethane (2 g/kg body weight; intraperitoneally) and transcardially perfused with 0.9% NaCl followed by 4% paraformaldehyde in PBS. The GRIN lens was removed and brains post-fixed in 4% paraformaldehyde for at least 2 h at 4 °C. Coronal sections (120 µm) containing the BLA were cut with a vibratome (VT1000S), immediately mounted on glass slides and cover-slipped using Vectashield (Vector Laboratories). To verify the GRIN lens position, sections were scanned with a laser scanning confocal microscope (LSM700, Carl Zeiss AG) equipped with a 10x air objective (Plan-Apochromat 10x/0.45) and matched against the Allen Mouse Brain Reference Atlas (https://mouse.brain-map.org).

A subset of animals was used for immunohistochemical analysis of interneuron marker gene expression in GCaMP6f+ neurons of *GAD2-cre* mice. Here, brains were cut into 80 µm coronal slices with a vibratome. Sections were washed in PBS four times and blocked in 10% normal horse serum (NHS, Vector Laboratories) and 0.5% Triton X-100 (Sigma-Aldrich) in PBS for 2 h at room temperature. Slices were subsequently incubated in a combination of the following primary antibodies in carrier solution (1% NHS, 0.5% Triton X-100 in PBS) for 48 h at 4 °C: rabbit anti-VIP (1:1000, Immunostar, 20077, LOT# 1339001), rat anti-SST (1:500, Merck Millipore, MAB354, LOT# 232625), guinea pig anti-PV (1:500, Synaptic Systems, 195004, LOT# 195004/10). Separate sections were stained using rabbit anti-pro-CCK (1:500, Frontiers Institute, CCK-pro-Rb-Af350, LOT# N/A). After washing three times with 0.1% Triton X-100 in PBS, sections were incubated for 12-24 h at 4 °C with a combination of the following secondary antibodies in carrier solution: goat anti-rabbit Alexa Fluor 647 (1:750, Thermo Fisher Scientific, A21245, Lot# 1778005), goat anti-rat Alexa Fluor 568 (1:750, Thermo Fisher Scientific, A11077, Lot# 692966), goat anti-guinea pig DyLight 405 (1:250, Jackson ImmunoResearch, 106-475-003, Lot# 126016). After washing four times in PBS, sections were mounted on glass slides and cover-slipped with Vectashield. Sections were scanned using a laser scanning confocal microscope (LSM700) equipped with a 10x air objective (Plan-Apochromat 10x/0.45) or 20x air objective (Plan-Apochromat 20x/0.8). Tiled z-stacks (3 µm step size) of the BLA were acquired and stitched with Zeiss software processing tool (ZEN 2.3, black edition, Carl Zeiss AG).

### Calcium imaging analysis

#### Pre-processing

Raw image data was analysed as previously described^18,25^. In brief, videos were spatially down sampled (4x), bandpass filtered (Fourier transform) and normalized by the filtered image in ImageJ. The movies from all the sessions of each animal were concatenated into a single file and motion-corrected with the non-rigid algorithm NormCorre using the CaImAn package^57^. Cell detection was carried out using Principal Component Analysis (PCA) and Independent Component Analysis (ICA) with the CIAtah package^58^. ROIs were initially oversampled and then manually inspected. ROIs not matching individual neurons in the motion-corrected video were removed. The selected ROIs were then used to extract the raw fluorescence traces using CIAtah based on the 20 Hz video. Fluorescence traces were z-scored and binned in 250 ms for all further analyses. *Data analysis*. To analyse the responses around events (e.g., tone presentation), z-scored traces were baselined to the period before the event. For CS presentations, the baseline was set to 30 s and for US presentations to 5 s. Then, to detect statistically responsive interneurons, we compared the fluorescence trace during the baseline period to the trace evoked during the event using the non-parametric Wilcoxon rank-sum test. Only cells that displayed statistically significant responses (*p* < 0.01) to at least 3 CS or US presentations were classified as ‘responsive’. For across-day comparisons, the first four CS presentations during habituation were compared to the four CS– presentations during day 3 (test/extinction 1) and 4 (extinction 2), and the first and last four CS+ presentations for test and extinction sessions, respectively. This was done to compensate for the uneven number of CS+ and CS− presentations across habituation and test/extinction days. To identify subclasses of responses, PCAs with K-means clustering were performed on the baselined traces of significantly responsive interneurons (using an explained variance of at least 80%). The number of clusters generated by K-means were initially set to 16 for US response patterns and 8 for CS+ and CS– response patterns during conditioning. For across days, they were set to 10 for both CS+ and CS– response patterns. All activity clusters were visually inspected, classified, and merged if necessary. For *VIP-Cre* mice, N = 2 animals were excluded from the conditioning day analysis as one of the five US exposures could not be verified with the behavioural recordings. However, these mice were included for across day analysis, since they learned the association between CS+ and US with the remaining 4 pairings.

#### Population analysis

To measure the similarity between two sets of neuronal ensemble response patterns, we calculated the population vector distance (PVD) between activity vectors as described in Taylor et al. (2021)^30^. The distance between CS and US activity vectors was calculated using 30 s binned responses to CS presentations and the mean 5 s response to the US. Activity vectors were of length *n* interneurons, for example, a population vector for the US presentation was created based on the mean response of each interneuron to the foot shocks (e.g., for an animal with 40 cells, the vector would have 40 mean fluorescence values). The Euclidean distance between each CS bin vector and the mean US response vector was then calculated and averaged for each CS presentation. During fear conditioning, the change in distance between CS and US vectors was normalised to the PVD of the first CS/US pairing in the session. Negative percentages indicate that CS and US activity patterns are becoming more similar, while positive percentages indicate they are diverging. For the fear conditioning session, we furthermore calculated the difference between the mean change in PVD in the early stage of conditioning (pairings 1-2) and the late stage of conditioning (pairings 4-5). For across-days PVD analysis, the PVD in each day was the result of averaging the PVD from the first 4 CS presentations to the mean US vector from conditioning, and the change in PVD was normalised to the distance between CS and US during habituation day.

#### Decoders

The calcium traces to the CS presentations and baselines were used to train binary linear support vector machines (SVM) using MATLAB built-in functions *fitcsvm* and *fitcecoc*. To account for the fact that animals had different numbers of cells, we randomly selected 37 cells from each animal, given that this was the minimum number of cells detected in one animal, and report the mean results from 100 independent iterations. To balance the data, we used the same amount of data for each CS and baseline period, each with 30 s windows binned in 1 s. Intraday decoders were validated using a 10-fold cross-validation procedure using MATLAB function *crossval*, in which decoders are trained on a partition of 90% of the data and tested on the remaining 10%, this procedure is carried out 10 times. Within each day, we trained a multiclass decoder to decode baseline vs. CS+ vs. CS– using different subsets of interneurons (e.g., only CS responsive or only non-CS responsive). As a control, we trained SVMs on temporally shuffled CS+, CS- and baseline labels. Additionally, to rule out the possibility that differences in non-CS decoder accuracy were due to fewer available cells, we trained control decoders on a random sample taken from all interneurons, matching the number of cells used in the non-CS decoders.

To compare how CS+ and CS– coding changed across days, we trained separately two-way decoders (baseline vs. CS+ and baseline vs. CS–) and used them to predict the stimulus identity during the other sessions. Overall accuracy across days was calculated as the sum of the diagonal of the confusion matrix (i.e., the number of true positives) divided by the overall sum of the confusion matrix. Using the confusion matrix, we also report precision (i.e. True positives/(True positives + False positives)), recall (i.e. True positives/(True positives + False negatives)) and F1 score (2 * (precision * recall) / (precision + recall)) for each class decoded.

Finally, we used decoder weights as a proxy to indicate the contribution of each interneuron to distinguishing between the trained classes (baseline, CS+ and CS–). Thus, to investigate interneuron selectivity for CS+ or CS–, we obtained the corresponding absolute decoding weights from the baseline vs. CS+ and baseline vs. CS– two-way decoders for each interneuron and calculated the correlation between them. A negative correlation between the weights would suggest stimulus selectivity^29^. We report the average correlation of each animal from 100 iterations. Furthermore, as a control, we calculated the correlation between CS+ decoding weights at the *ith* and *ith+1* iteration of decoders trained on a randomly selected partition of 90% of the data from all interneurons. Average correlation between CS+ decoding weight at the *ith* and *ith+1* iterations were calculated per animal (N = 9 mice), using a total of 200 iterations to obtain 100 correlation values.

### Behaviour analysis

Pose estimation was performed using DeepLabCut version 2.2.2^59^. A ResNet-50-based neural network^60^ was employed, with the network running for 1,060,000 training iterations. For training, locations of eight mouse body parts were manually labelled in video frames: nose, base of miniature microscope, left ear base, right ear base, neck, upper spine, middle spine, and tail base. A total of 146 frames were labelled from eight videos, which included both behavioural contexts and lighting conditions. Of these labelled frames, 95% were used for training. Post-processing of DeepLabCut output files, which included x/y coordinates and likelihood values for each body part, involved filtering low likelihood positions. These positions were replaced with the most recent highly probable positions. The displacement of each point over time was then calculated. Periods of pause were identified based on the displacement thresholds of five tracked points: left ear base, right ear base, upper spine, middle spine, and tail base. All points had to meet a threshold for displacement (0.7-3.5) to be considered in the analysis. Immobility was defined as a period during where the animal did not move its body for at least 2 seconds. The automated behavioural scoring was subsequently validated by a human scorer to ensure accuracy and reliability of the pose estimation results.

## Statistical analysis and data presentation

The number of analysed cells is indicated with ‘n’, while ‘N’ declares the number of animals. Averaging across multiple trials per cell/animal is indicated in the figure legends and respective methods sections where applicable. Reported n/N numbers always refer to data from individual cells/animals, no samples were measured repeatedly for statistical analysis. No statistical methods were used to predetermine sample sizes.

Statistical analysis was carried out using R, Matlab or Prism 10 (GraphPad Software). All datasets were tested for Gaussian distribution using the Shapiro Wilk test. Equal variance across samples was assessed with Levene’s test. Overall, the data did not pass normality or homoscedasticity tests, thus we report only non-parametric tests results. For comparisons of multiple groups in a repeated measures variable, we used a Friedman test followed by Dunn’s multiple comparisons. Comparisons of individual cellular responses to specific stimuli were analysed with paired Wilcoxon tests, comparisons between subpopulations with Mann-Whitney tests. In both cases, *p* values were adjusted with Bonferroni correction. To assess whether the distribution of cells in defined activity patterns was significantly different between stimuli or interneuron subpopulations, we applied Chi-Square tests. If significant, proportions were compared in a pairwise manner using the R function *prop*.*test*, which calculates a Chi-Square with Yates continuity correction for small expected n numbers (n < 5). Statistical significance threshold was set at 0.05 and significance levels are presented as * (*p* < 0.05), ** (*p* < 0.01) or *** (*p* < 0.001) in all figures. Statistical tests and results are reported in the respective figure legends, as well as in Supplementary Table 1.

Contrast and brightness of representative example images were minimally adjusted using ImageJ. For figure display, confocal images were further scaled (0.5×0.5), and calcium activity was resampled to 4 Hz (traces and heatmaps). Averaged traces are displayed as mean with s.e.m. Violin plots illustrate distribution of all data points. Tukey box-and-whisker plots show median values, 25^th^ and 75^th^ percentiles, and min to max whiskers with exception of outliers (beyond 1.5 times interquartile range).

## Data and code availability

Custom code and full datasets will be made available upon final publication and are currently available upon request from SK.

## Author contributions

Conceptualization: YB, JG, AL, SK; Methodology: NF, JCM, YB, JG, SK; Investigation: NF, JCM, JG, SK; Analysis: NF, JCM, BE, CP, YB, JG, SK; Visualization: NF, JCM, SK; Funding acquisition: AL, JG, SK; Supervision: SK, AL; Writing – original draft: NF, SK; Writing – review & editing: all authors.

## Acknowledgements

The authors thank all members of the Krabbe lab for helpful discussions and comments. They thank Olga Sharma, Tobias Eichlisberger, Christian Müller and all staff of the FMI and DZNE Animal Facilities for excellent technical assistance. They further thank the Facility for Imaging and Microscopy at the FMI, and the Light Microscopy Facility at the DZNE for their support with data acquisition and analysis, and Isaac Samuel Racine for advice on statistics. They are grateful to the GENIE Program at Janelia Research Campus of the Howard Hughes Medical Institute for making GCaMP6 available.

## Funding

Deutsches Zentrum für Neurodegenerative Erkrankungen (DZNE), Bonn (SK, JG); Friedrich Miescher Institute for Biomedical Research (FMI), Basel (AL); Chan Zuckerberg Initiative, Ben Barres Early Career Acceleration Award (SK); Brain & Behavior Research Foundation, Young Investigator Award (SK); Dementia Research Switzerland – Synapsis Foundation, Career Development Award (SK); Deutsche Forschungsgemeinschaft (DFG), SFB 1089 Teilprojekt (SK, JG) and SPP 2411 Teilprojekt (JG); iBehave Network – sponsored by the Ministry of Culture and Science of the State of North Rhine-Westphalia (SK, JG); Swiss National Science Foundation (SNSF) Grant 310030_189123 and Adv Grant TMAG-3_209270 (AL); European Research Council, Starting Grant, AXPLAST (JG). The funders had no role in study design, data collection and analysis, decision to publish or preparation of the manuscript.

## Supplementary Materials

Supplementary Figures 1-9

Supplementary Table 1

---

**Supplementary Figure 1:**
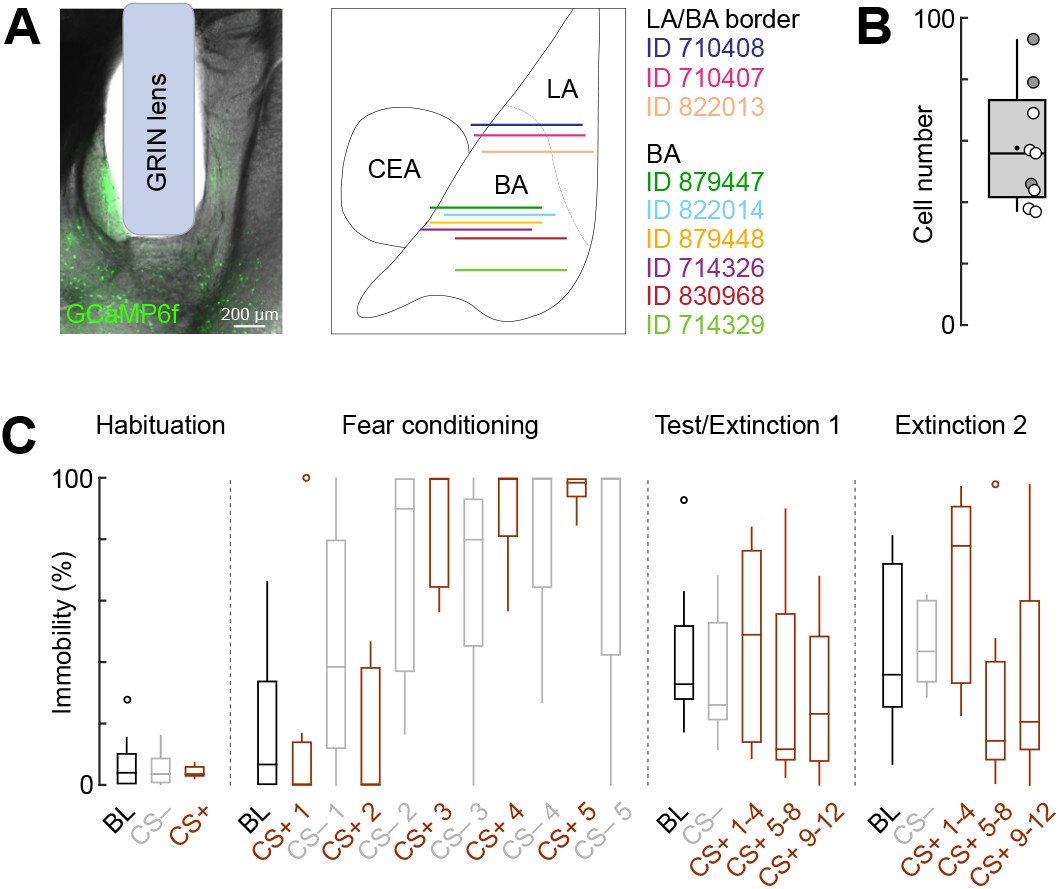
Imaging of basolateral amygdala interneurons during fear and extinction learning. **A**, Representative implant site (ID 879448) and schematic illustrating all reconstructed implant sites of GRIN lenses (lens front) within the BLA of *GAD2-Cre* mice for deep brain imaging experiments matched to a mouse brain atlas (N = 9 mice). LA, lateral amygdala; BA, basal amygdala; CEA, central amygdala. **B**, Average cell numbers recorded across the four-day paradigm (N = 9). Tukey box-and-whisker plot illustrates median values, 25^th^ and 75^th^ percentiles, and min to max whiskers, dot indicates the mean. Circles represent individual animals (open circles, imaging sites in the basal amygdala (N = 6); filled circles, at the border of the lateral and basal amygdala (N = 3). **C**, Immobility levels throughout the fear conditioning and extinction paradigm in GRIN lens-implanted *GAD2-Cre* mice (N = 9). Tukey box-and-whisker plots illustrates median values, 25^th^ and 75^th^ percentiles, and min to max whiskers, circles indicate outliers. Favila, Capece Marsico et al. 2024 – Supplementary Figure 2

**Supplementary Figure 2:**
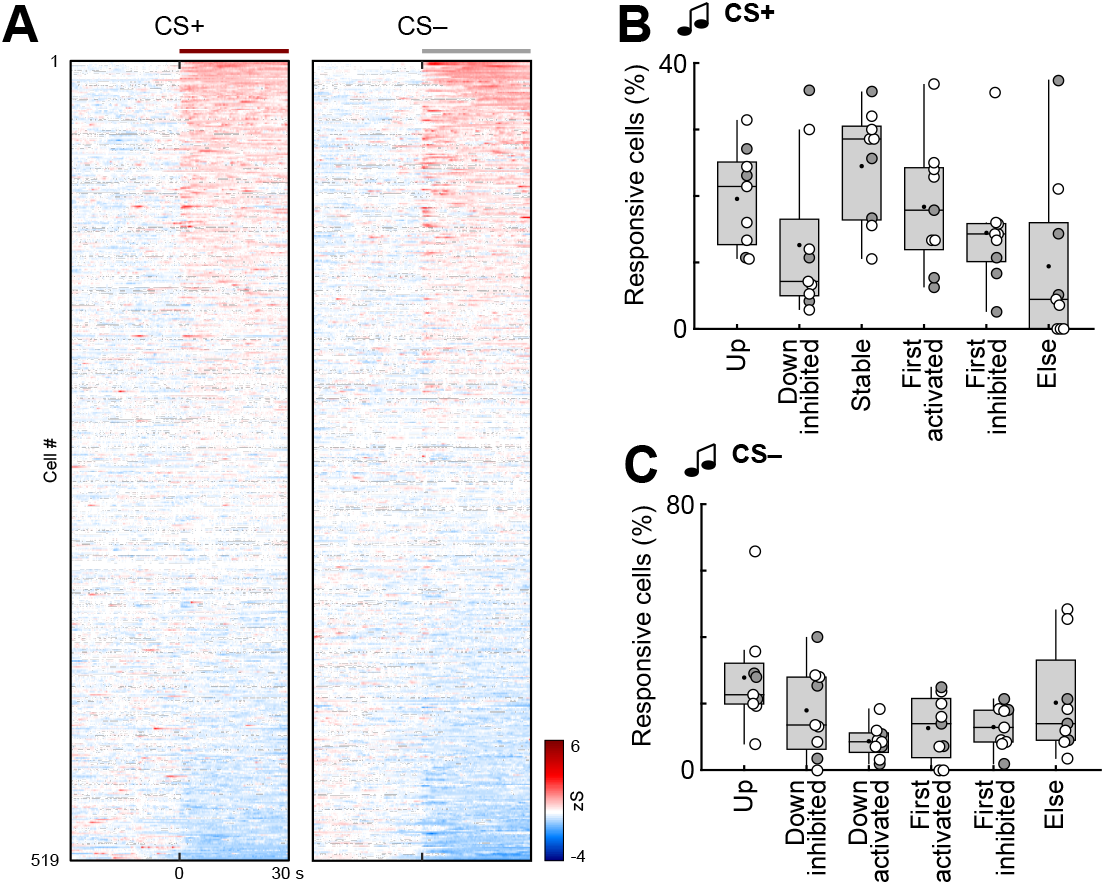
CS coding in amygdala interneurons during fear conditioning. **A**, Heatmap of CS+ and CS– responses in BLA interneurons during conditioning, averaged across all five trials and ordered individually by response amplitude (n = 519 cells from N = 9 mice). Lines indicate CS duration. **B**, Fraction of interneurons according to CS+ cluster membership across animals (see Figure 4; N = 9). Friedman test (χ2 = 11.26), *p* = 0.0465; followed by Dunn’s multiple comparisons (non-significant). **C**, Fraction of interneurons according to CS– cluster membership across animals (see Figure 4; N = 9). Tukey box-and-whisker plots in B and C show median values, 25^th^ and 75^th^ percentiles, and min to max whiskers with exception of outliers, dots indicate the mean. Circles represent individual animals (open circles, imaging sites in the basal amygdala (N = 6); filled circles, at the border of the lateral and basal amygdala (N = 3). Additional details of statistical analyses are provided in Supplementary Table 1. Favila, Capece Marsico et al. 2024 – Supplementary Figure 3

**Supplementary Figure 3:**
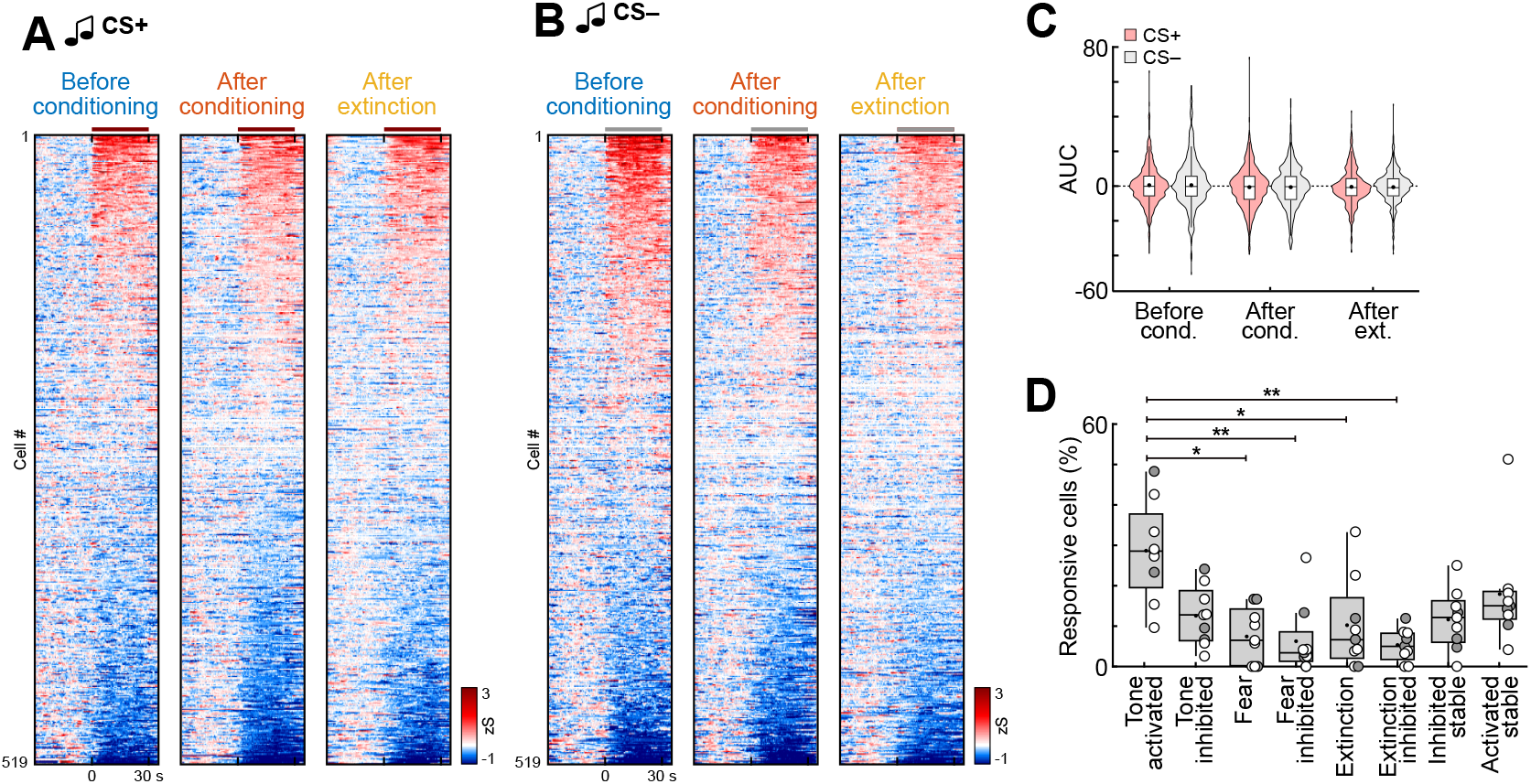
CS encoding across learning days in basolateral amygdala interneurons. **A**, Heatmap of CS+ and **B**, CS– responses in BLA interneurons before conditioning, after conditioning and after extinction, averaged across the four CS presentations used for clustering (n = 519 cells from N = 9 mice). Line indicates CS duration. **C**, Comparison of area under the curve (AUC) for CS+ and CS– responses in BLA interneurons (n = 519). **D**, Fraction of interneurons according to CS– cluster membership across animals (see Figure 5; N = 9). Friedman test (χ2 = 25.22), *p* = 0.0007, followed by Dunn’s multiple comparisons (‘Tone activated’/activated before conditioning vs. ‘Fear’/activated after conditioning, *p* = 0.0494; ‘Tone activated’/activated before conditioning vs. ‘Fear inhibited’/inhibited after conditioning, *p* = 0.0072; ‘Tone activated’/activated before conditioning vs. ‘Extinction’/activated after extinction, *p* = 0.0212; ‘Tone activated’/activated before conditioning vs. ‘Extinction inhibited’/inhibited after extinction, *p* = 0.0022). Violin plots in C show distribution of all data points, Tukey box-and-whisker plots in C and D show median values, 25^th^ and 75^th^ percentiles, and min to max whiskers with exception of outliers, dots indicate the mean. Circles in D represent individual animals (open circles, imaging sites in the basal amygdala (N = 6); filled circles, at the border of the lateral and basal amygdala (N = 3). \**p* < 0.05, \*\**p* < 0.01. Additional details of statistical analyses are provided in Supplementary Table 1.

**Supplementary Figure 4:**
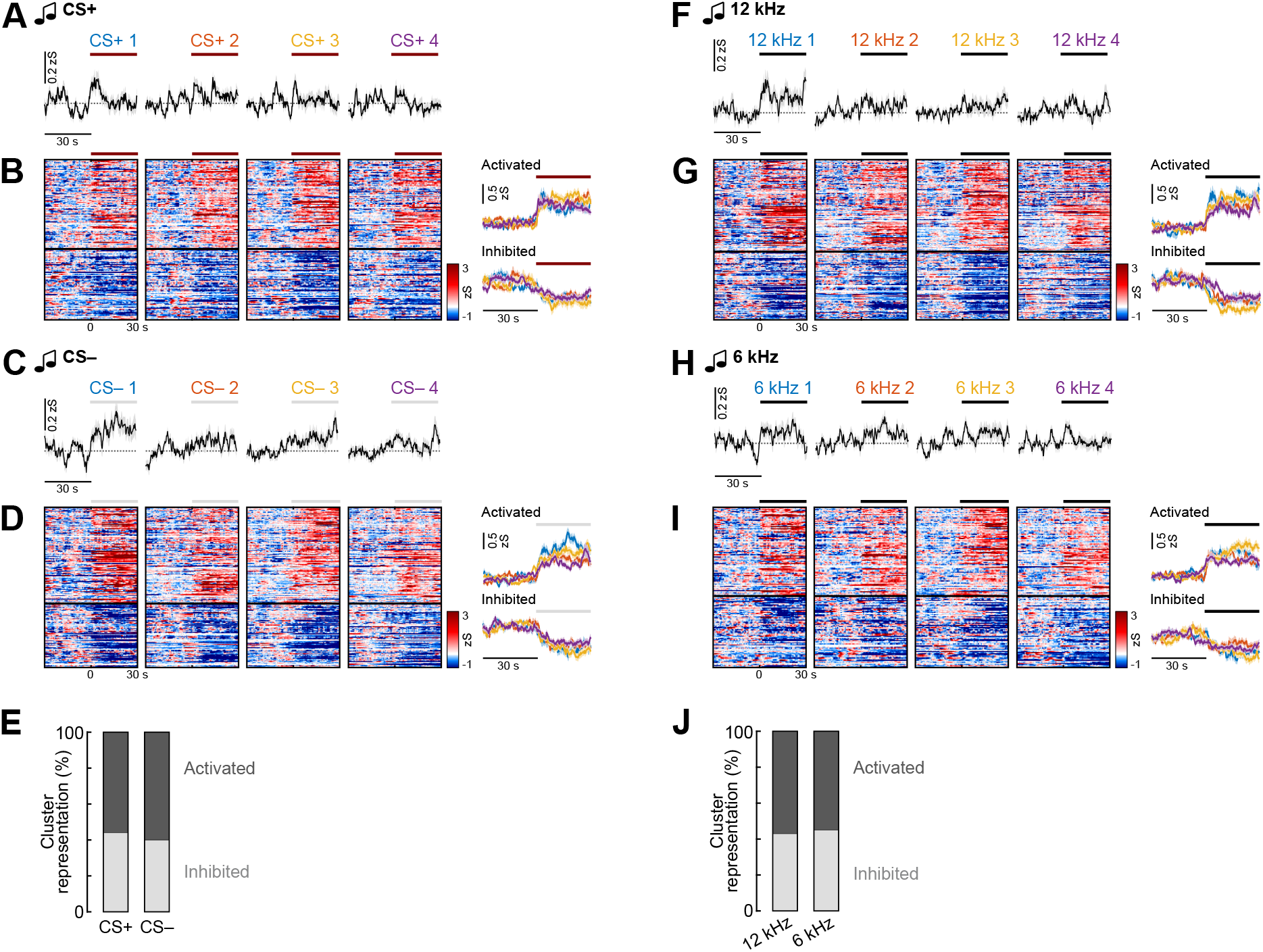
Interneuron responses to auditory stimuli. **A**, Average traces of basolateral amygdala interneurons during the first four CS+ presentations during habituation (n = 519 cells from N = 9 mice; counterbalanced for 6 kHz and 12 kHz). Line indicates CS duration. **B**, Heatmap (left) of CS+ responses clustered into groups depending on their response pattern across the four presentations and corresponding average traces of clusters (right). n = 165 responsive cells; ‘Activated’, n = 92; ‘Inhibited’, n = 73. **C**, Average traces of basolateral amygdala interneurons during the first four CS– presentations during habituation (n = 519; counterbalanced for 6 kHz and 12 kHz). **D**, Heatmap (left) of CS– responses clustered into groups depending on their response pattern across the four presentations and corresponding average traces of clusters (right). n = 185 responsive cells; ‘Activated’, n = 111; ‘Inhibited’, n = 74. **E**, Proportion of cells in CS+ and CS– clusters (CS+, n = 165; CS–, n = 185). **F**, Average traces of basolateral amygdala interneurons during the first four 12 kHz presentations during habituation (n = 519; later assigned to be CS+ or CS–). **G**, Heatmap (left) of 12 kHz responses clustered into groups depending on their response pattern across the four presentations and corresponding average traces of clusters (right). n = 202 responsive cells; ‘Activated’, n = 116; ‘Inhibited’, n = 86. **H**, Average traces of basolateral amygdala interneurons during the first four 6 kHz presentations during habituation (n = 519; later assigned to be CS+ or CS–). **I**, Heatmap (left) of 6 kHz responses clustered into groups depending on their response pattern across the four presentations and corresponding average traces of clusters (right). n = 148 responsive cells; ‘Activated’, n = 82; ‘Inhibited’, n = 66. **J**, Proportion of cells in 12 kHz and 6 kHz clusters (12 kHz, n = 202; 6 kHz, n = 148). Average traces across panels are mean with s.e.m.. Additional details of statistical analyses are provided in Supplementary Table 1. Favila, Capece Marsico et al. 2024 – Supplementary Figure 5

**Supplementary Figure 5:**
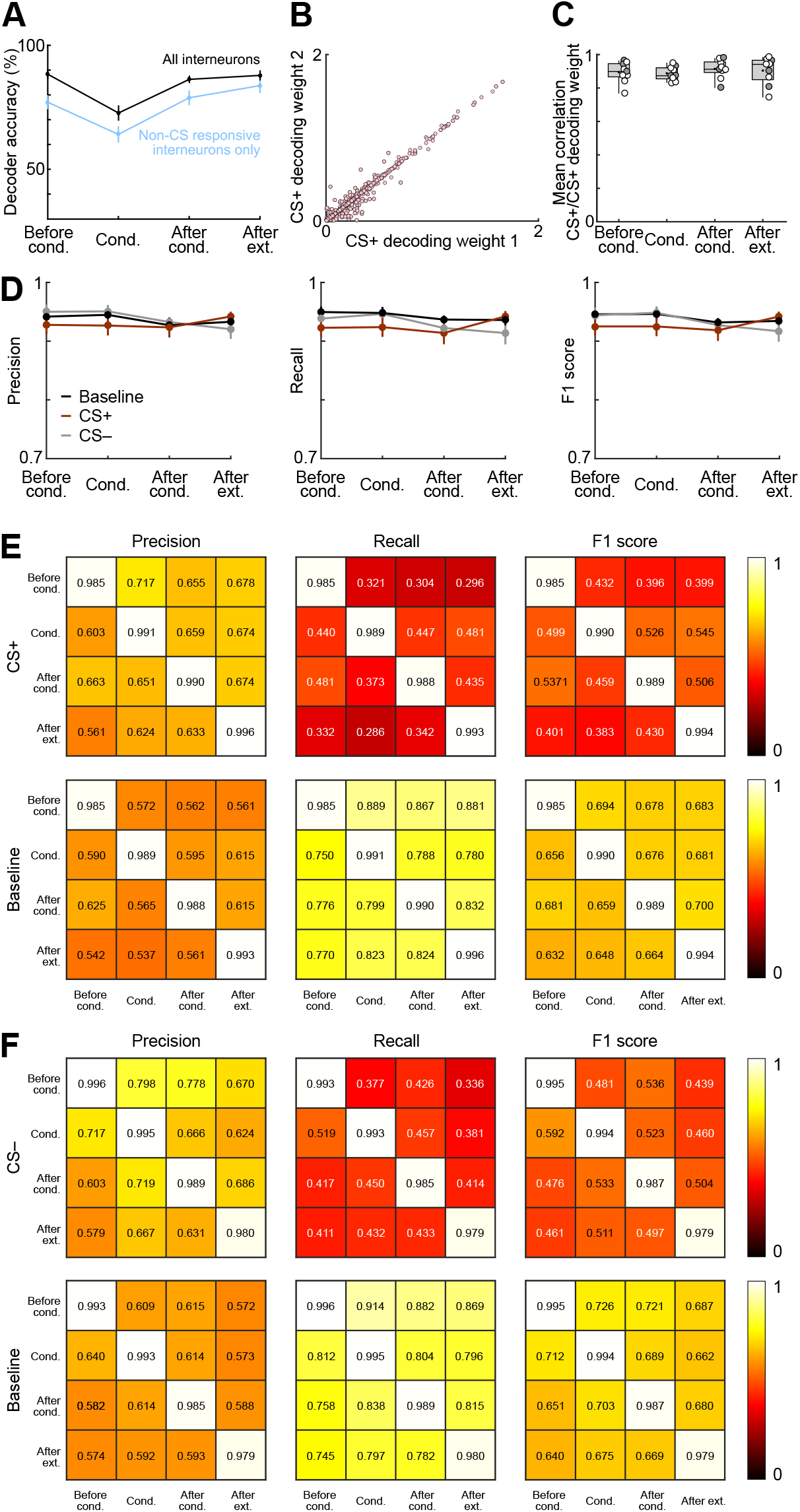
Performance metrics of classifiers. **A**, Comparison of mean accuracy of multiclass intra-day decoders of CS+, CS– and baseline for each day of the behavioural paradigm, averaged across all animals (N = 9 mice) and iterations (n = 100 iterations). Light blue bars indicate the accuracy of decoders trained on a random sample of non-CS-responsive cells. Black bars represent the accuracy of control decoders trained on a random sample of cells selected from all interneurons (regardless of CS responsiveness), with the number of cells matched to those used in the non-CS decoders. **B**, Example scatterplot showing the absolute value of the decoding weight for the CS+ at iteration *i*, and at iteration *i+1*, from all animals and interneurons after conditioning (n = 519 cells, N = 9 mice). **C**, Average correlation between CS+_*i*_ and CS+_*i+1*_ decoding weights for each session, calculated per animal (N = 9 mice). Correlations were calculated between the decoding weights of the *ith* and the *ith+1* iteration, using a total of 200 iterations to obtain 100 correlation values. **D**, Precision, recall and F1 score calculated for each class independently on each day for the intraday multiclass decoders classifying CS+, CS– and baseline. **E**, Precision, recall and F1 score calculated for each class for the intra and inter day performance of the two-way decoder classifying CS+ and baseline. **F**, Precision, recall and F1 score calculated for each class for the intra and inter day performance of the two-way decoder classifying CS– and baseline. Tukey box-and-whisker plots in C show median values, 25^th^ and 75^th^ percentiles, and min to max whiskers with exception of outliers, dots indicate the mean, circles represent individual animals (open circles, imaging sites in the basal amygdala (N = 6); filled circles, at the border of the lateral and basal amygdala (N = 3), see also Supplementary Fig. 1). All other plots display means from all animals (N = 9 mice) and all iterations (n = 100). Favila, Capece Marsico et al. 2024 – Supplementary Figure 6

**Supplementary Figure 6:**
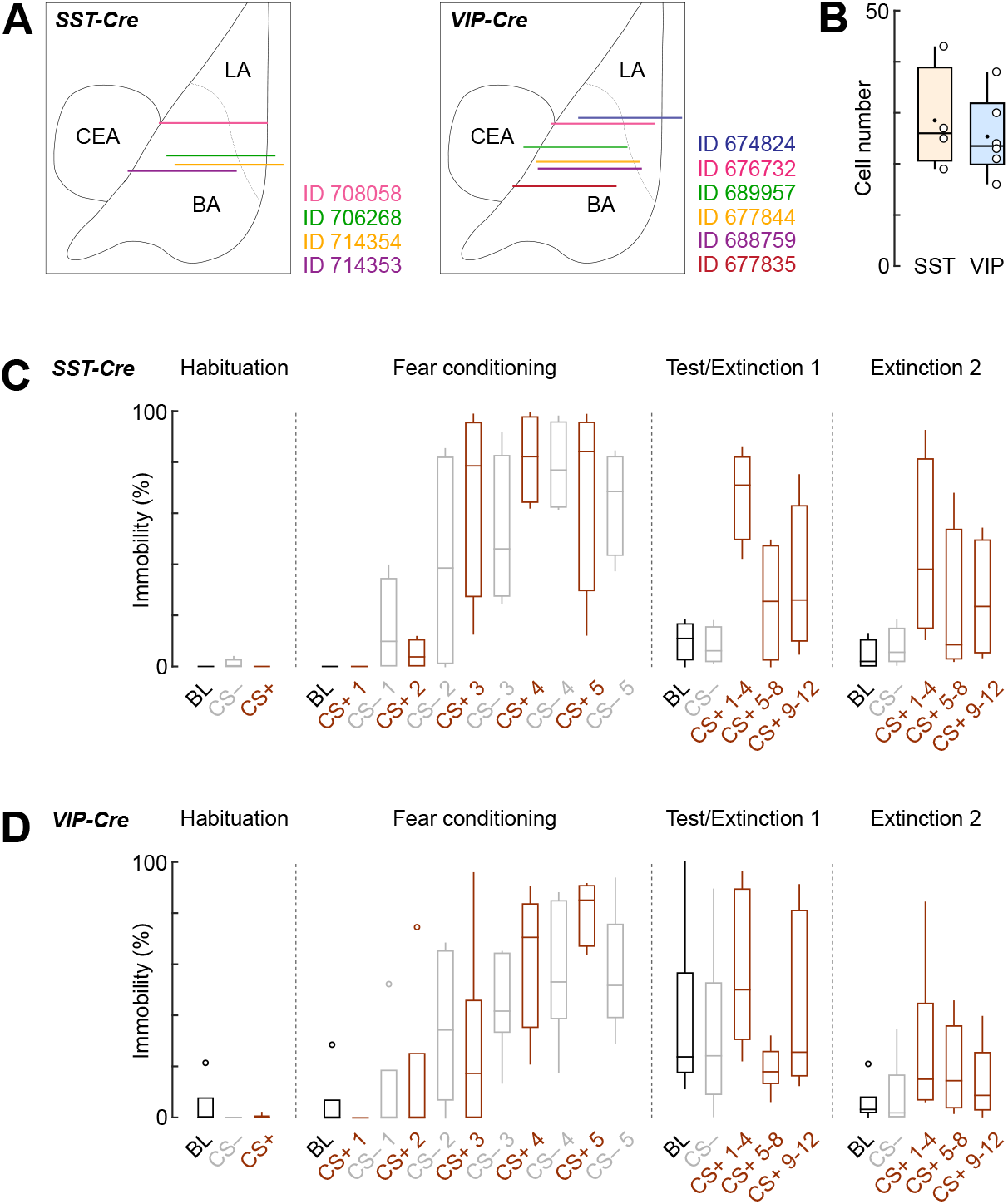
Imaging of molecular interneuron subpopulations during fear and extinction learning. **A**, Schematic illustrating all reconstructed implant sites of GRIN lenses (lens front) within the BLA of *SST-Cre* (N = 4) and *VIP-Cre* mice (N = 6) for deep brain imaging experiments matched to a mouse brain atlas. LA, lateral amygdala; BA, basal amygdala; CEA, central amygdala. **B**, Average cell numbers recorded across the four-day paradigm (SST, N = 4; VIP, N = 6). Box-and-whisker plots show median values, 25^th^ and 75^th^ percentiles, and min to max whiskers, dots indicate the mean, circles are individual animals. **C**, Immobility levels throughout the fear conditioning and extinction paradigm in GRIN lens-implanted *SST-Cre* mice (N = 4). Tukey box-and-whisker plots illustrates median values, 25th and 75th percentiles, and min to max whiskers. **D**, Immobility levels throughout the fear conditioning and extinction paradigm in GRIN lens-implanted *VIP-Cre* mice (N = 6). Tukey box-and-whisker plots illustrate median values, 25^th^ and 75^th^ percentiles, and min to max whiskers, circles indicate outliers. Favila, Capece Marsico et al. 2024 – Supplementary Figure 7

**Supplementary Figure 7:**
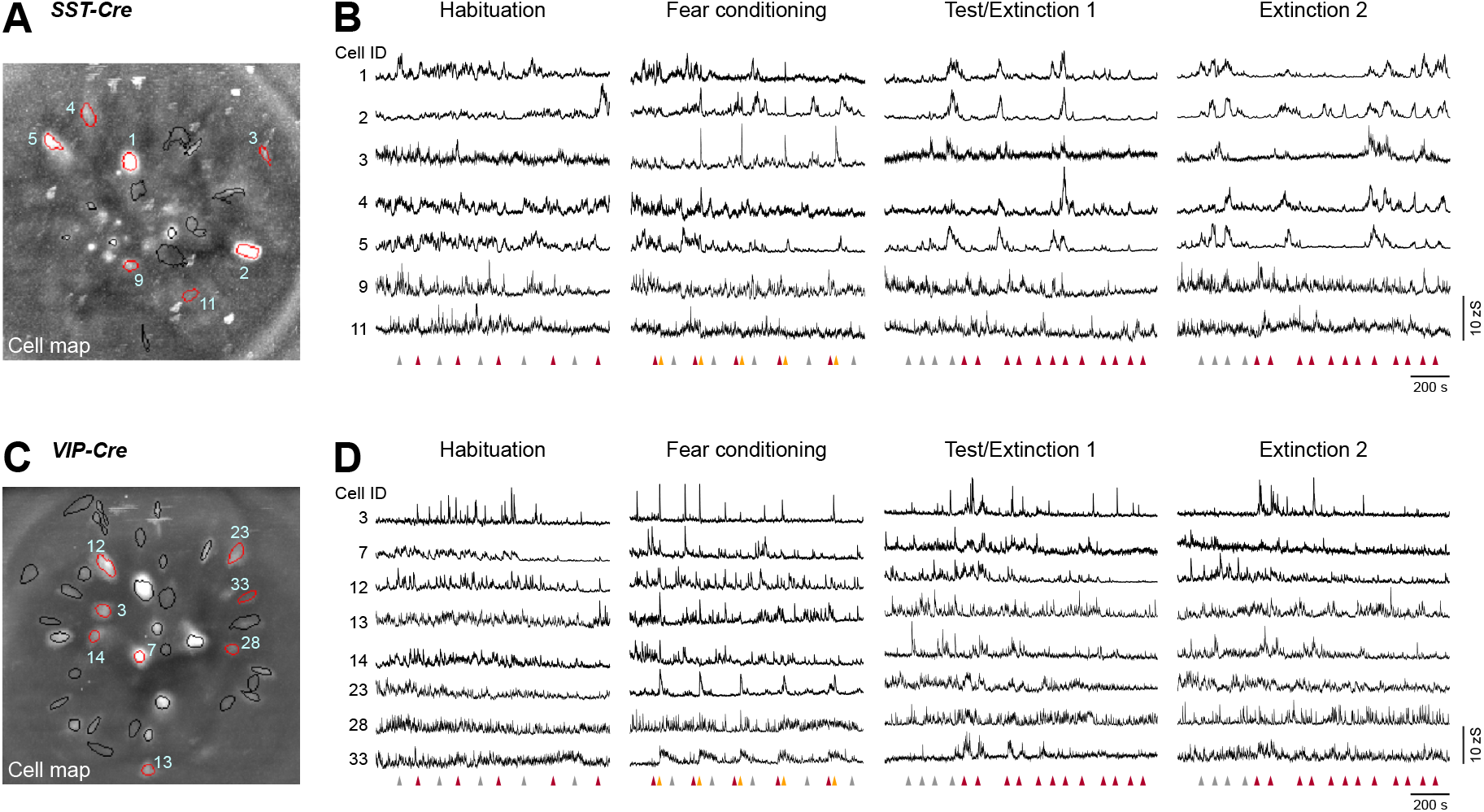
Recording calcium activity across days in *SST-Cre* and *VIP-Cre* mice. **A**, Example field of view (maximum intensity projection across four-day paradigm) for an *SST-Cre* mouse. Circles indicate selected individual components. **B**, Representative example traces from the same animal. Cell IDs correspond to SST interneurons highlighted with red outlines in A. **C**, Example field of view (maximum intensity projection across four-day paradigm) for a *VIP-Cre* mouse. Circles indicate selected individual components. **D**, Representative example traces from the same animal. Cell IDs correspond to VIP interneurons highlighted with red outlines in C. Arrows in B and D indicate starting points of CS+ (red), CS– (grey) and US (yellow).

**Supplementary Figure 8:**
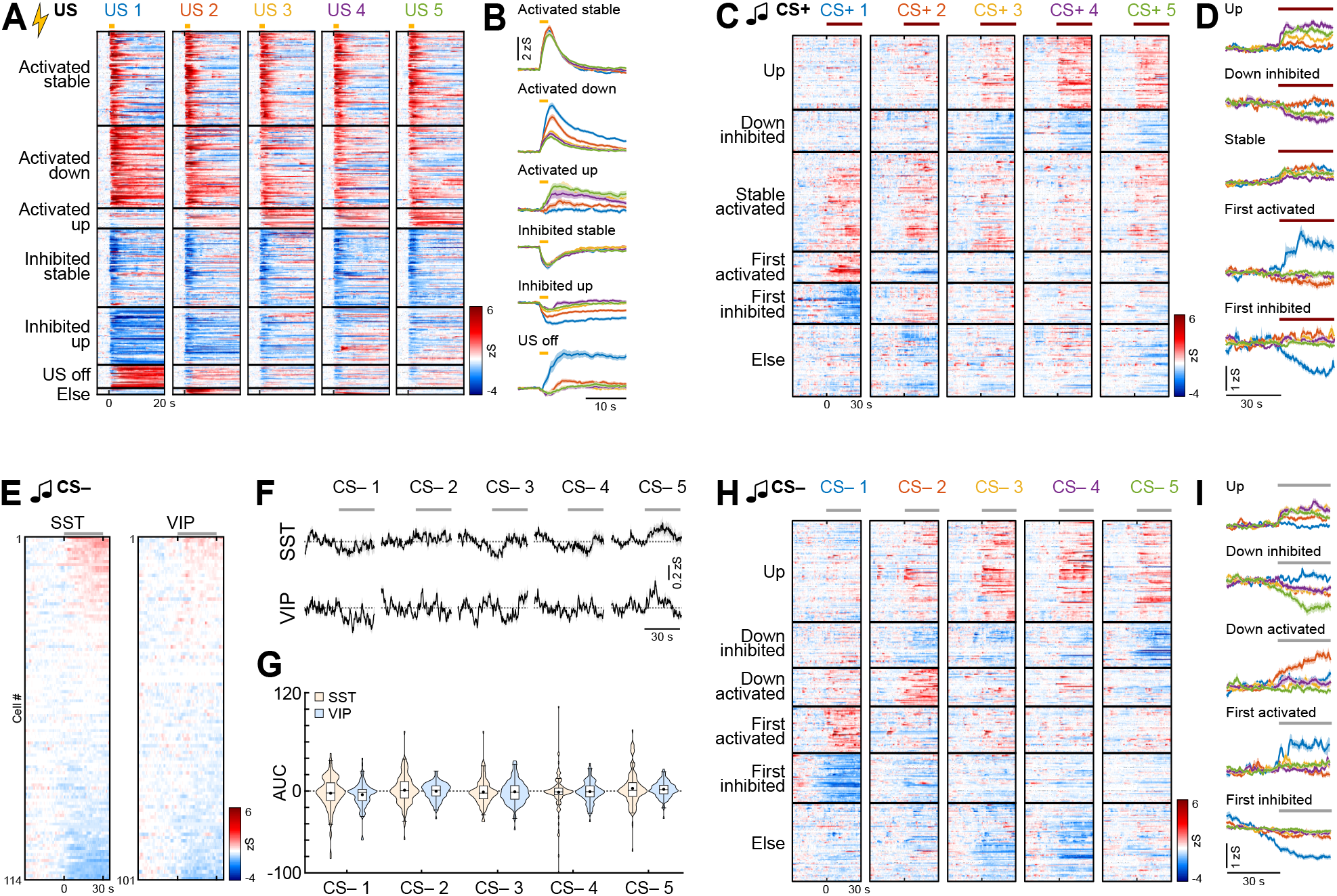
Responses of interneuron subpopulations during fear learning. **A**, Heatmap of US responses in BLA interneurons (including data from all *GAD2-Cre, SST-Cre* and *VIP-Cre* mice) clustered into groups depending on their US response pattern across the five trials (n = 562 responsive cells; ‘Activated stable’, n = 146; ‘Activated down’, n = 128; ‘Activated up’, n = 31; ‘Inhibited stable’, n = 122; ‘Inhibited up’, n = 89; ‘US off’, n = 36; ‘Else’, n = 10). **B**, Average traces of US clusters shown in A. **C**, Heatmap of CS+ responses in BLA interneurons (including data from all *GAD2-Cre, SST-Cre* and *VIP-Cre* mice) clustered into groups depending on their response pattern across the five trials (n = 428 responsive cells; ‘Up’, n = 88; ‘Down inhibited’, n = 50; Stable activated’, n = 118; ‘First activated’, n = 37; ‘First inhibited’, n = 49; ‘Else’, n = 86). **D**, Average traces of CS+ clusters shown in C. **E**, Heatmap of SST and VIP BLA interneuron responses to the control CS– during conditioning (averaged across all five presentations), sorted by individual response amplitude (SST, n = 114 cells from N = 4 mice; VIP, n = 101, N = 4). **F**, Average CS– responses in SST and VIP interneurons across the five presentations (SST, n = 114; VIP, n = 101). **G**, Area under the curve (AUC) during CS– presentations in conditioning for SST and VIP interneurons (SST, n = 114; VIP, n = 101). **H**, Heatmap of CS– responses in BLA interneurons (including data from all *GAD2-Cre, SST-Cre* and *VIP-Cre* mice) clustered into groups depending on their response pattern across the five trials (n = 441 responsive cells; ‘Up’, n = 124; ‘Down inhibited’, n = 56; ‘Down activated’, n = 47; ‘First activated’, n = 57; ‘First inhibited’, n = 61; ‘Else’, n = 96). **I**, Average traces of CS– clusters shown in H. Average traces in B, D, F and I are mean with s.e.m.; violin plots in G show distribution of all data points, Tukey box-and-whisker plots in show median values, 25^th^ and 75^th^ percentiles, and min to max whiskers with exception of outliers, dots indicate the mean. Additional details of statistical analyses are provided in Supplementary Table 1.

**Supplementary Figure 9:**
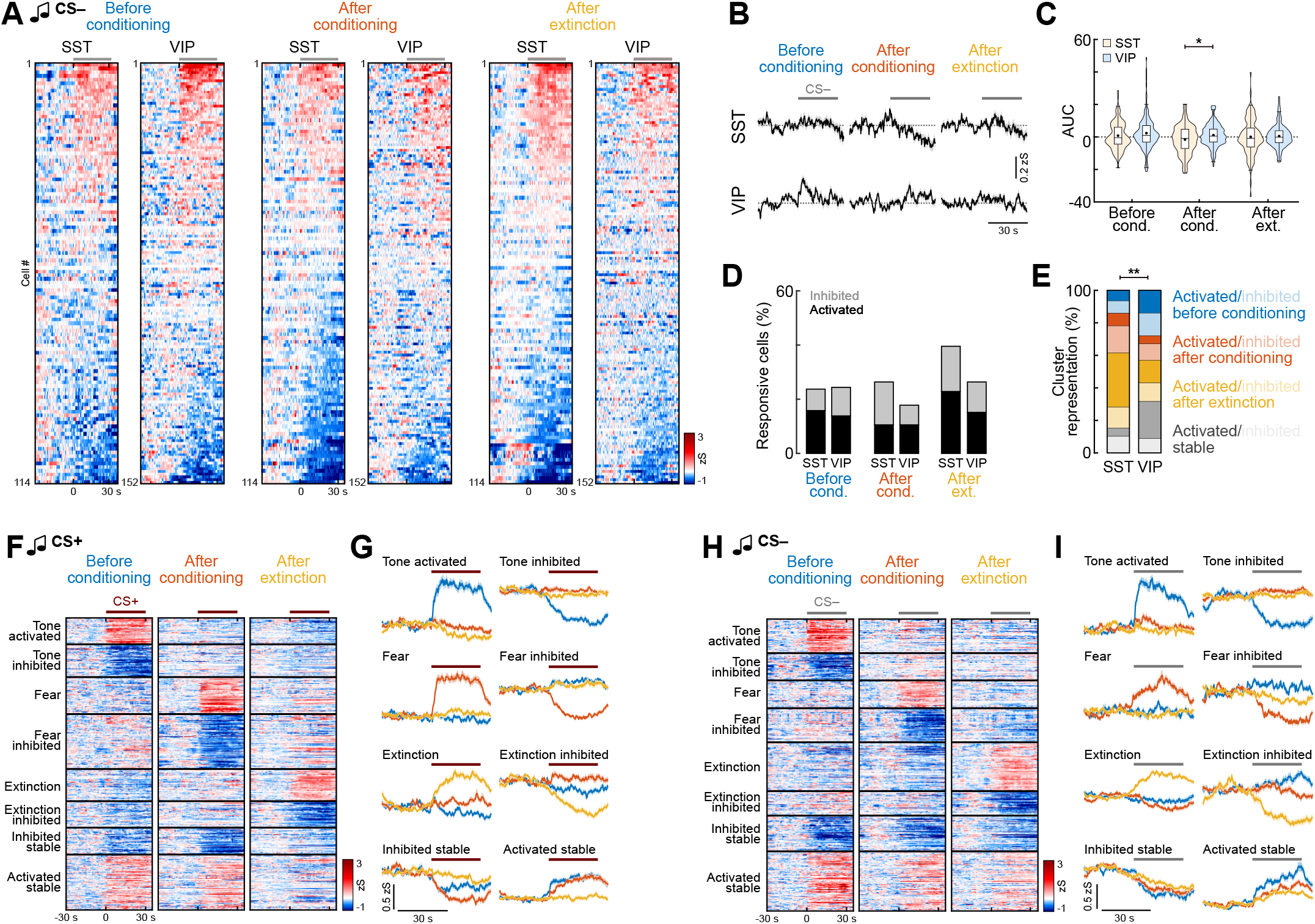
Across-day plasticity of interneuron subpopulations. **A**, Heatmap of SST and VIP interneuron responses to the control CS– before conditioning, after conditioning and after extinction (averaged across four presentations each), sorted individually by response amplitude (SST, n = 114 cells from N = 4 mice; VIP, n = 152, N = 6). Grey line indicates CS duration. **B**, Corresponding average CS– responses in SST and VIP interneurons across days (SST, n = 114; VIP, n = 152). **C**, Area under the curve (AUC) during CS– presentations in conditioning for SST and VIP interneurons SST, n = 114; VIP, n = 152). Mann-Whitney test with Bonferroni correction; SST vs. VIP; After conditioning’, *p* = 0.0324. **D**, Proportions of responsive neurons across the behavioural paradigm (SST, n = 114; VIP, n = 152). **E**, Proportion of cells in CS– clusters for SST and VIP interneurons (SST, n = 78; VIP, n = 79). Chi-Square test (χ2(7) = 20.415), *p* = 0.0047; SST vs. VIP, *post hoc* Chi-Square test with Bonferroni correction, ‘Stable activated’, *p* = 0.0249. **F**, Heatmap of CS+ responses in BLA interneurons across days (including data from all *GAD2-Cre, SST-Cre* and *VIP-Cre* mice) clustered into groups depending on their response pattern (n = 553 responsive cells; ‘Tone activated’/activated before conditioning, n = 50; ‘Tone inhibited’/inhibited before conditioning, n = 62; ‘Fear’/activated after conditioning, n = 70; ‘Fear inhibited’/inhibited after conditioning n = 104; ‘Extinction’/ activated after extinction, n = 61; ‘Extinction inhibited’/inhibited after extinction’, n = 50; ‘Activated stable’, n = 51; ‘Inhibited stable’, n = 105). **G**, Average traces of CS+ clusters shown in F. **H**, Heatmap of CS– responses in BLA interneurons across days (including data from all *GAD2-Cre, SST-Cre* and *VIP-Cre* mice) clustered into groups depending on their response pattern (n = 514 responsive cells; ‘Tone activated’/activated before conditioning, n = 60; ‘Tone inhibited’/inhibited before conditioning, n = 48; ‘Fear’/activated after conditioning, n = 49; ‘Fear inhibited’/inhibited after conditioning, n = 60; ‘Extinction’/activated after extinction, n = 86; ‘Extinction inhibited’/inhibited after extinction, n = 43; ‘Activated stable’, n = 63; ‘Inhibited stable’, n = 105). **I**, Average traces of CS– clusters shown in H. Average traces in B, G and I are mean with s.e.m.; violin plots in C show distribution of all data points, Tukey box-and-whisker plots in show median values, 25^th^ and 75^th^ percentiles, and min to max whiskers with exception of outliers, dots indicate the mean. \**p* < 0.05, \*\**p* < 0.01. Additional details of statistical analyses are provided in Supplementary Table 1. Favila, Capece Marsico et al. 2024 – Supplementary Table 1

**Supplementary Table 1:**
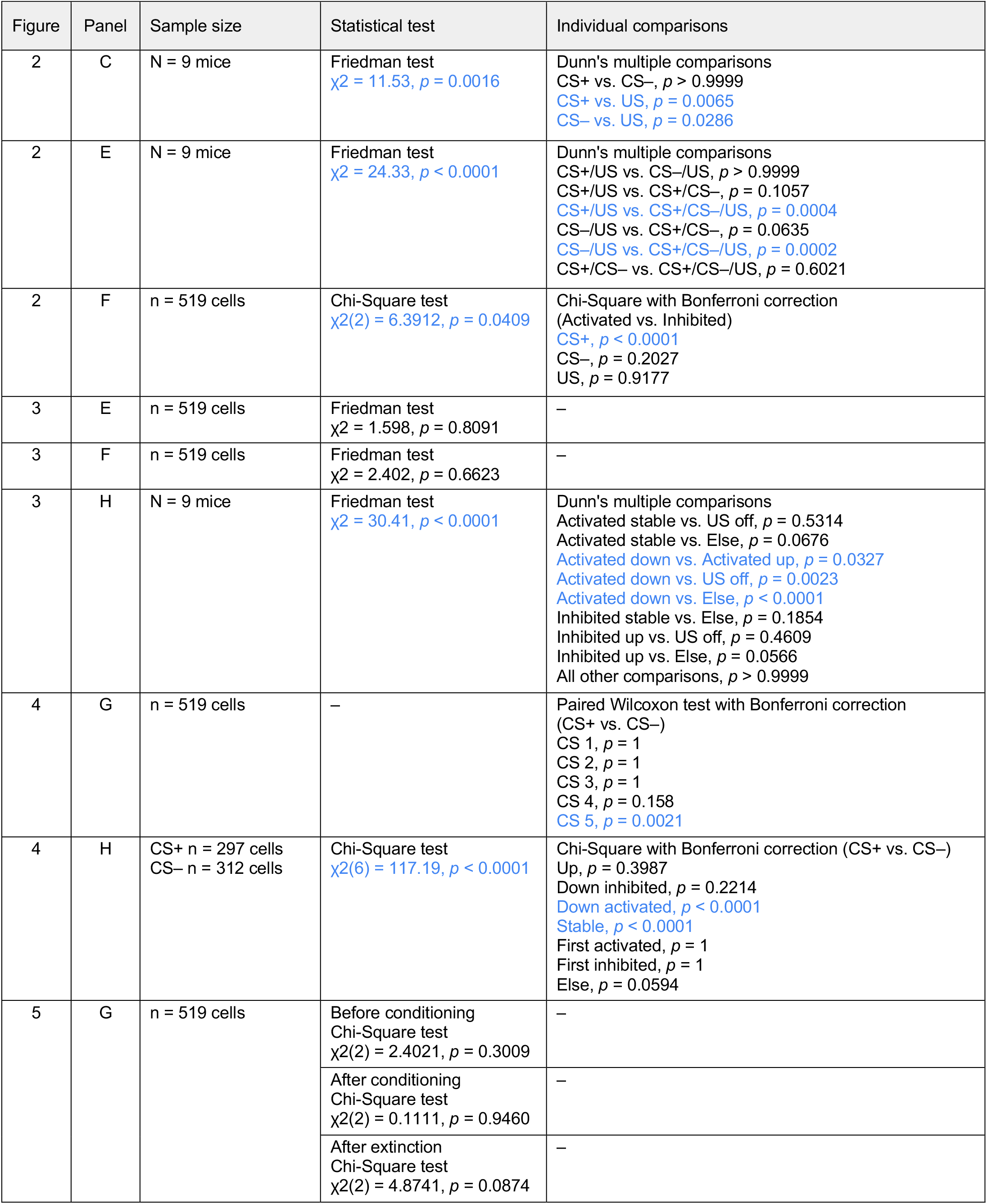
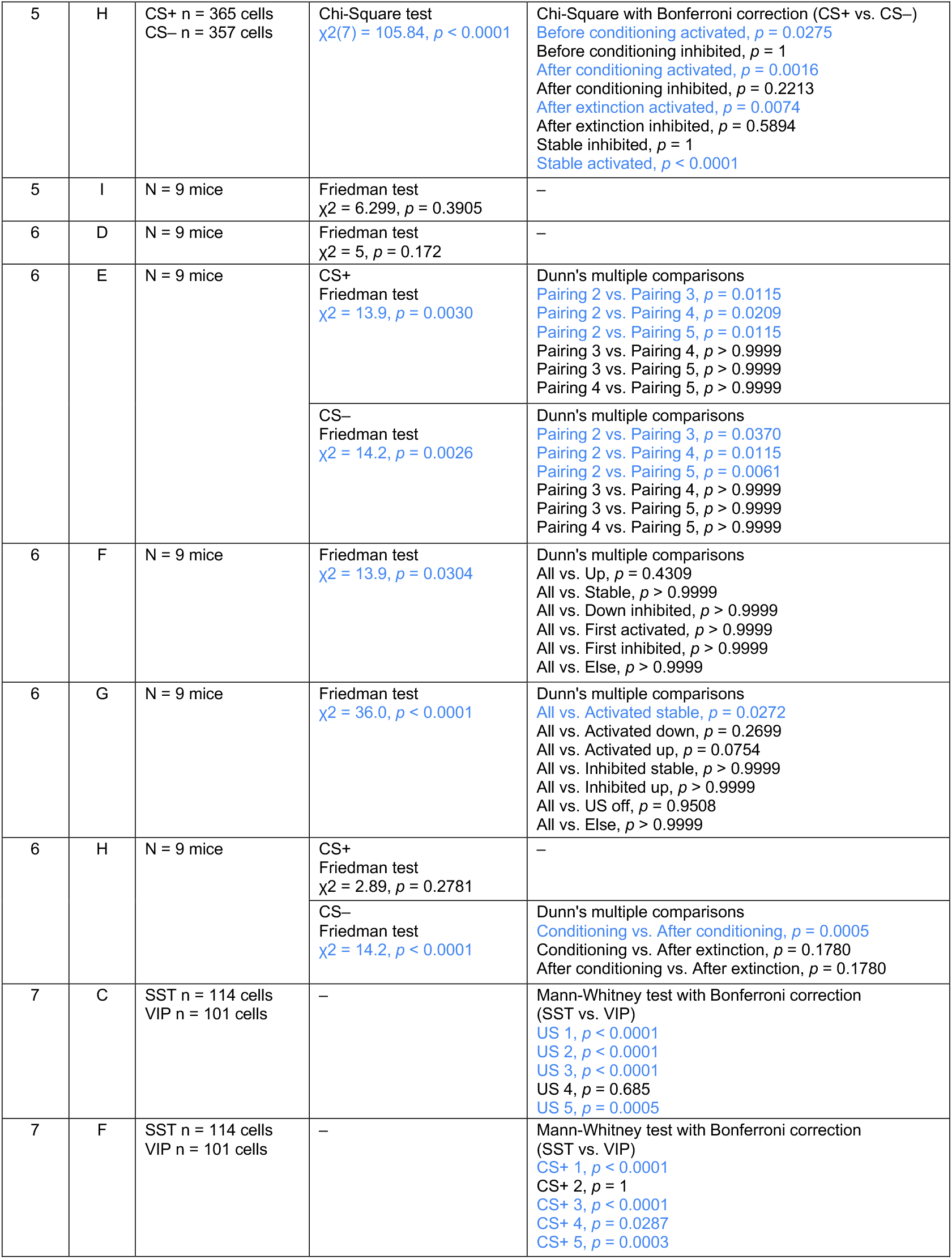
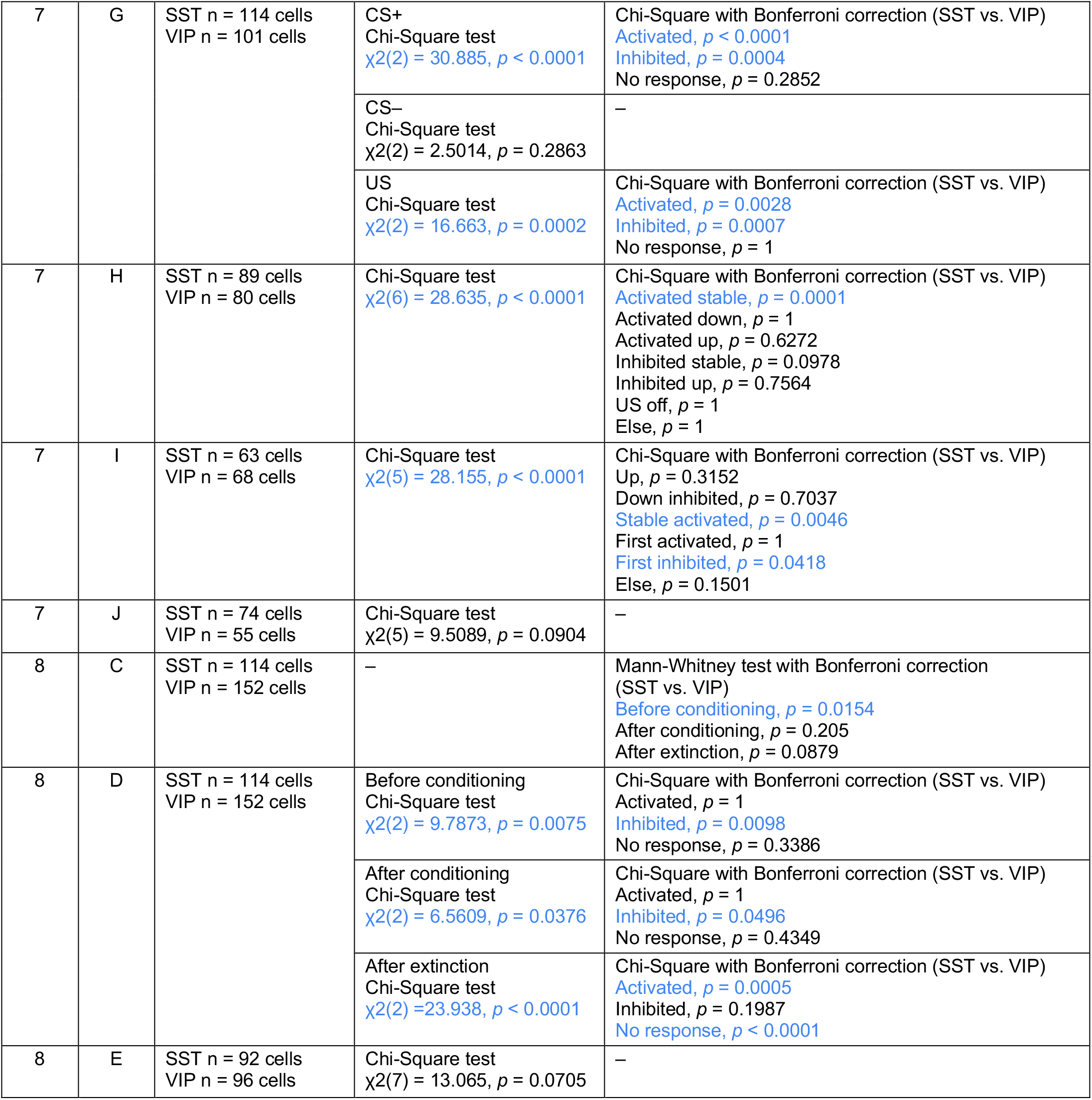
Summary of all statistical analyses for data presented in main and supplementary figures.

## Supplementary Figures

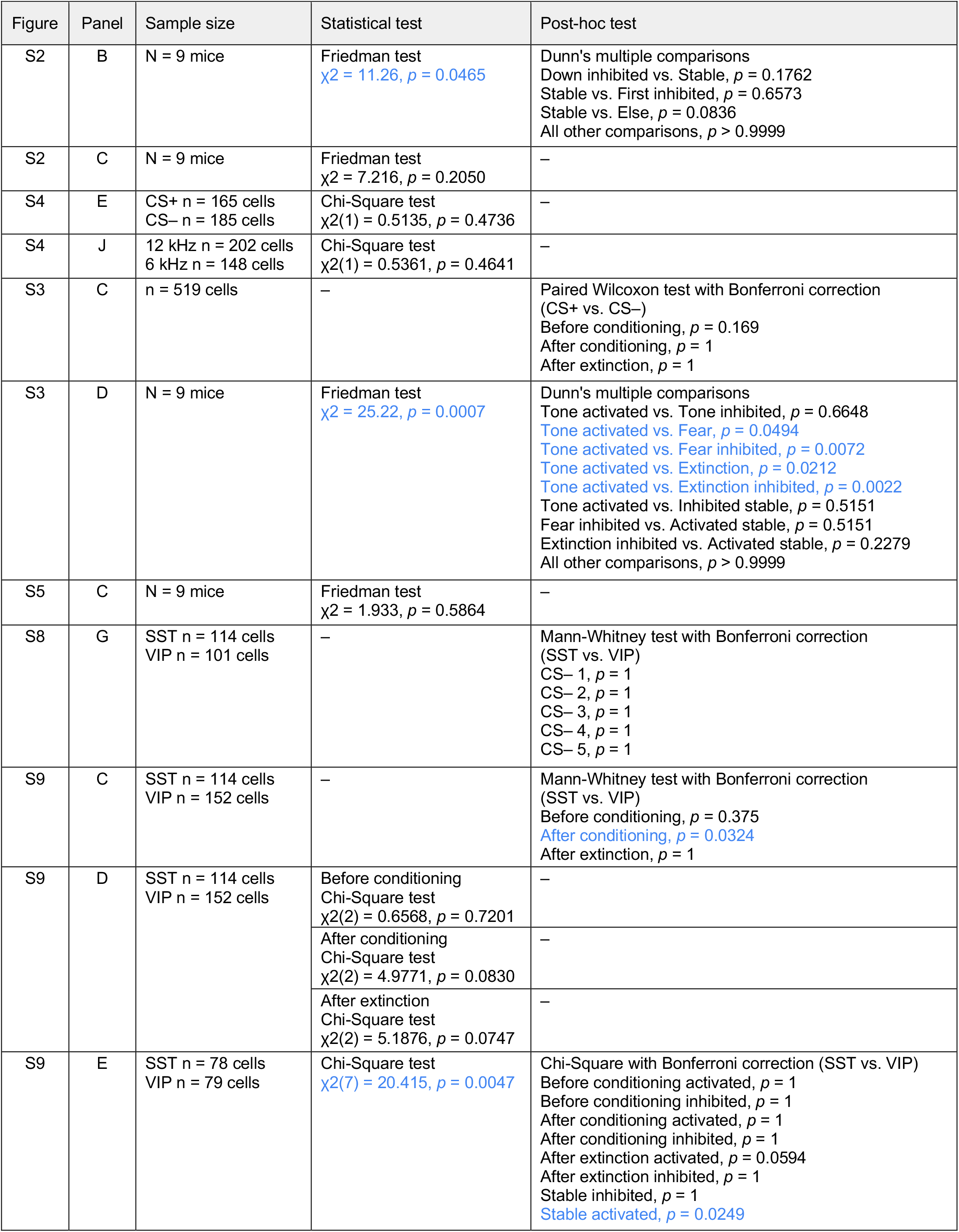

## Notes

### Competing Interest Statement

The authors have declared no competing interest.

