## Supplementary Figures 1-9 for "Heterogeneous plasticity of amygdala interneurons in associative learning and extinction"

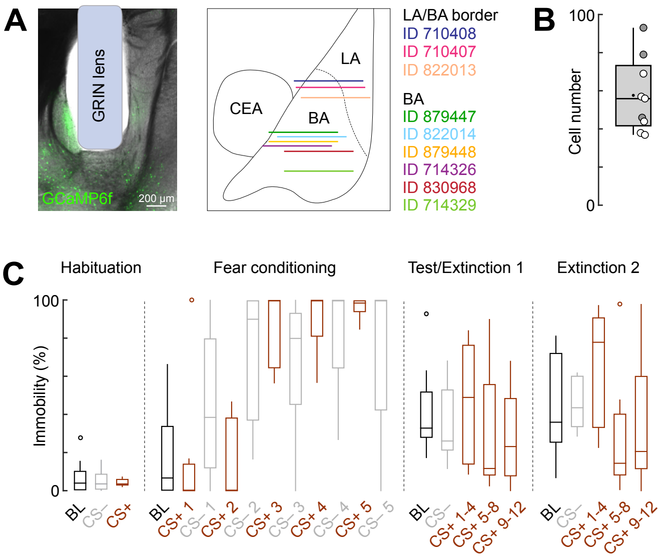


**Supplementary Figure 1: Imaging of basolateral amygdala interneurons during fear and extinction learning**

**A**, Representative implant site (ID 879448) and schematic illustrating all reconstructed implant sites of GRIN lenses (lens front) within the BLA of *GAD2-Cre* mice for deep brain imaging experiments matched to a mouse brain atlas (N = 9 mice). LA, lateral amygdala; BA, basal amygdala; CEA, central amygdala. **B**, Average cell numbers recorded across the four-day paradigm (N = 9). Tukey box-and-whisker plot illustrates median values, 25$^{th}$ and 75$^{th}$ percentiles, and min to max whiskers, dot indicates the mean. Circles represent individual animals (open circles, imaging sites in the basal amygdala (N = 6); filled circles, at the border of the lateral and basal amygdala (N = 3). **C**, Immobility levels throughout the fear conditioning and extinction paradigm in GRIN lens-implanted *GAD2-Cre* mice (N = 9). Tukey box-and-whisker plots illustrates median values, 25$^{th}$ and 75$^{th}$ percentiles, and min to max whiskers, circles indicate outliers.

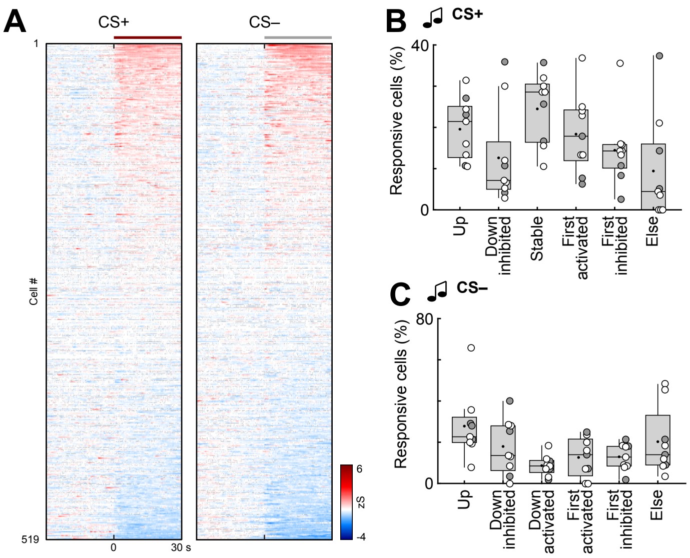


**Supplementary Figure 2: CS coding in amygdala interneurons during fear conditioning**

**A**, Heatmap of CS+ and CS– responses in BLA interneurons during conditioning, averaged across all five trials and ordered individually by response amplitude (n = 519 cells from N = 9 mice). Lines indicate CS duration. **B**, Fraction of interneurons according to CS+ cluster membership across animals (see Figure 4; N = 9). Friedman test ($\chi 2 = 11.26$), $p = 0.0465$; followed by Dunn's multiple comparisons (non-significant). **C**, Fraction of interneurons according to CS– cluster membership across animals (see Figure 4; N = 9).

Tukey box-and-whisker plots in B and C show median values, 25$^{th}$ and 75$^{th}$ percentiles, and min to max whiskers with exception of outliers, dots indicate the mean. Circles represent individual animals (open circles, imaging sites in the basal amygdala (N = 6); filled circles, at the border of the lateral and basal amygdala (N = 3). Additional details of statistical analyses are provided in Supplementary Table 1.

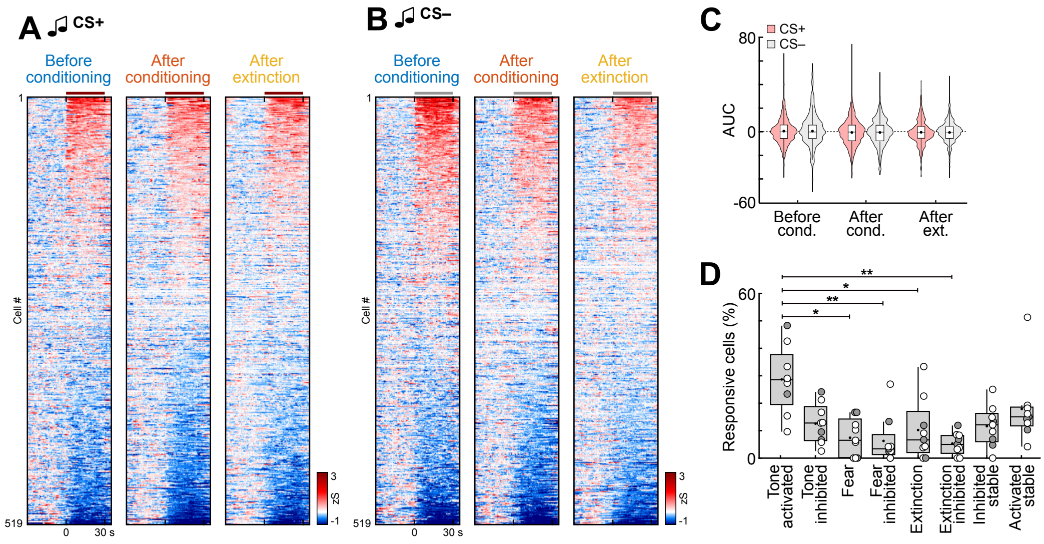


**Supplementary Figure 3: CS encoding across learning days in basolateral amygdala interneurons**

**A**, Heatmap of CS+ and **B**, CS– responses in BLA interneurons before conditioning, after conditioning and after extinction, averaged across the four CS presentations used for clustering (n = 519 cells from N = 9 mice). Line indicates CS duration. **C**, Comparison of area under the curve (AUC) for CS+ and CS– responses in BLA interneurons (n = 519). **D**, Fraction of interneurons according to CS– cluster membership across animals (see Figure 5; N = 9). Friedman test ($\chi2 = 25.22$), $p = 0.0007$, followed by Dunn's multiple comparisons (‘Tone activated’/activated before conditioning vs. ‘Fear’/activated after conditioning, $p = 0.0494$; ‘Tone activated’/activated before conditioning vs. ‘Fear inhibited’/inhibited after conditioning, $p = 0.0072$; ‘Tone activated’/activated before conditioning vs. ‘Extinction’/activated after extinction, $p = 0.0212$; ‘Tone activated’/activated before conditioning vs. ‘Extinction inhibited’/inhibited after extinction, $p = 0.0022$).

Violin plots in C show distribution of all data points, Tukey box-and-whisker plots in C and D show median values, $25^{th}$ and $75^{th}$ percentiles, and min to max whiskers with exception of outliers, dots indicate the mean. Circles in D represent individual animals (open circles, imaging sites in the basal amygdala (N = 6); filled circles, at the border of the lateral and basal amygdala (N = 3). *$p < 0.05$, **$p < 0.01$. Additional details of statistical analyses are provided in Supplementary Table 1.

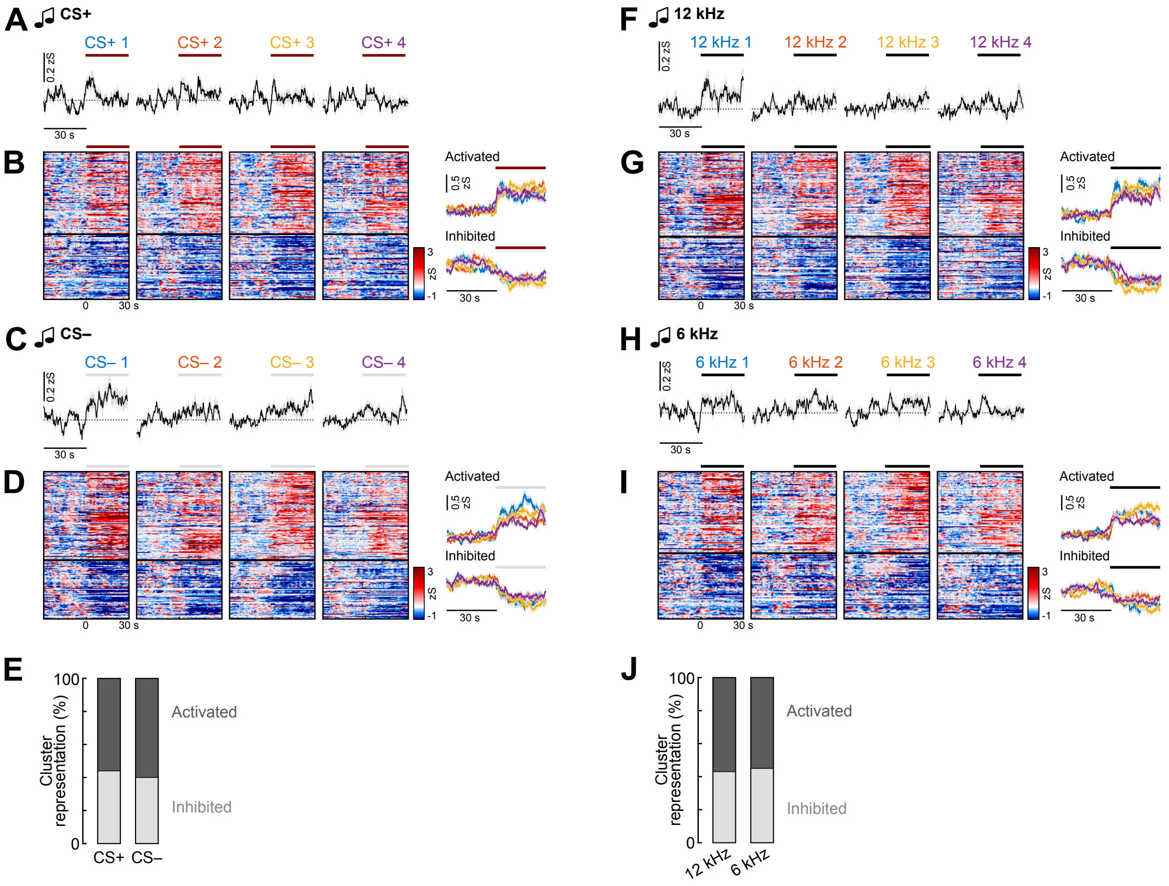


**Supplementary Figure 4: Interneuron responses to auditory stimuli**

**A**, Average traces of basolateral amygdala interneurons during the first four CS+ presentations during habituation (n = 519 cells from N = 9 mice; counterbalanced for 6 kHz and 12 kHz). Line indicates CS duration. **B**, Heatmap (left) of CS+ responses clustered into groups depending on their response pattern across the four presentations and corresponding average traces of clusters (right). n = 165 responsive cells; 'Activated', n = 92; 'Inhibited', n = 73. **C**, Average traces of basolateral amygdala interneurons during the first four CS– presentations during habituation (n = 519; counterbalanced for 6 kHz and 12 kHz). **D**, Heatmap (left) of CS– responses clustered into groups depending on their response pattern across the four presentations and corresponding average traces of clusters (right). n = 185 responsive cells; 'Activated', n = 111; 'Inhibited', n = 74. **E**, Proportion of cells in CS+ and CS– clusters (CS+, n = 165; CS–, n = 185). **F**, Average traces of basolateral amygdala interneurons during the first four 12 kHz presentations during habituation (n = 519; later assigned to be CS+ or CS–). **G**, Heatmap (left) of 12 kHz responses clustered into groups depending on their response pattern across the four presentations and corresponding average traces of clusters (right). n = 202 responsive cells; 'Activated', n = 116; 'Inhibited', n = 86. **H**, Average traces of basolateral amygdala interneurons during the first four 6 kHz presentations during habituation (n = 519; later assigned to be CS+ or CS–). **I**, Heatmap (left) of 6 kHz responses clustered into groups depending on their response pattern across the four presentations and corresponding average traces of clusters (right). n = 148 responsive cells; 'Activated', n = 82; 'Inhibited', n = 66. **J**, Proportion of cells in 12 kHz and 6 kHz clusters (12 kHz, n = 202; 6 kHz, n = 148).

Average traces across panels are mean with s.e.m.. Additional details of statistical analyses are provided in Supplementary Table 1.

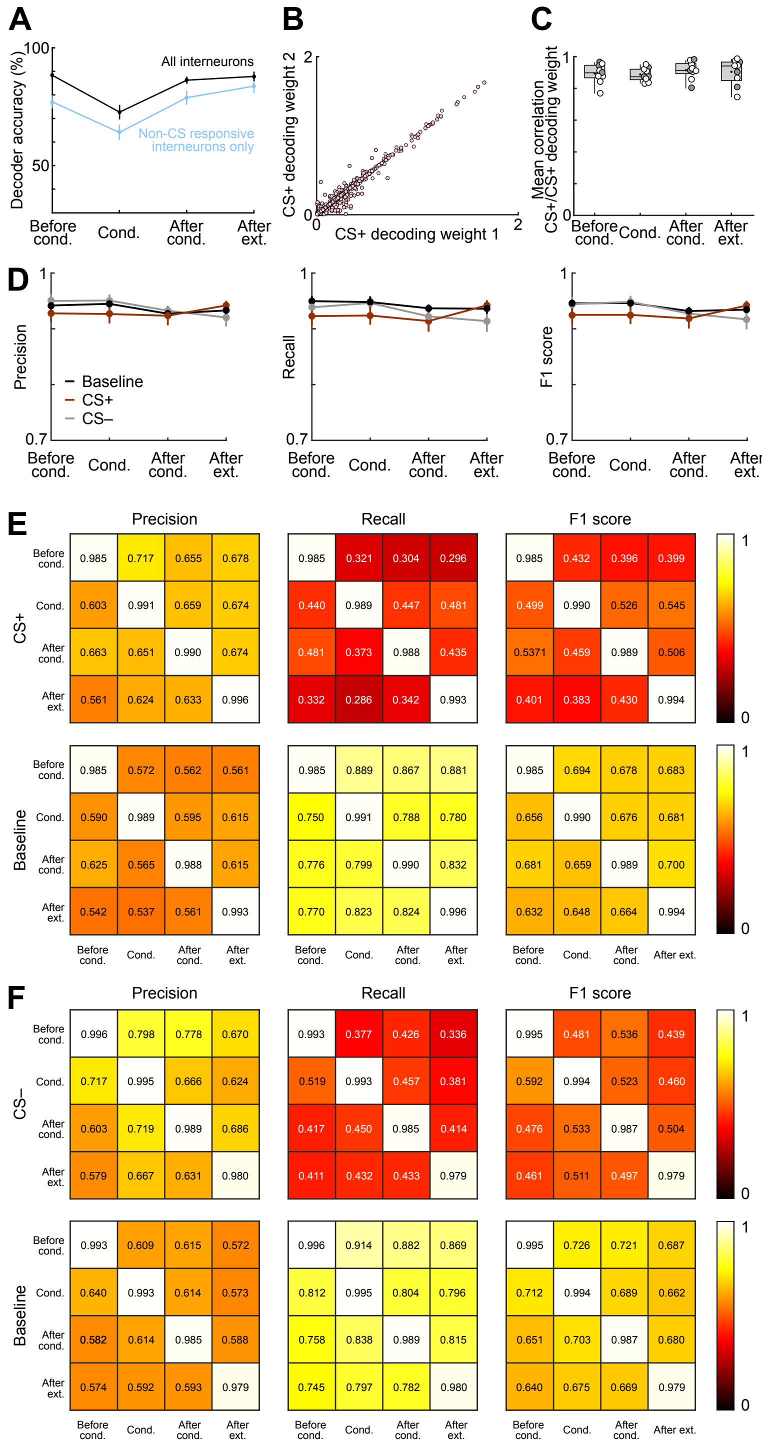


**Supplementary Figure 5: Performance metrics of classifiers**

**A**, Comparison of mean accuracy of multiclass intra-day decoders of CS+, CS– and baseline for each day of the behavioural paradigm, averaged across all animals (N = 9 mice) and iterations (n = 100 iterations). Light blue bars indicate the accuracy of decoders trained on a random sample of non-CS-responsive cells. Black bars represent the accuracy of control decoders trained on a random sample of cells selected from all interneurons (regardless of CS responsiveness), with the number of cells matched to those used in the non-CS decoders. **B**, Example scatterplot showing the absolute value of the decoding weight for the CS+ at iteration *i*, and at iteration *i+1*, from all animals and interneurons after conditioning (n = 519 cells, N = 9 mice). **C**, Average correlation between $CS+_{i}$ and $CS+_{i+1}$ decoding weights for each session, calculated per animal (N = 9 mice). Correlations were calculated between the decoding weights of the *ith* and the *ith+1* iteration, using a total of 200 iterations to obtain 100 correlation values. **D,** Precision, recall and F1 score calculated for each class independently on each day for the intraday multiclass decoders classifying CS+, CS– and baseline. **E,** Precision, recall and F1 score calculated for each class for the intra and inter day performance of the two-way decoder classifying CS+ and baseline. **F,** Precision, recall and F1 score calculated for each class for the intra and inter day performance of the two-way decoder classifying CS– and baseline.

Tukey box-and-whisker plots in C show median values, $25^{th}$ and $75^{th}$ percentiles, and min to max whiskers with exception of outliers, dots indicate the mean, circles represent individual animals (open circles, imaging sites in the basal amygdala (N = 6); filled circles, at the border of the lateral and basal amygdala (N = 3), see also Supplementary Fig. 1). All other plots display means from all animals (N = 9 mice) and all iterations (n = 100).

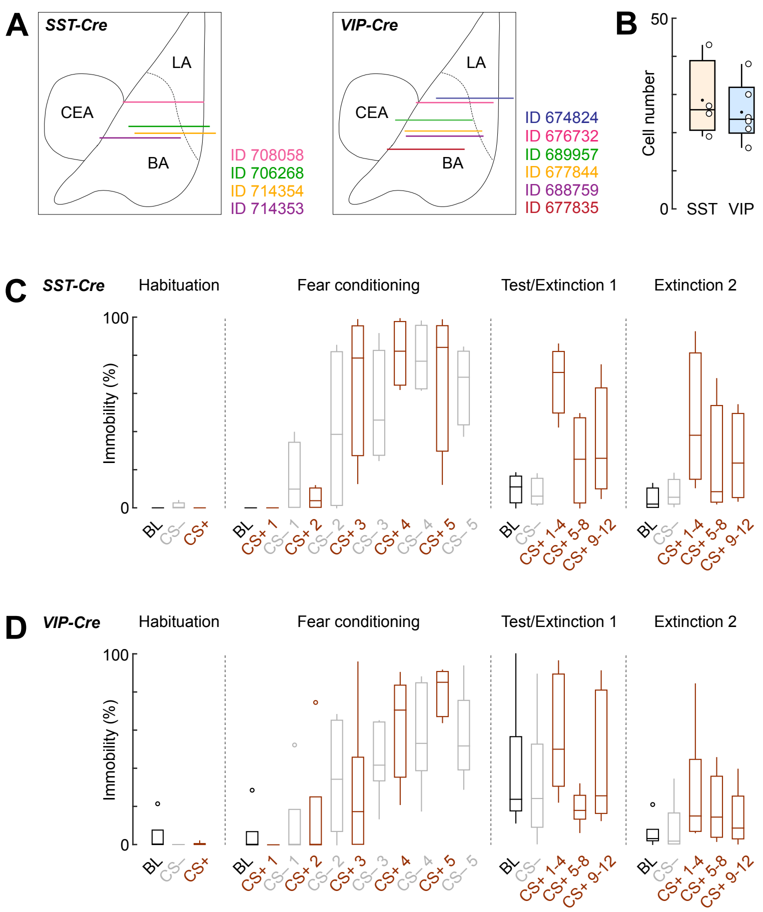


**Supplementary Figure 6: Imaging of molecular interneuron subpopulations during fear and extinction learning**

**A**, Schematic illustrating all reconstructed implant sites of GRIN lenses (lens front) within the BLA of *SST-Cre* (N = 4) and *VIP-Cre* mice (N = 6) for deep brain imaging experiments matched to a mouse brain atlas. LA, lateral amygdala; BA, basal amygdala; CEA, central amygdala. **B**, Average cell numbers recorded across the four-day paradigm (SST, N = 4; VIP, N = 6). Box-and-whisker plots show median values, 25$^{th}$ and 75$^{th}$ percentiles, and min to max whiskers, dots indicate the mean, circles are individual animals. **C**, Immobility levels throughout the fear conditioning and extinction paradigm in GRIN lens-implanted *SST-Cre* mice (N = 4). Tukey box-and-whisker plots illustrates median values, 25th and 75th percentiles, and min to max whiskers. **D**, Immobility levels throughout the fear conditioning and extinction paradigm in GRIN lens-implanted *VIP-Cre* mice (N = 6). Tukey box-and-whisker plots illustrate median values, 25$^{th}$ and 75$^{th}$ percentiles, and min to max whiskers, circles indicate outliers.

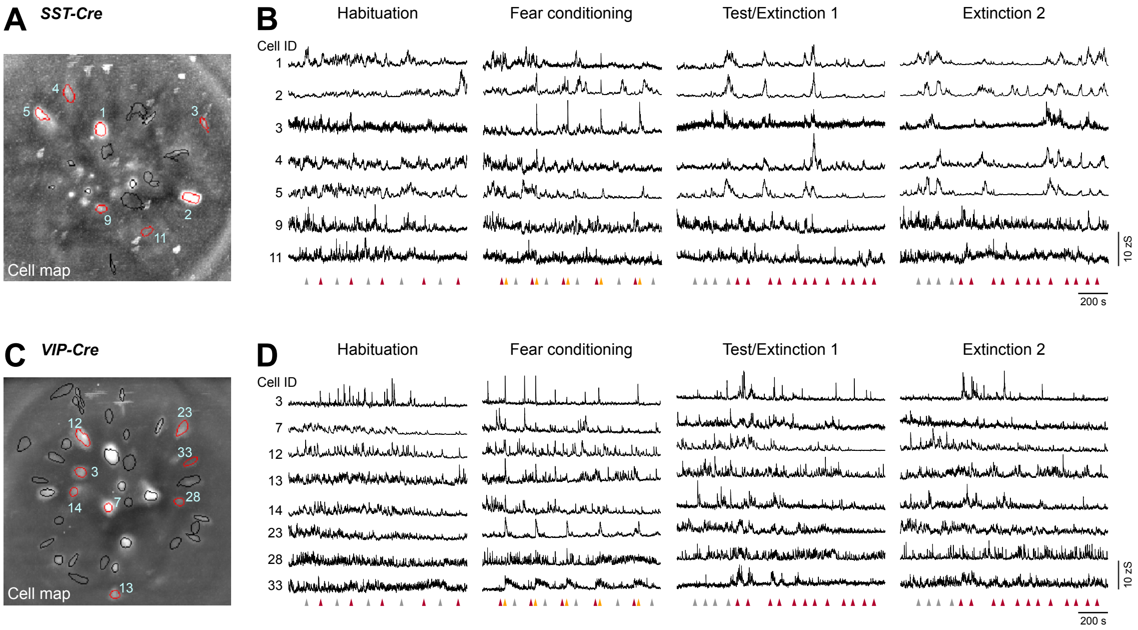


**Supplementary Figure 7: Recording calcium activity across days in *SST-Cre* and *VIP-Cre* mice**

**A**, Example field of view (maximum intensity projection across four-day paradigm) for an *SST-Cre* mouse. Circles indicate selected individual components. **B**, Representative example traces from the same animal. Cell IDs correspond to SST interneurons highlighted with red outlines in A. **C**, Example field of view (maximum intensity projection across four-day paradigm) for a *VIP-Cre* mouse. Circles indicate selected individual components. **D**, Representative example traces from the same animal. Cell IDs correspond to VIP interneurons highlighted with red outlines in C.

Arrows in B and D indicate starting points of CS+ (red), CS– (grey) and US (yellow).

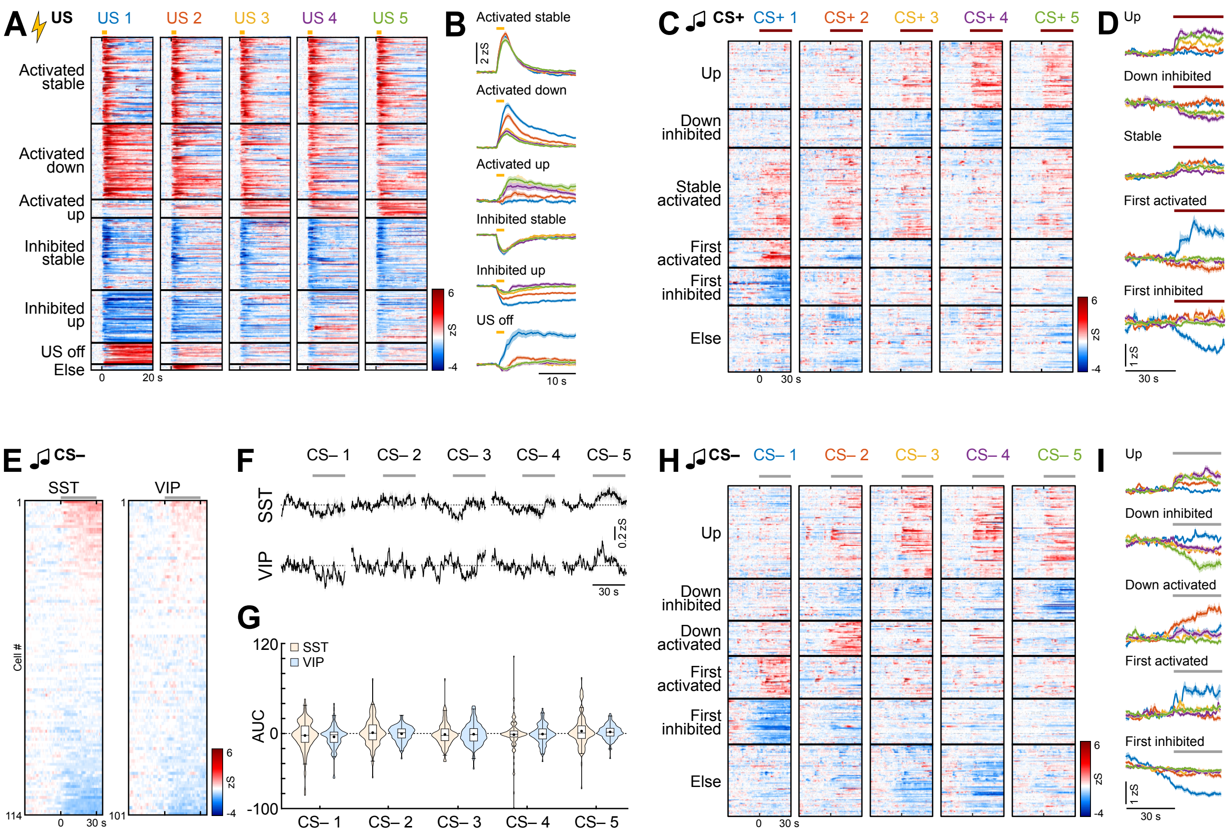

**Supplementary Figure 8: Responses of interneuron subpopulations during fear learning**

**A**, Heatmap of US responses in BLA interneurons (including data from all *GAD2-Cre*, *SST-Cre* and *VIP-Cre* mice) clustered into groups depending on their US response pattern across the five trials (n = 562 responsive cells; 'Activated stable', n = 146; 'Activated down', n = 128; 'Activated up', n = 31; 'Inhibited stable', n = 122; 'Inhibited up', n = 89; 'US off', n = 36; 'Else', n = 10). **B**, Average traces of US clusters shown in A. **C**, Heatmap of CS+ responses in BLA interneurons (including data from all *GAD2-Cre*, *SST-Cre* and *VIP-Cre* mice) clustered into groups depending on their response pattern across the five trials (n = 428 responsive cells; 'Up', n = 88; 'Down inhibited', n = 50; Stable activated', n = 118; 'First activated', n = 37; 'First inhibited', n = 49; 'Else', n = 86). **D**, Average traces of CS+ clusters shown in C. **E**, Heatmap of SST and VIP BLA interneuron responses to the control CS– during conditioning (averaged across all five presentations), sorted by individual response amplitude (SST, n = 114 cells from N = 4 mice; VIP, n = 101, N = 4). **F**, Average CS– responses in SST and VIP interneurons across the five presentations (SST, n = 114; VIP, n = 101). **G**, Area under the curve (AUC) during CS– presentations in conditioning for SST and VIP interneurons (SST, n = 114; VIP, n = 101). **H**, Heatmap of CS– responses in BLA interneurons (including data from all *GAD2-Cre*, *SST-Cre* and *VIP-Cre* mice) clustered into groups depending on their response pattern across the five trials (n = 441 responsive cells; 'Up', n = 124; 'Down inhibited', n = 56; 'Down activated', n = 47; 'First activated', n = 57; 'First inhibited', n = 61; 'Else', n = 96). **I**, Average traces of CS– clusters shown in H.

Average traces in B, D, F and I are mean with s.e.m.; violin plots in G show distribution of all data points, Tukey box-and-whisker plots in show median values, 25th and 75th percentiles, and min to max whiskers with exception of outliers, dots indicate the mean. Additional details of statistical analyses are provided in Supplementary Table 1.

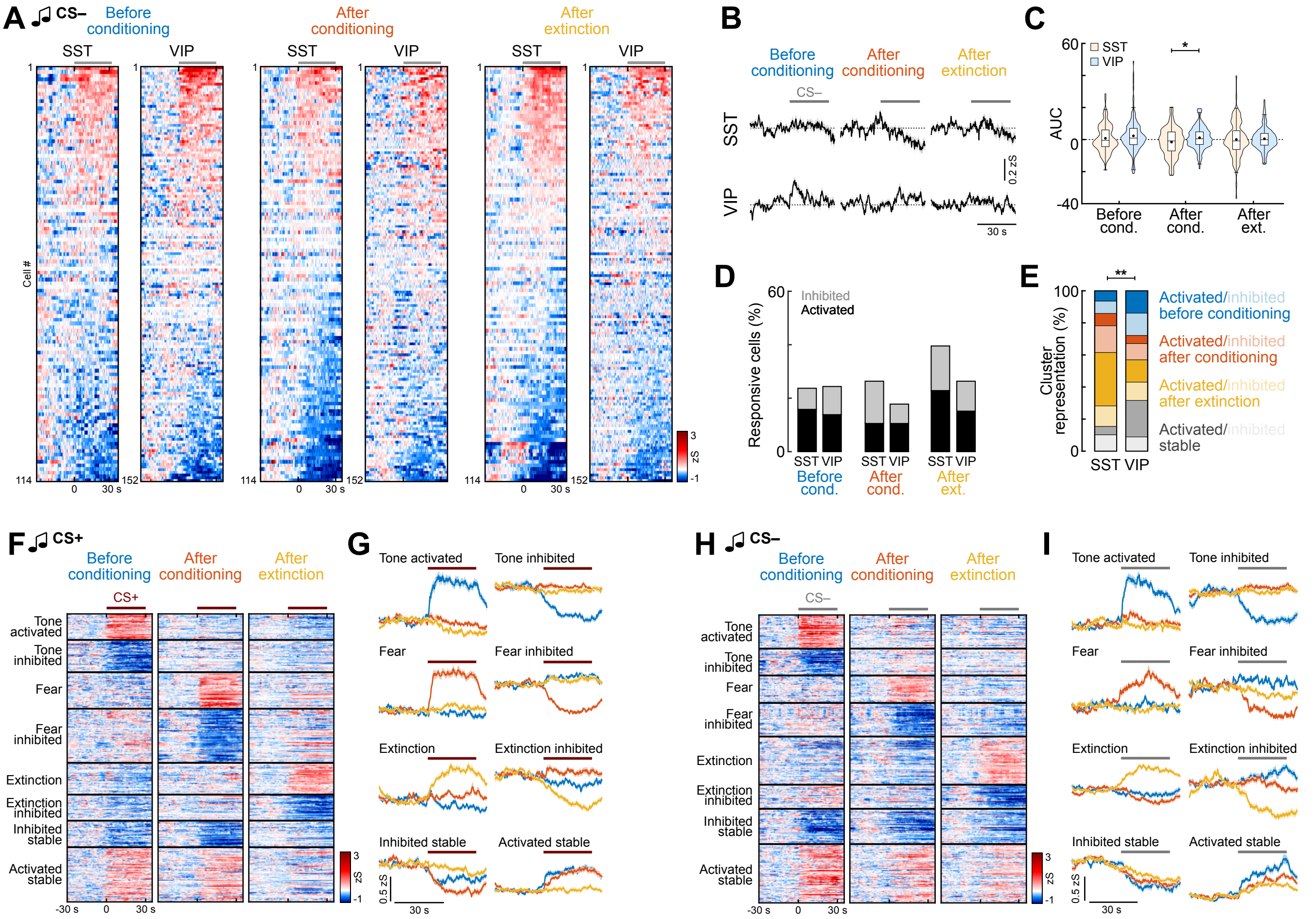


**Supplementary Figure 9: Across-day plasticity of interneuron subpopulations**

**A**, Heatmap of SST and VIP interneuron responses to the control CS– before conditioning, after conditioning and after extinction (averaged across four presentations each), sorted individually by response amplitude (SST, n = 114 cells from N = 4 mice; VIP, n = 152, N = 6). Grey line indicates CS duration. **B**, Corresponding average CS– responses in SST and VIP interneurons across days (SST, n = 114; VIP, n = 152). **C**, Area under the curve (AUC) during CS– presentations in conditioning for SST and VIP interneurons SST, n = 114; VIP, n = 152). Mann-Whitney test with Bonferroni correction; SST vs. VIP; After conditioning', *p* = 0.0324. **D**, Proportions of responsive neurons across the behavioural paradigm (SST, n = 114; VIP, n = 152). **E**, Proportion of cells in CS– clusters for SST and VIP interneurons (SST, n = 78; VIP, n = 79). Chi-Square test (χ2(7) = 20.415), *p* = 0.0047; SST vs. VIP, *post hoc* Chi-Square test with Bonferroni correction, 'Stable activated', *p* = 0.0249. **F**, Heatmap of CS+ responses in BLA interneurons across days (including data from all *GAD2-Cre*, *SST-Cre* and *VIP-Cre* mice) clustered into groups depending on their response pattern (n = 553 responsive cells; 'Tone activated'/activated before conditioning, n = 50; 'Tone inhibited'/inhibited before conditioning, n = 62; 'Fear'/activated after conditioning, n = 70; 'Fear inhibited'/inhibited after conditioning n = 104; 'Extinction'/ activated after extinction, n = 61; 'Extinction inhibited'/inhibited after extinction', n = 50; 'Activated stable', n = 51; 'Inhibited stable', n = 105). **G**, Average traces of CS+ clusters shown in F. **H**, Heatmap of CS– responses in BLA interneurons across days (including data from all *GAD2-Cre*, *SST-Cre* and *VIP-Cre* mice) clustered into groups depending on their response pattern (n = 514 responsive cells; 'Tone activated'/activated before conditioning, n = 60; 'Tone inhibited'/inhibited before conditioning, n = 48; 'Fear'/activated after conditioning, n = 49; 'Fear inhibited'/inhibited after conditioning, n = 60; 'Extinction'/activated after extinction, n = 86; 'Extinction inhibited'/inhibited after extinction, n = 43; 'Activated stable', n = 63; 'Inhibited stable', n = 105). **I**, Average traces of CS– clusters shown in H.

Average traces in B, G and I are mean with s.e.m.; violin plots in C show distribution of all data points, Tukey box-and-whisker plots in show median values, 25th and 75th percentiles, and min to max whiskers with exception of outliers, dots indicate the mean. *$p$ < 0.05, **$p$ < 0.01. Additional details of statistical analyses are provided in Supplementary Table 1.
