## Supplementary Table 1 for "Heterogeneous plasticity of amygdala interneurons in associative learning and extinction"

### Supplementary Table 1: Summary of all statistical analyses for data presented in main and supplementary figures.

Main figures

| Figure | Panel | Sample size | Statistical test | Individual comparisons |
|---|---|---|---|---|
| 2 | C | N = 9 mice | Friedman test<br>$\chi2 = 11.53$, $p = 0.0016$ | Dunn's multiple comparisons<br>CS+ vs. CS–, $p > 0.9999$<br>CS+ vs. US, $p = 0.0065$<br>CS– vs. US, $p = 0.0286$ |
| 2 | E | N = 9 mice | Friedman test<br>$\chi2 = 24.33$, $p < 0.0001$ | Dunn's multiple comparisons<br>CS+/US vs. CS–/US, $p > 0.9999$<br>CS+/US vs. CS+/CS–, $p = 0.1057$<br>CS+/US vs. CS+/CS–/US, $p = 0.0004$<br>CS–/US vs. CS+/CS–, $p = 0.0635$<br>CS–/US vs. CS+/CS–/US, $p = 0.0002$<br>CS+/CS– vs. CS+/CS–/US, $p = 0.6021$ |
| 2 | F | n = 519 cells | Chi-Square test<br>$\chi2(2) = 6.3912$, $p = 0.0409$ | Chi-Square with Bonferroni correction<br>(Activated vs. Inhibited)<br>CS+, $p < 0.0001$<br>CS–, $p = 0.2027$<br>US, $p = 0.9177$ |
| 3 | E | n = 519 cells | Friedman test<br>$\chi2 = 1.598$, $p = 0.8091$ | – |
| 3 | F | n = 519 cells | Friedman test<br>$\chi2 = 2.402$, $p = 0.6623$ | – |
| 3 | H | N = 9 mice | Friedman test<br>$\chi2 = 30.41$, $p < 0.0001$ | Dunn's multiple comparisons<br>Activated stable vs. US off, $p = 0.5314$<br>Activated stable vs. Else, $p = 0.0676$<br>Activated down vs. Activated up, $p = 0.0327$<br>Activated down vs. US off, $p = 0.0023$<br>Activated down vs. Else, $p < 0.0001$<br>Inhibited stable vs. Else, $p = 0.1854$<br>Inhibited up vs. US off, $p = 0.4609$<br>Inhibited up vs. Else, $p = 0.0566$<br>All other comparisons, $p > 0.9999$ |
| 4 | G | n = 519 cells | – | Paired Wilcoxon test with Bonferroni correction<br>(CS+ vs. CS–)<br>CS 1, $p = 1$<br>CS 2, $p = 1$<br>CS 3, $p = 1$<br>CS 4, $p = 0.158$<br>CS 5, $p = 0.0021$ |
| 4 | H | CS+ n = 297 cells<br>CS– n = 312 cells | Chi-Square test<br>$\chi2(6) = 117.19$, $p < 0.0001$ | Chi-Square with Bonferroni correction (CS+ vs. CS–)<br>Up, $p = 0.3987$<br>Down inhibited, $p = 0.2214$<br>Down activated, $p < 0.0001$<br>Stable, $p < 0.0001$<br>First activated, $p = 1$<br>First inhibited, $p = 1$<br>Else, $p = 0.0594$ |
| 5 | G | n = 519 cells | Before conditioning<br>Chi-Square test<br>$\chi2(2) = 2.4021$, $p = 0.3009$ | – |
| | | | After conditioning<br>Chi-Square test<br>$\chi2(2) = 0.1111$, $p = 0.9460$ | – |
| | | | After extinction<br>Chi-Square test<br>$\chi2(2) = 4.8741$, $p = 0.0874$ | – |

| | | | | |
|---|---|---|---|---|
| 5 | H | CS+ n = 365 cells<br>CS– n = 357 cells | Chi-Square test<br>$\chi2(7) = 105.84$, $p < 0.0001$ | Chi-Square with Bonferroni correction (CS+ vs. CS–)<br>Before conditioning activated, $p = 0.0275$<br>Before conditioning inhibited, $p = 1$<br>After conditioning activated, $p = 0.0016$<br>After conditioning inhibited, $p = 0.2213$<br>After extinction activated, $p = 0.0074$<br>After extinction inhibited, $p = 0.5894$<br>Stable inhibited, $p = 1$<br>Stable activated, $p < 0.0001$ |
| 5 | I | N = 9 mice | Friedman test<br>$\chi2 = 6.299$, $p = 0.3905$ | – |
| 6 | D | N = 9 mice | Friedman test<br>$\chi2 = 5$, $p = 0.172$ | – |
| 6 | E | N = 9 mice | CS+<br>Friedman test<br>$\chi2 = 13.9$, $p = 0.0030$ | Dunn's multiple comparisons<br>Pairing 2 vs. Pairing 3, $p = 0.0115$<br>Pairing 2 vs. Pairing 4, $p = 0.0209$<br>Pairing 2 vs. Pairing 5, $p = 0.0115$<br>Pairing 3 vs. Pairing 4, $p > 0.9999$<br>Pairing 3 vs. Pairing 5, $p > 0.9999$<br>Pairing 4 vs. Pairing 5, $p > 0.9999$ |
| | | | CS–<br>Friedman test<br>$\chi2 = 14.2$, $p = 0.0026$ | Dunn's multiple comparisons<br>Pairing 2 vs. Pairing 3, $p = 0.0370$<br>Pairing 2 vs. Pairing 4, $p = 0.0115$<br>Pairing 2 vs. Pairing 5, $p = 0.0061$<br>Pairing 3 vs. Pairing 4, $p > 0.9999$<br>Pairing 3 vs. Pairing 5, $p > 0.9999$<br>Pairing 4 vs. Pairing 5, $p > 0.9999$ |
| 6 | F | N = 9 mice | Friedman test<br>$\chi2 = 13.9$, $p = 0.0304$ | Dunn's multiple comparisons<br>All vs. Up, $p = 0.4309$<br>All vs. Stable, $p > 0.9999$<br>All vs. Down inhibited, $p > 0.9999$<br>All vs. First activated, $p > 0.9999$<br>All vs. First inhibited, $p > 0.9999$<br>All vs. Else, $p > 0.9999$ |
| 6 | G | N = 9 mice | Friedman test<br>$\chi2 = 36.0$, $p < 0.0001$ | Dunn's multiple comparisons<br>All vs. Activated stable, $p = 0.0272$<br>All vs. Activated down, $p = 0.2699$<br>All vs. Activated up, $p = 0.0754$<br>All vs. Inhibited stable, $p > 0.9999$<br>All vs. Inhibited up, $p > 0.9999$<br>All vs. US off, $p = 0.9508$<br>All vs. Else, $p > 0.9999$ |
| 6 | H | N = 9 mice | CS+<br>Friedman test<br>$\chi2 = 2.89$, $p = 0.2781$ | – |
| | | | CS–<br>Friedman test<br>$\chi2 = 14.2$, $p < 0.0001$ | Dunn's multiple comparisons<br>Conditioning vs. After conditioning, $p = 0.0005$<br>Conditioning vs. After extinction, $p = 0.1780$<br>After conditioning vs. After extinction, $p = 0.1780$ |
| 7 | C | SST n = 114 cells<br>VIP n = 101 cells | – | Mann-Whitney test with Bonferroni correction<br>(SST vs. VIP)<br>US 1, $p < 0.0001$<br>US 2, $p < 0.0001$<br>US 3, $p < 0.0001$<br>US 4, $p = 0.685$<br>US 5, $p = 0.0005$ |
| 7 | F | SST n = 114 cells<br>VIP n = 101 cells | – | Mann-Whitney test with Bonferroni correction<br>(SST vs. VIP)<br>CS+ 1, $p < 0.0001$<br>CS+ 2, $p = 1$<br>CS+ 3, $p < 0.0001$<br>CS+ 4, $p = 0.0287$<br>CS+ 5, $p = 0.0003$ |

| | | | | |
|---|---|---|---|---|
| 7 | G | SST n = 114 cells<br>VIP n = 101 cells | CS+<br>Chi-Square test<br>$\chi 2(2) = 30.885$, $p < 0.0001$ | Chi-Square with Bonferroni correction (SST vs. VIP)<br>Activated, $p < 0.0001$<br>Inhibited, $p = 0.0004$<br>No response, $p = 0.2852$ |
| | | | CS–<br>Chi-Square test<br>$\chi 2(2) = 2.5014$, $p = 0.2863$ | – |
| | | | US<br>Chi-Square test<br>$\chi 2(2) = 16.663$, $p = 0.0002$ | Chi-Square with Bonferroni correction (SST vs. VIP)<br>Activated, $p = 0.0028$<br>Inhibited, $p = 0.0007$<br>No response, $p = 1$ |
| 7 | H | SST n = 89 cells<br>VIP n = 80 cells | Chi-Square test<br>$\chi 2(6) = 28.635$, $p < 0.0001$ | Chi-Square with Bonferroni correction (SST vs. VIP)<br>Activated stable, $p = 0.0001$<br>Activated down, $p = 1$<br>Activated up, $p = 0.6272$<br>Inhibited stable, $p = 0.0978$<br>Inhibited up, $p = 0.7564$<br>US off, $p = 1$<br>Else, $p = 1$ |
| 7 | I | SST n = 63 cells<br>VIP n = 68 cells | Chi-Square test<br>$\chi 2(5) = 28.155$, $p < 0.0001$ | Chi-Square with Bonferroni correction (SST vs. VIP)<br>Up, $p = 0.3152$<br>Down inhibited, $p = 0.7037$<br>Stable activated, $p = 0.0046$<br>First activated, $p = 1$<br>First inhibited, $p = 0.0418$<br>Else, $p = 0.1501$ |
| 7 | J | SST n = 74 cells<br>VIP n = 55 cells | Chi-Square test<br>$\chi 2(5) = 9.5089$, $p = 0.0904$ | – |
| 8 | C | SST n = 114 cells<br>VIP n = 152 cells | – | Mann-Whitney test with Bonferroni correction (SST vs. VIP)<br>Before conditioning, $p = 0.0154$<br>After conditioning, $p = 0.205$<br>After extinction, $p = 0.0879$ |
| 8 | D | SST n = 114 cells<br>VIP n = 152 cells | Before conditioning<br>Chi-Square test<br>$\chi 2(2) = 9.7873$, $p = 0.0075$ | Chi-Square with Bonferroni correction (SST vs. VIP)<br>Activated, $p = 1$<br>Inhibited, $p = 0.0098$<br>No response, $p = 0.3386$ |
| | | | After conditioning<br>Chi-Square test<br>$\chi 2(2) = 6.5609$, $p = 0.0376$ | Chi-Square with Bonferroni correction (SST vs. VIP)<br>Activated, $p = 1$<br>Inhibited, $p = 0.0496$<br>No response, $p = 0.4349$ |
| | | | After extinction<br>Chi-Square test<br>$\chi 2(2) =23.938$, $p < 0.0001$ | Chi-Square with Bonferroni correction (SST vs. VIP)<br>Activated, $p = 0.0005$<br>Inhibited, $p = 0.1987$<br>No response, $p < 0.0001$ |
| 8 | E | SST n = 92 cells<br>VIP n = 96 cells | Chi-Square test<br>$\chi 2(7) = 13.065$, $p = 0.0705$ | – |

Supplementary Figures

| Figure | Panel | Sample size | Statistical test | Post-hoc test |
|---|---|---|---|---|
| S2 | B | N = 9 mice | Friedman test<br>$\chi2 = 11.26$, $p = 0.0465$ | Dunn's multiple comparisons<br>Down inhibited vs. Stable, $p = 0.1762$<br>Stable vs. First inhibited, $p = 0.6573$<br>Stable vs. Else, $p = 0.0836$<br>All other comparisons, $p > 0.9999$ |
| S2 | C | N = 9 mice | Friedman test<br>$\chi2 = 7.216$, $p = 0.2050$ | – |
| S4 | E | CS+ n = 165 cells<br>CS– n = 185 cells | Chi-Square test<br>$\chi2(1) = 0.5135$, $p = 0.4736$ | – |
| S4 | J | 12 kHz n = 202 cells<br>6 kHz n = 148 cells | Chi-Square test<br>$\chi2(1) = 0.5361$, $p = 0.4641$ | – |
| S3 | C | n = 519 cells | – | Paired Wilcoxon test with Bonferroni correction<br>(CS+ vs. CS–)<br>Before conditioning, $p = 0.169$<br>After conditioning, $p = 1$<br>After extinction, $p = 1$ |
| S3 | D | N = 9 mice | Friedman test<br>$\chi2 = 25.22$, $p = 0.0007$ | Dunn's multiple comparisons<br>Tone activated vs. Tone inhibited, $p = 0.6648$<br>Tone activated vs. Fear, $p = 0.0494$<br>Tone activated vs. Fear inhibited, $p = 0.0072$<br>Tone activated vs. Extinction, $p = 0.0212$<br>Tone activated vs. Extinction inhibited, $p = 0.0022$<br>Tone activated vs. Inhibited stable, $p = 0.5151$<br>Fear inhibited vs. Activated stable, $p = 0.5151$<br>Extinction inhibited vs. Activated stable, $p = 0.2279$<br>All other comparisons, $p > 0.9999$ |
| S5 | C | N = 9 mice | Friedman test<br>$\chi2 = 1.933$, $p = 0.5864$ | – |
| S8 | G | SST n = 114 cells<br>VIP n = 101 cells | – | Mann-Whitney test with Bonferroni correction<br>(SST vs. VIP)<br>CS– 1, $p = 1$<br>CS– 2, $p = 1$<br>CS– 3, $p = 1$<br>CS– 4, $p = 1$<br>CS– 5, $p = 1$ |
| S9 | C | SST n = 114 cells<br>VIP n = 152 cells | – | Mann-Whitney test with Bonferroni correction<br>(SST vs. VIP)<br>Before conditioning, $p = 0.375$<br>After conditioning, $p = 0.0324$<br>After extinction, $p = 1$ |
| S9 | D | SST n = 114 cells<br>VIP n = 152 cells | Before conditioning<br>Chi-Square test<br>$\chi2(2) = 0.6568$, $p = 0.7201$ | – |
| | | | After conditioning<br>Chi-Square test<br>$\chi2(2) = 4.9771$, $p = 0.0830$ | – |
| | | | After extinction<br>Chi-Square test<br>$\chi2(2) = 5.1876$, $p = 0.0747$ | – |
| S9 | E | SST n = 78 cells<br>VIP n = 79 cells | Chi-Square test<br>$\chi2(7) = 20.415$, $p = 0.0047$ | Chi-Square with Bonferroni correction (SST vs. VIP)<br>Before conditioning activated, $p = 1$<br>Before conditioning inhibited, $p = 1$<br>After conditioning activated, $p = 1$<br>After conditioning inhibited, $p = 1$<br>After extinction activated, $p = 0.0594$<br>After extinction inhibited, $p = 1$<br>Stable inhibited, $p = 1$<br>Stable activated, $p = 0.0249$ |
